# Topoisomerase IIβ targets DNA crossovers formed between distant homologous sites to modulate chromatin structure and gene expression

**DOI:** 10.1101/484956

**Authors:** Mary Miyaji, Ryohei Furuta, Osamu Hosoya, Kuniaki Sano, Norikazu Hara, Ryozo Kuwano, Jiyoung Kang, Masaru Tateno, Kimiko M. Tsutsui, Ken Tsutsui

**Affiliations:** Graduate School of Medicine, Dentistry and Pharmaceutical Sciences, Okayama University, Okayama, Japan; Department of Molecular Genetics, Bioresource Science Branch, Center for Bioresources, Brain Research Institute, Niigata University, Niigata, Japan; Asahigawaso Research Institute, Asahigawaso Medical-Welfare Center, Okayama, Japan; Graduate School of Life Science, University of Hyogo, Kamigori, Hyogo, Japan; Institute of Human Complexity and Systems Science, System Science Center for Brain and Cognition, Yonsei University, Seoul, Republic of Korea

**Keywords:** Topoisomerase II, Gene regulation, DNA crossover, Homologous pairing, Repetitive element, Chromatin structure

## Abstract

**Background:** Type II DNA topoisomerases (topo II) flip the spatial positions of two DNA duplexes, called G- and T-segments, by a cleavage-passage-resealing mechanism. In living cells, these DNA segments can be placed far from each other on the same chromosome. However, no direct evidence for this to occur has been described so far due to lack of proper methodology.

**Results:** The beta isoform of topo II (topo IIβ) is essential for transcriptional regulation of genes expressed in the final stage of neuronal differentiation. To elucidate the enzyme’s role in the process, here we devise a genome-wide mapping technique for topo IIβ target sites that can measure the genomic distance between G- and T-segments. It became clear that the enzyme operates in two distinctive modes, termed proximal strand passage (PSP) and distal strand passage (DSP). PSP sites are concentrated around transcription start sites, whereas DSP sites are heavily clustered in small number of hotspots. While PSP represent the conventional topo II targets that remove local torsional stresses, DSP sites have not been described previously. Most remarkably, DSP is driven by the pairing between homologous sequences or repeats located in a large distance. A model-building approach suggested that the DSP sites are intertwined or knotted and topo IIβ is engaged in unknotting reaction that leads to chromatin decondensation and gene regulation.

**Conclusions:** When combined with categorized gene expression analysis, the model-based prediction of DSP sites reveals that DSP is one of the key factors for topo IIβ-dependency of neuronal gene regulation.

## Background

Several lines of evidence demonstrated that the beta isoform of type II DNA topoisomerase (topo IIβ), which is reviewed recently [1, 2], is essential for transcriptional regulation of genes expressed in the last stage of neuronal development [3–5]. However, detailed mechanism for this process remains largely unknown. Locating the enzyme’s action sites on the genome would be a logical strategy to elucidate the role of topo IIβ. Type II DNA topoisomerases (topo II) catalyze the interconversion of the spatial positions of two DNA duplexes by a cleavage-passage-resealing mechanism. To discriminate these duplexes, the cleaved (gapped) strand and the other strand that is transferred through the gap are called G-segment and T-segment, respectively [6–8]. Topo II can be cross-linked to the G-segment right at the site of action by treating living cells with etoposide, a topo II-specific ‘poison’-type inhibitor. By taking advantage of this property of the enzyme and immuno-selecting the bound DNA, its action sites on the genome have been mapped in several studies [9–12]. Since the cleaved strand is covalently bound to the enzyme at the 5’ end, this procedure is similar to chromatin immunoprecipitation (ChIP) but essentially different in that no chemical cross-linker is used. Instead, cross-linking is based on the arrested enzymatic reaction intermediate. The procedure thus detects DNA sites directly involved in the reaction and not simply associated with the enzyme.

We have successfully used this type of strategy, termed “etoposide-mediated topoisomerase immunoprecipitation” or eTIP, for mapping of topo IIβ action sites (toposites) in selected genomic regions [11]. In that study we adopted oligonucleotide-tiling arrays to map the bound DNA fragments. In the present study we now extend this technology to a genome-wide scale by massive direct sequencing on NGS (named ‘eTIP-seq’). As a result, mapping resolution was improved significantly and, most importantly, repetitive DNA sequences became a reasonable subject of analysis. Another unique feature of the method is that the immunoprecipitated complex contains not only the covalently linked G-segment but also the T-segment associated with the complex non-covalently, which is released by high-salt wash and analyzed separately afterwards. This provides additional useful information unattainable from conventional mapping techniques.

One of the cellular events where topo II becomes essential is the disentanglement of intertwined chromatid DNAs generated at the final stage of cell division [8, 13]. In vertebrates, topo IIα is exclusively responsible for this reaction, namely decatenation. As for topo IIβ, it is believed to relax torsional stresses accumulated locally from various DNA transactions such as transcription [14]. In this reaction called relaxation, the enzyme removes writhes of DNA axis, either positive or negative, by passage between nearby DNA segments. In principle, however, distance between G- and T-segments can be much larger, like hundreds of kilo-bases, although no supporting evidence for this to occur has been reported to date. To detect these distant segmental passage events catalyzed by topo IIβ, we introduced additional steps to eTIP-seq, which include ligation between G- and T-segments via an oligomer adaptor attached to their sheared ends.

Using these techniques, we show here that topo IIβ operates in two distinct modes termed proximal strand passage (PSP) and distal strand passage (DSP) depending on the distance between G- and T-segments. PSP and DSP differ significantly not only in mechanistic sense but also in physiological consequences. While PSP sites are concentrated around transcription start sites (TSS) and also distributes throughout the genome to contribute the relaxation of local torsional stresses, DSP sites are heavily clustered in relatively small number of regions (hotspots) on chromosome. Most remarkable finding was that DSP occurs at the DNA crossovers facilitated by pairing of homologous DNA segments located at a long distance. This is an entirely novel perspective on the cellular function of topo II enzyme, in that a ‘DNA-centric’ mechanism may govern the target selection. The present study suggests that PSP is not involved in transcriptional induction at the final stage of neuronal differentiation. Instead, we provide some evidence that one of the major roles of topo IIβ in genomic context is to regulate higher-order chromatin structure through DSP events, leading to activation of suppressed neuronal genes.

## Results

### Genome-wide identification of topo II**β** target sites by eTIP-seq

The experimental procedure for eTIP-seq analysis is outlined in Fig. 1. Precursor cells of cerebellar granule neurons (CGN) isolated from infant rats were allowed to differentiate *in vitro*. Using the same culture system we have shown that topo IIβ is essential for transcriptional induction of a group of genes involved in mature neuronal function [4, 11]. In the present study, cells were treated with etoposide at the second day in culture to trap topo IIβ on target DNA. After lysis of treated cells with detergent sarkosyl followed by DNA fragmentation, the topo IIβ-DNA complex was recovered on magnetic beads coated with specific antibody to topo IIβ as described previously for eTIP procedure [11]. The topo IIβ-DNA complexes captured under these conditions are consisted of multiple forms as illustrated in the box of Fig. 1. They represent three major aspects: i) G- and T-segments can be either contiguous or segregated. ii) The covalent linkage between G-segment and the enzyme is partially reversed in the absence of etoposide. iii) DNA fragments interacting non-covalently with the enzyme remain bound within the complex.

**Fig. 1.**
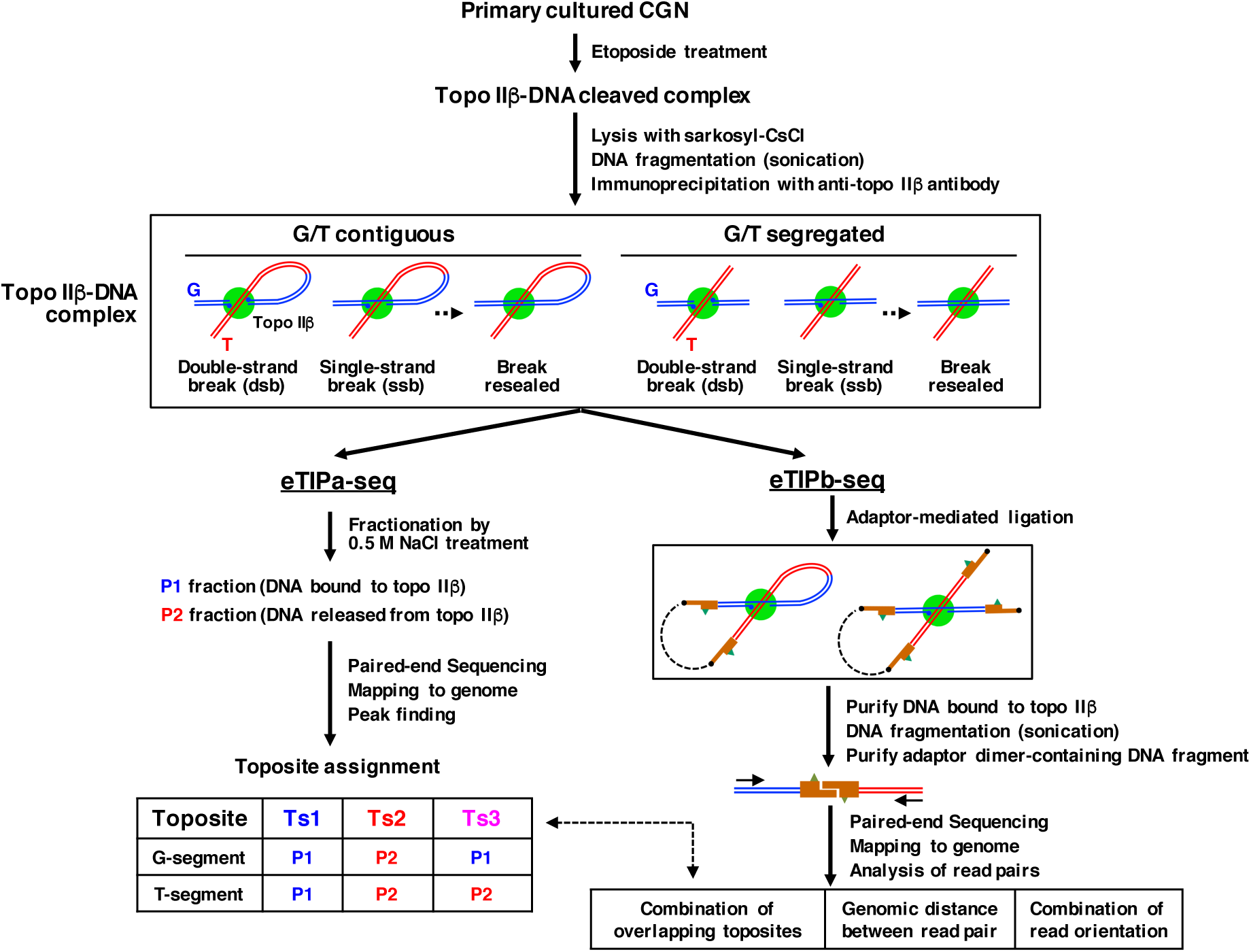
Overview of the eTIP-seq, a mapping technique used in the study. The topo IIβ-DNA complex illustrated in the box was isolated by eTIP procedure as described previously [11]. DNA in the topo IIβ-DNA complex captured on magnetic beads was then processed in either ways: fractionation by 0.5 M NaCl treatment (eTIPa-seq) or adaptor-mediated ligation (eTIPb-seq). Experimental results, data processing and toposite assignments for eTIPa-seq are summarized in Fig. S1a-d. Theoretical reasoning for the generation of three categories of toposites is shown in Fig. S2 by presenting all possible combinations of DNA fragments bound to topo IIβ. The eTIPb-seq procedure is outlined in Fig. S4a. See ‘Supplementary’ for detailed explanation on multiple forms of topo IIβ-DNA complex depicted in the box.

The procedure hereafter branches into two routes. In the left route labeled eTIPa-seq, the complex bound-DNA on magnetic beads was fractionated into P1 and P2 fractions and sequenced separately on NGS to categorize the topo IIβ target sites termed ‘toposites’. The eTIPb-seq on the right branch determines the G- and T-segments bound to the same topo IIβ molecule by paired-end sequencing of chimeric fragments generated by the adaptor-mediated ligation. To eliminate noise, resulting chimeras were selected by checking whether or not their ends overlap with toposites that are detected in eTIPa-seq.

Supporting evidence for the entire process of eTIP-seq is described in detail in the supplementary file (Supplementary) under the subheading of “Notes on methodology”.

### Three classes of topo IIβ target sites (toposites) are identified

We classified toposites into Ts1, Ts2, and Ts3 by eTIPa-seq (Fig. 1 and Fig. S1). Distribution of toposites in a large scale, a whole chromosome, reveals a strong correlation between toposites and local GC content: Ts1 being located in GC-rich area whereas Ts2 resides in relatively AT-rich area (Fig. 2a). The toposite density plot clearly shows that Ts2 and Ts3 clusters are embedded between Ts1-dense regions in a mutually exclusive manner. Assembly of Ts2/Ts3 in relatively small number of clusters is a feature that distinguishes from Ts1 site. Although Ts2 and Ts3 clusters appear to occupy similar positions in this scale, further analysis demonstrated that Ts2 and Ts3 are distinct toposites with unique characters. As shown later, these clusters constitute hotspots of topo IIβ genomic targets that are likely to be involved in some regulatory processes. Positions of Ts2/Ts3 clusters are tabulated in Table S2. Relative numbers of toposites differ significantly, Ts1 being most prominent (Fig. 2b).

**Fig. 2.**
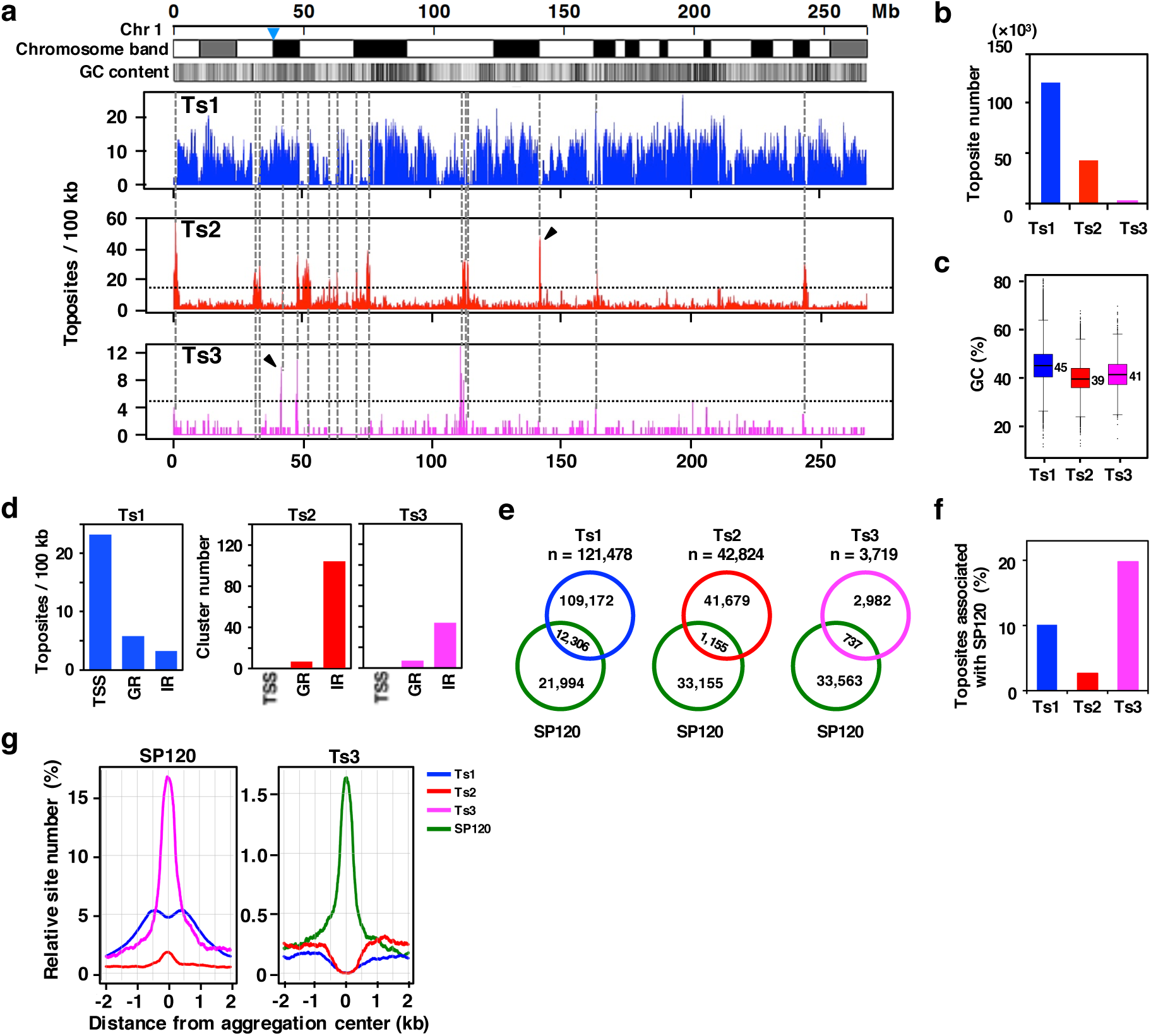
Genomic distribution of toposites and SP120 sites. **a** Density distribution of toposites on entire chromosome 1 as calculated by a sliding window algorithm (100 kb window, 10 kb step). Horizontal dotted lines show thresholds (Ts2: 15, Ts3: 5) for high-density peaks that are marked by vertical dotted lines. The position of centromere (p-q boundary) is indicated by nabla symbol and the arrowheads designate peaks that clearly distinguish Ts2 and Ts3. **b** Numbers of toposites detected. **c** A box plot of the GC content of toposites. Numbers indicate median values. **d** Toposite distribution among the three genomic regions: TSS zone, genic region (GR), and intergenic region (IR). TSS zone is excluded from GR and IR. Ts1 sites are expressed by number of sites per 100 kb, whereas number of clusters is plotted in Ts2 and Ts3 toposites. Positional data for toposite clusters are given in Table S2. **e** Overlapping situation of toposites with SP120 sites (n=34,300) as determined by ChIP-seq. Site numbers are shown. **f** Percentages of toposites overlapping with SP120 site. **g** Spatial relationships between SP120 site and toposites shown by aggregation plots. Centered sites are indicated on top.

GC contents of toposites are consistent with the above observation. Ts1 sites are significantly GC-rich as compared to Ts2/Ts3 sites (Fig. 2c). To investigate functional relevance, locations of these toposites were then looked at with respect to characteristic regions: transcription start site (TSS zone), genic region (GR), and intergenic region (IR) (Fig. 2d). Only protein-coding genes are taken into account using the “exRefSeq” which is derived from the rat reference sequence [11]. TSS zone (+/- 2 kb region encompassing the transcription start site) was included in the analysis as a functional region associated with genes. TSS zone normally overlaps with transcription control region containing promoter and binding sites for transcription factors. Ts1 was highly concentrated in gene-associated regions especially in the TSS zone. In contrast, Ts2/Ts3 sites, as viewed from the number of clusters, were enriched in intergenic regions.

Ts3 toposites frequently overlap with SP120 sites, which is a most remarkable finding (Fig. S1e, bottom track). SP120 (hnRNP U/SAF-A/SP120) is a multifunctional nuclear protein that binds both RNA and DNA [15–18]. A recent report shows that the protein binds chromatin-associated RNA and is involved in the organization of chromatin structure in a transcription-dependent manner [19]. Our previous study demonstrated that SP120 is a partner protein of topo IIβ that activates and stabilizes the enzyme through RNA-dependent association [20]. We have also shown that randomly cloned genomic targets of topo IIβ are frequently enriched with SP120 binding sites, suggesting that these proteins bind DNA within close distance [21].

Overlapping situation between toposites and SP120 sites is illustrated by Venn diagrams (Fig. 2e). The SP120 site overlaps with Ts1 site considerably but overlaps very little with Ts2 site. Also, a significant portion (20%) of Ts3 site overlaps with SP120 site (Fig. 2f), suggesting that at least two distinct genomic sites are involved in the interaction between SP120 and topo IIβ. Next, we examined the spatial relationship between toposites and SP120 sites by aggregation plot [22, 23], in which relative site numbers at varying positions around the aggregation center were plotted against the distance from the centered sites (Fig. 2g). Most significantly, SP120 and Ts3 toposites are aggregated at the same position indicating their close association. Ts1 sites, however, appears to aggregate about 400-500 bp apart from the SP120 site, forming two symmetrical peaks alongside the center. This characteristic pattern reflects the presence of a loop structure at Ts1 sites, which will be demonstrated by eTIPb-seq analysis (see Fig. 4h).

**Fig. 3.**
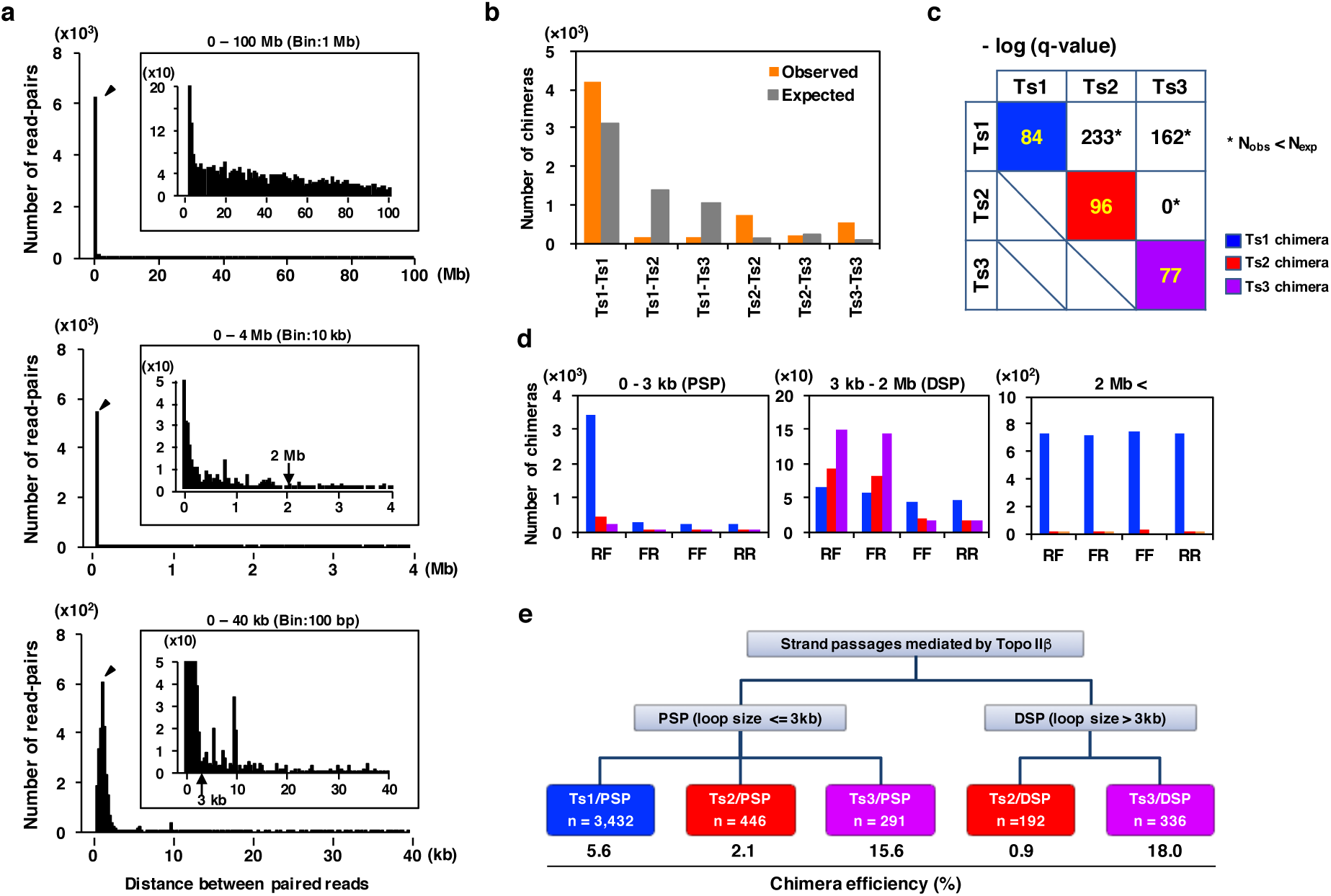
Analysis of intra-chromosomal chimeras identified by eTIP-seq. **a** Size distribution histograms of chimeras. Number of read-pairs (chimera) is plotted against distance between paired reads (chimera length) in 3 levels of magnifications (0-100 Mb, 0-4 Mb, and 0-40 kb). **b** Toposite combinations of chimeras. Expected numbers are calculated on the basis of experimental frequencies of each toposites at the end of chimeras. **c** Statistical analysis of toposite combinations. Chi-squared test was conducted between expected and observed frequencies of each combination. Resulting p-values were corrected by Benjamini-Hochberg method to obtain q-values. Shown in the matrix are minus-log transformed q-values. Asterisks indicate the combination where observed number is smaller than expected one. **d** Abundance of chimeras in 3 ranges of chimera length (0-3 kb, 3 kb-2 Mb, and 2 Mb<). Ts1, Ts2, and Ts3 chimeras were classified into 4 categories depending on the read combination (RF, FR, FF, and RR), which is designated by read orientations (forward=F, reverse=R) arranged in the order of upstream to downstream. Blue, Ts1 chimera; red, Ts2 chimera; magenta, Ts3 chimera. **e** Summary of classified chimeras. Abundance of each chimera group (n) is shown in color-coded boxes on the bottom. Chimera efficiency indicates the percentage of toposites associated with chimera ends against total number of each toposite.

**Fig. 4.**
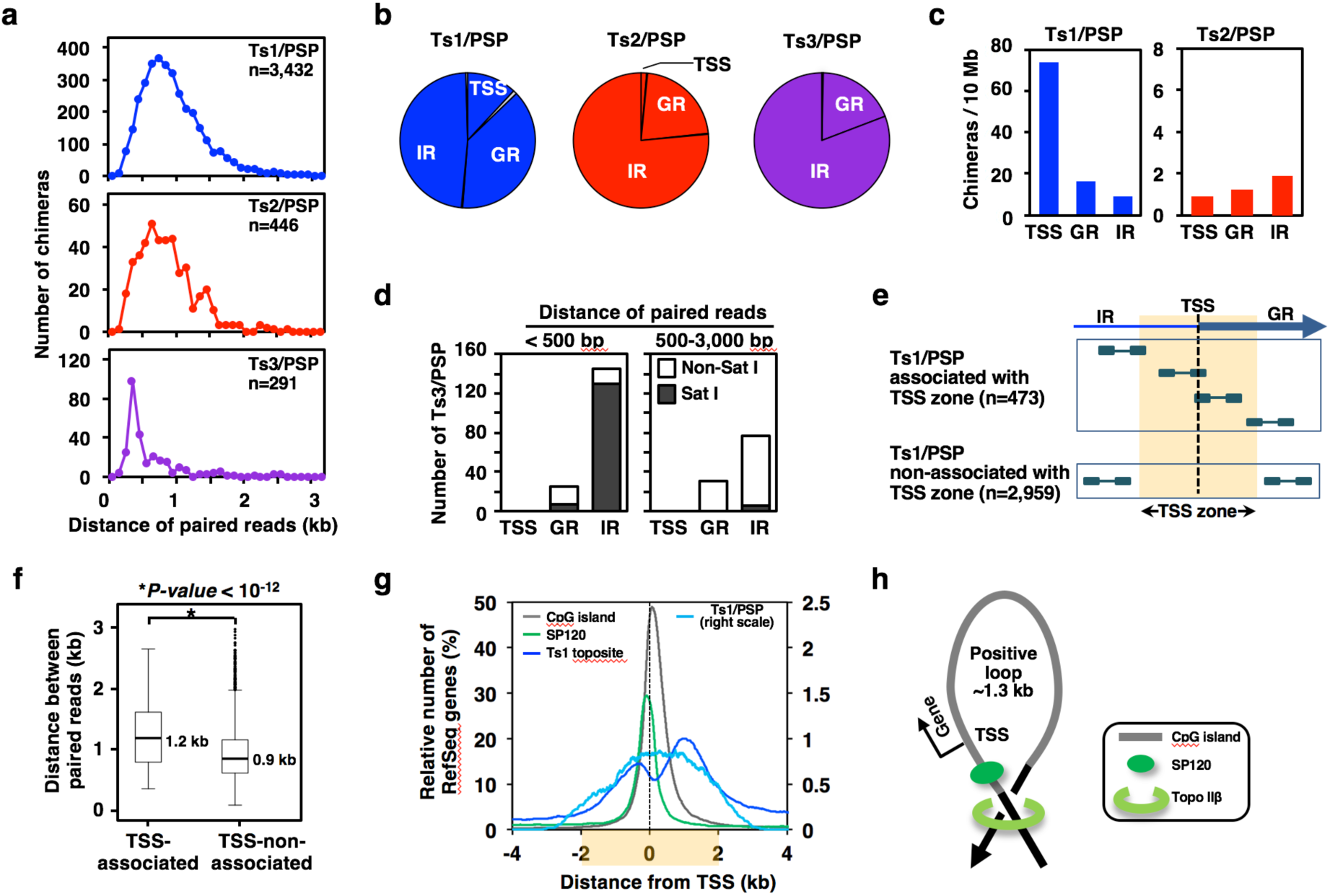
Analysis of PSP chimeras. **a** Size distribution curves. **b** Location of PSP chimeras in different genomic regions as defined in Fig. 2d. **c** Density plot of PSP chimera distribution. **d** Difference in the genomic distribution of Ts3/PSP in two groups of chimera size. Short Ts3/PSP is accounted for by satellite I repeat in intergenic region (filled area). **e** Conditions for dividing Ts1/PSP into two groups on the basis of association with TSS zone (TSS +/- 2 kb). Sequence reads on chimera ends are designated by rectangles. **f** A box plot for length distribution of Ts1/PSP. Horizontal bars indicate median length. TSS-associated Ts1/PSP is significantly longer than non-associated ones. **g** Aggregation plot. CpG island, SP120 site, Ts1 toposite, and Ts1/PSP chimera overlapping with TSS zone (shadowed) were aggregated. Annotation data for RefSeq genes and CpG island were downloaded from the UCSC genome browser site. The right scale applies only to Ts1/PSP. **h** A model for the relationship between topo IIβ action site and SP120 binding site in the vicinity of TSS. This is based on the aggregation data shown in ‘g’. A plectonemic loop with positive turn (PSP loop) is most likely to be formed in the region.

### The eTIP-seq opens a new perspective on the reaction mode of topo II enzyme

To acquire positional information between DNA fragments associated with topo IIβ, we devised a new procedure that includes a ligation step between G- and T-segments via an oligomer adaptor attached to their sheared ends. We analyzed the resulting library of paired-read sequences that are referred to as ‘chimeras’. With respect to intra-chromosomal chimeras, various features like chimera length, combination of nearby toposites, read orientations, and genomic locations provide useful information. It should be noted that ‘chimera length’ indicates the map distance between paired reads and not the physical length of chimeric DNA fragments. We first plotted histograms for distribution of chimera length (Fig. 3a). Most evidently, a large proportion of chimera is rather short (51% are shorter than 3 kb; marked by arrowhead). Longer chimeras of certain length, however, do exist and appear as peaks between 3 kb and 2 Mb but chimeras longer than 2 Mb rarely form peaks and the number levels off to a background level. Therefore, only chimeras shorter than 2 Mb will be analyzed hereafter.

In theory, chimeras can be classified into 6 groups as to the combination of nearby toposites on both ends. From the number of toposites contributing to chimeras, expected number of chimera for randomly chosen toposite pairs of every combination can be calculated. Observed numbers of chimera were then plotted along with expected ones (Fig. 3b). Significance of toposite combinations were determined statistically by comparison of ‘observed’ versus ‘expected’ and the significance levels are summarized (Fig. 3c). The result clearly indicates that only homologous combinations (Ts1-Ts1, Ts2-Ts2, and Ts3-Ts3) are highly favored whereas heterologous ones are avoided conversely. The presence of heterologous chimeras is probably caused by occasional errors in peak-calling and toposite assignments. Based on these data, only chimeras of homologous combinations will be taken into account and are referred to simply as Ts1, Ts2, and Ts3 chimeras, respectively.

We next examined what sort of information would be available from orientations of read pairs on chimera ends (Fig. 3d). The situation changed dramatically depending on the range of chimera length. In chimeras shorter than 3 kb (PSP, proximal strand passage), RF is by far the most dominant orientation for Ts1, Ts2, and Ts3 chimeras, suggesting that sonication-resistant loop structure (PSP loop) is still present at the ligation step (Fig. S4d). In chimeras between 3 kb and 2 Mb (DSP, distal strand passage), however, the RF dominance in Ts1 chimera is no longer apparent, whereas FR becomes as dominant as RF in Ts2 and Ts3 chimeras. This appears to indicate that the loop structure is disrupted in this range of length and FR is created as an additional combination for ligation (Fig. S4d). Since *cis*-ligation is more feasible to occur than *trans*-ligation as mentioned in the eTIPb-seq procedure (Supplementary), the high ratio of (RF+FR)/(FF+RR) in Ts2 and Ts3 chimeras implies that plectonemic loop-type crossover is favored over toroidal loop-type crossover in these chimeras. As chimera length exceeds 2 Mb, all classes of chimera show no preference in the read orientation, indicating that long chimeras are nonspecific in nature and very likely to be originated from the ligation events between neighboring IP complexes fixed on beads, thus termed inter-complex chimera (see Fig. S5a). In summary, the Ts1 toposite involves only PSP events whereas Ts2/Ts3 toposites are linked to both PSP and DSP events. The chimera categories and numbers identified are summarized in Fig. 3e. To avoid any confusion on chimera identity, chimeras will be referred hereafter with suffix (/PSP or /DSP) attached to the toposite name. Also shown in the figure is ‘chimera efficiency’, which roughly indicates what percentage of toposites is converted into chimeras. Ts3-involving chimeras showed extremely high efficiency compared to others.

### The three classes of PSP sites show distinct location

Ts1/PSP is most abundant in number compared to Ts2/PSP and Ts3/PSP chimeras (Fig. 3e and Fig. 4a). Comparison of length distribution showed that median length for Ts1/PSP and Ts2/PSP is around 700-800 bp, whereas Ts3/PSP is significantly shorter and peaks at around 400 bp (Fig. 4a). The similarity of length distribution suggests that Ts1/PSP and Ts2/PSP originate from similar complexes with looped DNA, although in Ts2/PSP the G-segment breakage might be reversed as in Ts2 toposites (Fig. S2b).

Location of PSP chimeras in the classified genomic regions was examined to assess their functional relevance (Fig. 4b). Ts1/PSP occurs frequently in TSS zone and genic regions (GR), implying that topo IIβ is involved in PSP events that are associated with transcription. Ts2/PSP and Ts3/PSP, however, are enriched in intergenic regions (IR) and rarely associated with TSS zone. The difference in regional distribution between Ts1/PSP and Ts2/PSP is demonstrated more clearly by the density plots (Fig. 4c). Abundance of Ts2/PSP correlates negatively with regional GC contents (median values for TSS, GR, IR are 48.7%, 45.4%, and 44.7%, respectively). The GC content of Ts2/PSP was significantly lower than that of Ts1/PSP in all genomic regions, corroborating the notion that the single-strand breaks in G-segment are indeed labile if the involved region is AT-rich (41-43% GC).

As the length distribution curve of Ts3/PSP has a sharp peak around 400 bp (Fig. 4a), regional partition of Ts3/PSP was looked at in two groups of chimera size (shorter or longer than 500 bp) (Fig. 4d). We found that the 400 bp-peak is mainly composed of the satellite I (Sat I) sequence enriched in intergenic regions, whereas Sat I is only a minor component of the long chimera group. Sat I is a rat-specific repetitive element [24]. Multiple copies of Sat I repeats (unit length∼370 bp) aligned in tandem constitute the centromeric region spanning megabases on most chromosomes in the rat [25]. The repeat unit is composed of 4 subrepeats that are homologous to each other (Fig. S7a). Browser view of SP120/Ts3 overlapping sites revealed that they frequently coincide with satellite I repeats in the RepeatMasker track. MEME analysis of Ts3 toposites located multiple motifs with high confidence (low E-value) within the Sat I repeat (Fig. S7b). The Sat I family is composed of a vast variety of sequences in rat genome that are diverged from a prototype sequence in the process of evolution. Their similarity against the prototype is expressed by the Smith-Waterman score (SW score). It is believed that Sat I sequences with high SW score constitute a significant part of centromeric DNA. When frequency distribution of SW score for Sat I repeats were plotted, they appeared to be segregated into two groups: with high SW score (>2000) and low SW score (<1000) (Fig. S7c). If the Sat I sites overlapped with Ts3 toposites were looked at, more than 80% of them are classified in the high SW score group (Fig. S7d). Thus, it is suggested that the centromeric Sat I sequence is one of the major targets of topo IIβ /SP120 complex and frequently detected as Ts3/PSP sites.

The remarkable enrichment of Ts1/PSP in TSS zone suggests that it is involved in the control of transcriptional initiation. We divided Ts1/PSP into two groups: associated and non-associated with TSS zone (Fig. 4e). Analysis of length distribution showed that TSS-associated Ts1/PSPs are significantly longer than the other group (Fig. 4f). With respect to the sequences around TSS (+/- 4 kb) extracted from rat RefSeq genes, Ts1/PSP and other features were plotted in an aggregation plot (Fig. 4g). Peak regions covering each genomic position were counted-up and expressed as relative numbers of RefSeq genes. SP120 and CpG island peak around TSS, whereas Ts1 toposite forms twin peaks that locate ∼1,300 bp apart harboring the other peaks. These results are consistent with the model shown in Fig. 4h. As depicted in the figure, the left peak of Ts1 toposite very likely to correspond to the cleavage site of G-segment bound by topo IIβ in action. The enzyme can also approach the duplex crossover from the other side, which should create the right peak. Taken together it is strongly suggested that topo IIβ complexed with SP120 recognizes a right-handed crossover formed between the positions of ∼300 bp upstream TSS and ∼1 kb downstream into the gene. In average genes, therefore, topo IIβ /SP120 acts on positively supercoiled loop of ∼1,300 bp formed around TSS whose size matches well with the chimera size of ∼1,200 bp (Fig. 4f). After strand passage, the loop is converted to negative loop, which may facilitate the initiation step of transcription [26]. Ts1/PSPs that are not associated with TSS but enriched in genic region may represent topo IIβ involved in the relaxation of positive supercoils generated by ongoing transcription [27]. In this case the loop also contains interaction sites for SP120 but the loop size is smaller (800-900 bp).

### DSP sites are clustered to form a small number of ‘hotspots’

Now we investigate DSP chimeras that reflect the topo IIβ-mediated interaction between distant genomic loci. These chimeras are categorized into two groups, Ts3/DSP and Ts2/DSP, which link Ts3 and Ts2 toposites, respectively (Fig. 3e). Genomic distribution of DSP chimeras showed a great deal of clustering, which often coincided with toposite clusters of the same categories, Ts2 or Ts3. A list of these “chimera-rich regions”, termed DSP clusters, is given in Table S3. By comparing the position of these clusters on the genome browser, here we present a new concept termed ‘hotspot’ that integrates the toposite clusters and DSP clusters (Table S4). The hotspots, either Ts2- or Ts3-hotspot, are depicted in a karyogram together with toposite clusters and DSP clusters (Fig. 5a). With the exception of Chr19, hotspots occur at least once in most chromosomes, indicating that they can be regarded as chromosome landmarks for topo IIβ targets, which is significantly limited in number. The length distribution showed that Ts2/DSPs and their clusters are much longer than Ts3/DSPs and their clusters (Fig. 5b, c). With reference to gene density profile, the Ts2-hotspot usually resides in gene-poor regions, often adjacent to gene-rich region or “gene city”. Ts3-hotspots are fewer than Ts2-hotspots and contain several interesting loci of tandemly repeated gene clusters such as noncoding RNA gene clusters and clusters of MHC (major histocompatibility complex) genes. Browser views of representative loci are shown in Fig. 5d and Fig. S8.

**Fig. 5.**
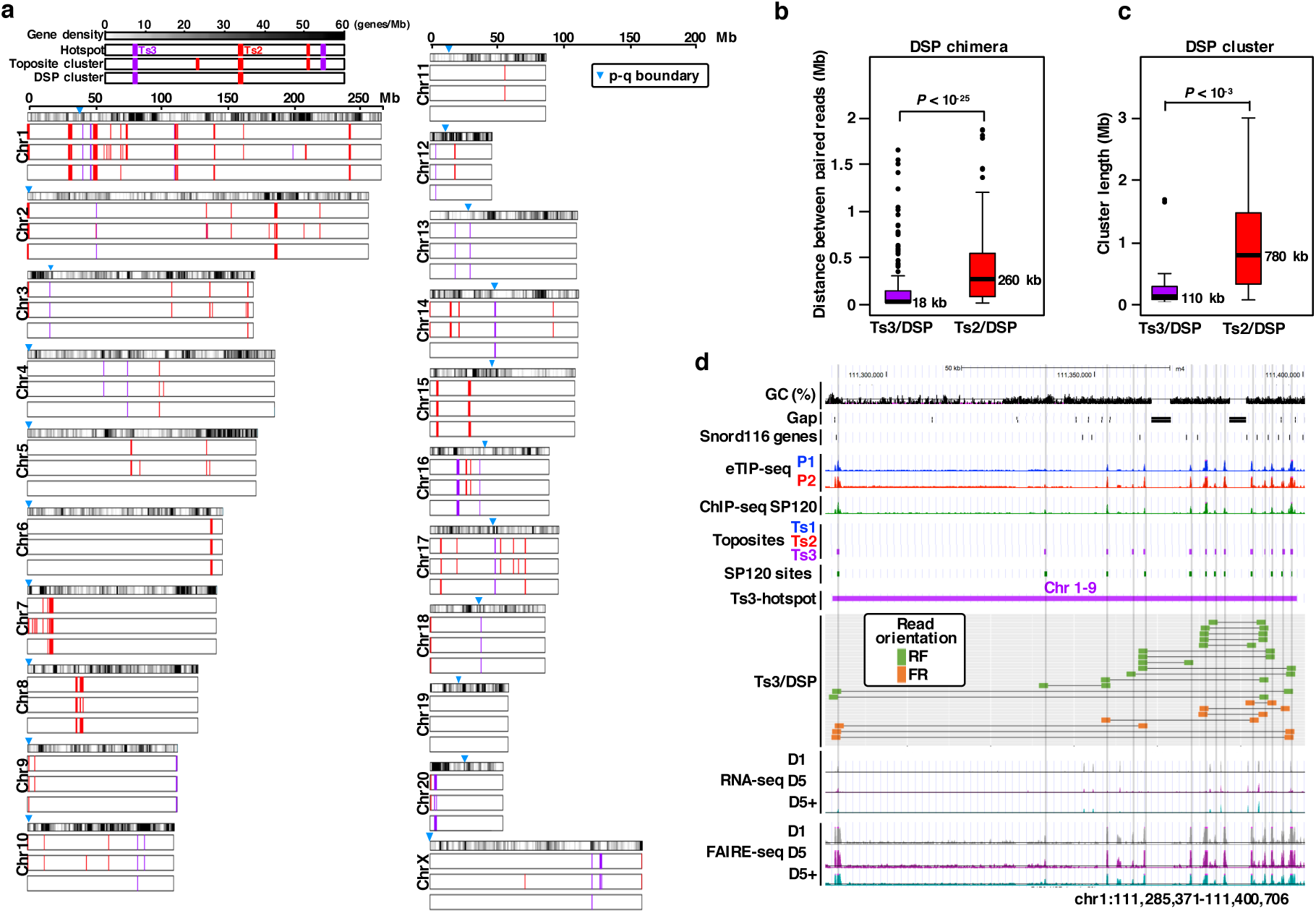
Analysis of DSP chimeras. **a** Karyogram of hotspots with reference to gene density, toposite cluster, and DSP cluster. Gene density (number of genes per 1 Mb) is depicted by grayscale gradient. Tracks are arranged in the same order as in the legend shown at the top-left corner. Ts2 (red) and Ts3 (magenta) sites are shown by bars in the same track. The boundary between cytobands p and q marked by nabla symbol indicates approximate position of centromere. The karyogram plots were generated by R package ggbio. **b** A box plot of length distribution of DSP chimeras. Horizontal bars indicate median length. Ts2/DSP is more than tenfold longer than Ts3/DSP. **c** A box plot of length distribution of DSP clusters. Ts2/DSP cluster is longer than Ts3/DSP cluster. **d** A browser view of the Snord 116 locus, a typical Ts3 hotspot. Entire region of the Ts3-hotspot (Chr1-9) is shown. Ts3 toposites are marked by vertical lines to indicate the accordance with other peaks. mRNA-seq and FAIRE-seq assays were done at culture day 1 (D1), day 5 (D5), and day 5 in the presence of topo II inhibitor ICRF-193 (D5+).

The Ts3-hotspot Chr1-9 shown in Fig. 5d constitutes a cluster of noncoding RNA gene (snoRNA), Snord116, whose transcripts are involved in modification of rRNA and are expressed prevalently in the brain [28]. This locus overlaps with the Prader-Willi syndrome (PWS) region (corresponds to human chromosome 15q11-13), which is expressed only paternally and if deleted the subject may suffer developmental delay and mental retardation [29]. Read positions of all the Ts3/DSP chimeras mapped in this region coincided well with Ts3 toposites and SP120 sites. Their read orientations were either RF or FR. It is worth noting that in accord with these sites high FAIRE-seq peaks were detected, indicating that these Ts3/DSP sites stay in an open chromatin conformation all the time in culture. The coincidence of these features is shared by all other Ts3-hotspots.

As for the possible link to genes, Ts2-hotspots are sometimes associated with clusters of olfactory/vomeronasal receptor genes and sperm-related genes (*Smok* and *Spetex*). Interestingly, intergenic Ts2-hotspots closely coincide with the distribution of segmental duplication and SNP rates (for functional implications of Ts2-hotspots, see Discussion).

### Sequences around the DSP chimera ends are homologous each other

In all the featured Ts3-hotspots (Fig. S8), repetitive or homologous sequences were present around the chimera ends. To generalize this observation, sequence similarities and read orientations were examined for all Ts3/DSP and Ts2/DSP chimera ends. We do not know the exact position of the crossover targeted by topo IIβ but can assume that it is relatively close to the chimera ends (Fig. S4d). Therefore, from each chimera ends we cut out 2 kb-genomic sequences that contain the reads in the center and measured the homology between the two. Since homologous sequences can reside either on plus (Watson) or on minus (Crick) strand, similarity alignments were made between both strands and the resulting pair of Smith-Waterman scores were compared (Table S5 and Fig. 6a). As expected, in all DSP chimeras (both Ts3 and Ts2) either one of the two alignments showed much higher score compared to the other, indicating that chimera ends are indeed highly homologous. Since homologous segments of duplex DNA aligned in parallel direction are likely to associate each other [30, 31], this may facilitate the formation of crossover for topo IIβ recognition. Both directions of repeats (direct and inverted) were present and, most importantly, the combination of read orientation precisely correlated with the directionality of repeats: RF/FR for direct repeat and FF/RR for inverted repeat (Table S5). This rule enabled us to propose a model that explains how interactions of homologous DNA segments located distantly can create crossover points for the action of topo IIβ (Fig. 9).

**Fig. 6.**
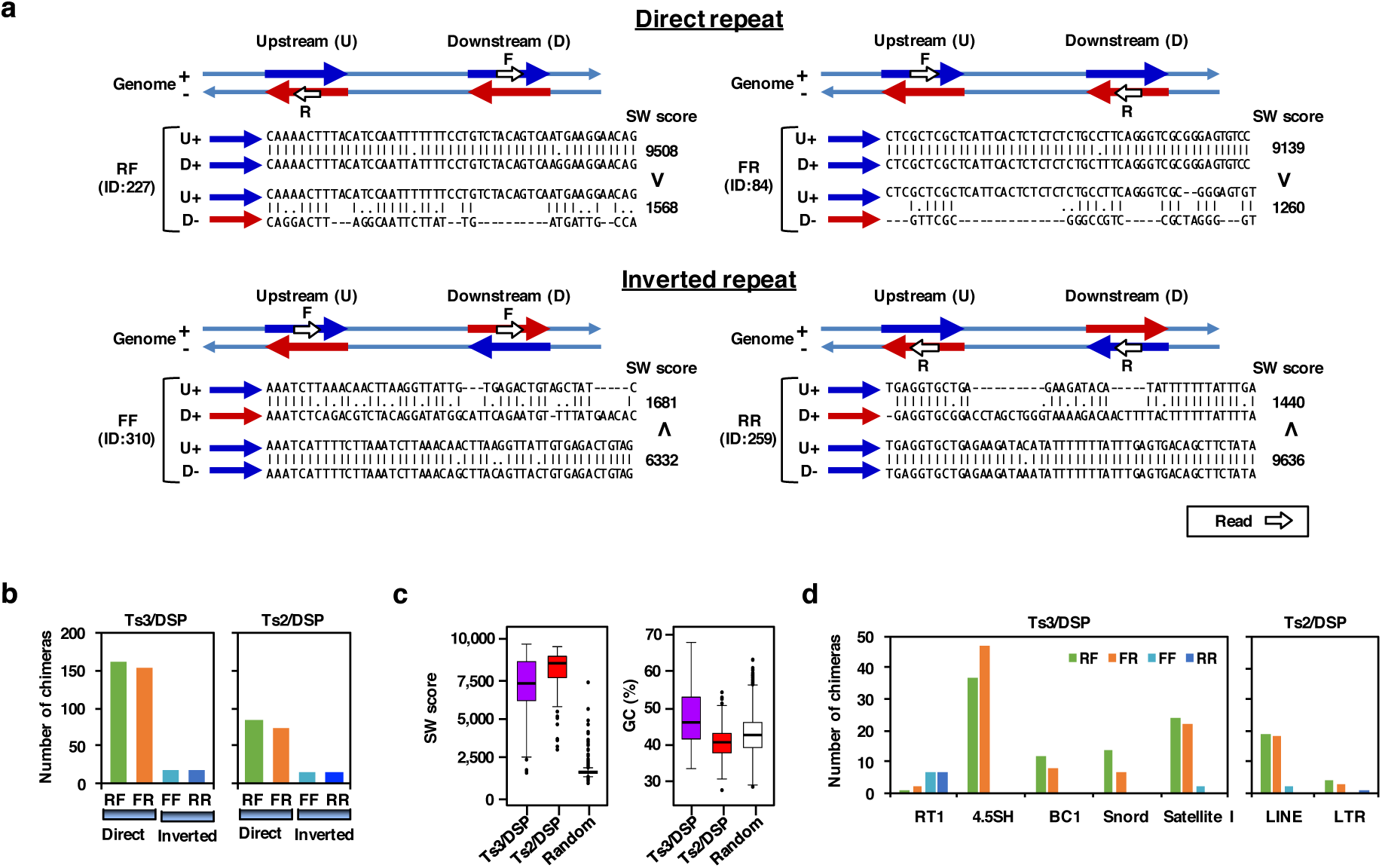
Read pairs associated with DSP chimera ends are homologous each other. **a** Comparison of the combination between read orientations and sequence homology at chimera ends. To exemplify the relationships leading to a general rule, four Ts3/DSPs with different read orientations (RF, FR, FF, RR) were chosen arbitrary from Table S5 (indicated by their ID in the figure). Two kilobases of genomic segments that contain sequence reads in the center were subjected to homology search by EMBOSS Water program. Homologous segments are referred to as direct or inverted ‘repeat’ depending on their orientation. Pairwise alignments between upstream and downstream segments were done in two ways: plus strand vs. plus strand or plus strand vs. minus strand (U+ vs. D+ or U+ vs. D-). Aligned sequences in the figure are 50-base region adjacent to the upstream reads. Resulting homology scores (SW score) and inequality signs are indicated on the right. **b** Differential abundance of DSP chimeras with direct repeats and inverted repeats at the homologous ends. **c** Box plots for distribution of SW score and GC content of DSP chimeras. Random pairing was made between randomly selected 2 kb-segments from Chr 1 that contained less than 20% gap coverage and are located within 2 Mb. **d** Matching of DSP chimera ends with annotated loci and RepeatMasker entries. RT1 stands for the rat major histocompatibility complex (MHC) genes. 4.5SH, BC1, and Snord are noncoding RNA genes.

Judging from the repeat orientation, homologous pairing of direct repeats occurs much more frequently than that of inverted repeats for both DSP chimeras (Fig. 6b). Although end-pairs of both DSP chimeras showed high homology scores over randomly matched pairs (Fig. 6c, left), Ts2/DSP scored significantly higher than Ts3/DSP. If higher homologous score reflects tighter association between the chimera ends, this may suggest that Ts2/DSP requires more stable homologous pairing to compensate its lower chimera efficiency (Fig. 3e). Considering the unstable nature of the DNA-topo IIβ complex at Ts2 sites (Fig. S2b), the difference in GC content between these DSP sites is also consistent with the above notion (Fig. 6c, right). A subset of Ts2/DSP chimera ends overlaps with known RepeatMasker entries, LINE and LTR (Fig. 6d). As described in the previous section, Ts3/DSP sites are enriched in the loci for MHC genes (RT1), noncoding RNA genes (4.5SH RNA, BC1 RNA, and Snord116) in addition to satellite I. The strong bias toward RF/FR orientation indicates that the homologous direct repeats are preferred target of topo IIβ, which was confirmed by examining individual loci for Ts3/DSP (Fig. S8). The RT1 locus is an exception because FF/RR is a favored combination, which can be explained by the presence of long Ts3/DSP chimeras between the two RT1 gene clusters that originated from a duplication/inverted insertion event, occurred during rodent evolution (Fig. S9). Major repeats like LINE or SINE were clearly minor constituents in Ts3/DSP. In contrast, however, LINE was the prevalent repeat associated with Ts2/DSP. It should be noted that a large proportion of chimera ends, 35% for Ts3/DSP and 66% for Ts2/DSP, are not attributable to any known repeats although these end-pairs are also highly homologous each other. We have not pursued their identity any further but they appear to be composed of several sequence groups that tend to cluster in separate DSP hotspots.

Homologous sequences are likely to be involved also in inter-chromosomal DSPs. As described in Discussion, Sat I subrepeats on different chromosomes are linked by Ts3/inter chimeras with statistical significance (Fig. S16). A recent report demonstrates that physical contacts between chromosomes are mediated by rDNA that are organized in tandem repeats of nearly identical copies [32]. The rDNA linkages are resolved in anaphase by topo II enzyme, suggesting that the linkages are topological intertwines. These observations are consistent with our model in that topo II is involved in formation-resolution of knotting between distant genomic regions (Fig. 9). However, DSP chimeras connecting rDNA repeats were not detected in our study since ribosomal repeating units are not currently annotated on the rat reference genome.

Taken together, there exists a certain genomic region where repetitive sequences are accumulated as DSP hotspots that serve as a platform for distant chromatin interactions mediated by topo IIβ.

### Topo IIβ-dependent alteration of nuclear structure during CGN differentiation

Analyses described hitherto are based on the data from eTIP-seq experiments that are performed at the day 2 of CGN (cerebellar granule neuron) culture, when differentiating cells are most active and abundant [4]. Hereafter, we will analyze the data sets obtained in three conditions: before differentiation at culture day 1 (D1), after differentiation at day 5 in culture (D5), and at day 5 cultured in the presence of topo II inhibitor (D5+). This type of experiment, as described in our previous study [11], is designed to identify differentiation-dependent changes (D1 vs. D5) and to determine whether or not topo IIβ is required in the process (D5 vs. D5+).

To assess changes in nuclear shape and chromatin state, cellular DNA was stained with Hoechst. When galleries of nuclear images from D1, D5, and D5+ were compared (Fig. 7a), the nuclear shape appeared more round and number of heterochromatic regions that are stained brightly decreased to leave a big blob in the center after 5 days in culture. However, this change does not occur when topo IIβ was suppressed by a specific inhibitor, ICRF-193 (D5+), suggesting that the enzyme is involved in the rearrangement of chromatin structure. This result was corroborated more objectively by a machine learning program called Wndchrm for image analysis [33]. The similarity matrix shows that the nuclear appearance is very different between D1 and D5 but it is more similar between D1 and D5+ (Fig. 7b). A 3D image analysis showed that nuclear volume enlarged by 20% during differentiation in a topo IIβ-dependent manner, which probably reflects the chromatin dispersion (Fig. 7c).

**Fig. 7.**
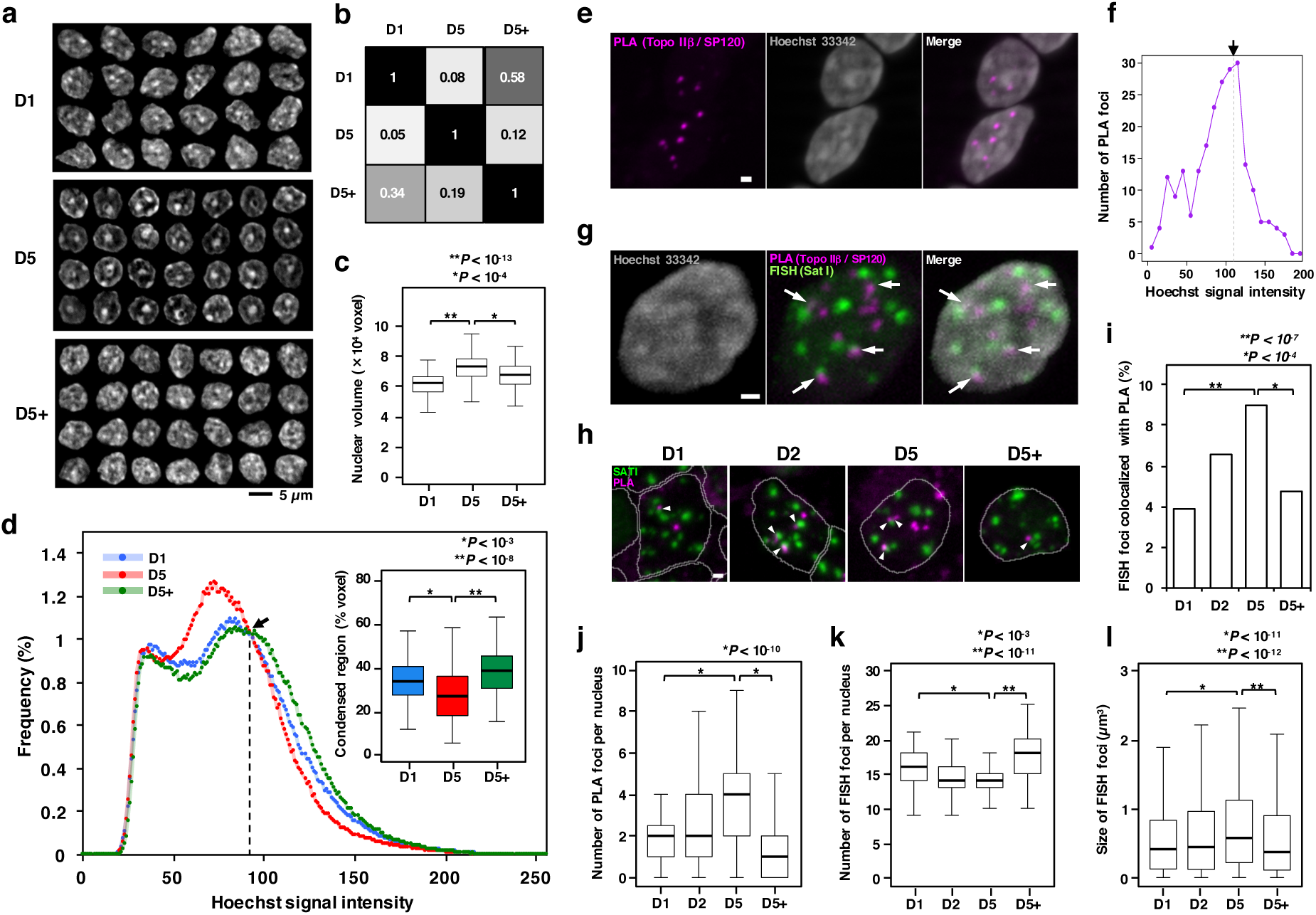
Morphological observations implicating the involvement of topo IIβ in the differentiation of CGN *in vitro*. **a** Changes of chromatin structure. Nuclear DNA in fixed cells was stained with Hoechst 33342 at the culture days indicated. Nuclear images (z-stack with maximal diameter) were collected and arranged in 3 panels. **b** Similarity analysis by Wndchrm. Eighty images each were used for learning. Similarity indexes (between 0 and 1) shown are average of 4 trials. **c** Measurement of nuclear volume. Horizontal bars in boxplot indicate median values and p-values are from Mann-Whitney U test. **d** Degree of chromatin condensation as estimated from DNA staining intensities. The isosbestic point marked by arrow indicates the border between condensed and dispersed chromatin regions. Percentages of condensed region (right side) were box-plotted (inset). **e** Visualization of topo IIβ-SP120 interacting loci by proximity ligation assay (PLA) using culture day 2 (D2) cells. Z-projection images are shown. Scale bar, 1 µm. **f** Number of PLA foci plotted against the brightness of DNA stain where PLA resides. Total number of PLA signals used was 225. Dotted line marked by arrow indicates the borderline between condensed and dispersed chromatin regions. **g** Simultaneous detection of PLA and FISH signals. CGN cells at D5 were used and Z-projection images are shown. Arrows indicate the signals that are in contact or colocalized. Scale bar, 1 µm. **h** Close positioning of PLA and FISH spots during CGN differentiation (arrowheads). Scale bar, 1 µm. **i** Numerical analysis of FISH/PLA colocalization. See Methods for details. P-values were obtained by chi-squared test with Holm’s correction for multiple comparison. **j** Numerical change of PLA foci during differentiation. **k** Numerical change of FISH foci (Sat I) during differentiation. **l** Volumetric change of FISH foci (Sat I) during differentiation.

To investigate the distribution of nuclear DNA more directly, spectrum of DNA staining intensity was measured (Fig. 7d). The graph represents the histogram of brightness (256 levels in grayscale) obtained from 100 nuclear images for each condition. The three curves had an isosbestic point at the brightness 92-95 (arrow), which was regarded as a boundary between bright (right side) and dark (left side) regions that correspond to condensed (heterochromatic) and dispersed (euchromatic) chromatin compartments, respectively. The ratio of heterochromatin was calculated from the total voxel number of condensed area and plotted (inset). The result indicates that overall chromatin state shifts toward decondensation in topo IIβ-dependent manner during differentiation.

Toposite distribution revealed that Ts1 and Ts3 sites often overlap with SP120 site, suggesting the presence of topo IIβ-SP120 complex (Fig. 2f). We tried to visualize these sites using a cytological technique called PLA (proximity ligation assay), which is designed to localize closely interacting proteins using specific antibodies followed by a signal amplification step involving rolling cycle amplification [34]. As shown in Fig. 7e, the topo IIβ/SP120 PLA signals appeared as relatively small number of demarcated spots, whereas immunostaining with individual antibodies gave much broader staining patterns [20, 35]. Considering the limited number of spots and the clustering tendency of Ts3 toposites, the result suggests that the visualized foci represent Ts3 clusters, most likely to be the Ts3-hotspots (Table S4). Judging from the image merged with DNA stain, these PLA signals reside close to condensed chromatin. This was confirmed by the DNA distribution analysis similar to Fig. 7d. PLA signals accumulate at the dispersed side of the condensed-dispersed chromatin boundary (arrow in Fig. 7f).

Three out of 21 Ts3-hotspots are Sat I clusters located in paracentromeric regions of chromosomes 13, 17 and 18 (Table S4). We thus looked at the spatial distance between FISH signals for Sat I probe and PLA signals (Fig. 7g). The FISH foci detected here represent centromeres, whose sequence information is not included in the standard reference genome. Although FISH and PLA signals frequently come close together, they rarely overlap exactly, suggesting that the PLA signal resides in pericentromeric region. Images from the time course analysis confirmed this observation (Fig. 7h). A 3D-colocalization analysis revealed that fraction of FISH foci colocalizing with PLA showed a significant positive correlation with the topo IIβ-dependent differentiation (Fig. 7i). Although the number of PLA foci increased with days in culture (Fig. 7j), number of FISH foci decreased slightly in topo IIβ-dependent manner (Fig. 7k). The increase in the volume of FISH foci may suggest that centromere structure undergoes a certain level of decondensation along with the global rearrangement of chromatin by topo IIβ (Fig. 7l). We thus speculate that the centromere structure is predisposed to disintegrate by topo IIβ/SP120 complex in terminally differentiating neurons.

We found that other Ts3-hotspots are also detected by PLA although their signal intensity is much weaker. For instance, significant colocalization between PLA and the Snord116 locus was observed (data not shown).

### Topo IIβ-dependent alteration of gene expression in terminally differentiating neurons

Now we investigate the expected correlation between genomic locations of toposites and gene expression. To achieve this, we first classified all genes with respect to expression patterns. Experimental protocol used here is illustrated in Fig. 8a (see Methods for details). RNA samples prepared from D1, D5, and D5+ cells were subjected to mRNA-seq analysis. Levels of mRNA expression for exRefSeq genes were deduced from FPKM values and tabulated in Table S6 together with a number of other features associated with the gene. Results were expressed by a two-dimensional logarithmic plot, in which the fold induction during differentiation (D5/D1, abscissa) was plotted against the fold susceptibility to topo II inhibitor (D5/D5+, ordinate) (Fig. 8b). The scattering pattern of gene dots roughly resembles a tilted ellipsoid. The graph area was divided into 9 parts by 4 borderlines passing through 1.5- or 0.66-fold points, which correspond to 9 distinct expression groups termed A1, A2, A3, B1, B2, B3, C1, C2, and C3. In addition to these, genes that are not expressed in any conditions were classified as group D. The largest gene group B2 represents the genes whose expression level is relatively constant with respect to both differentiation and topo II repression. Thus, the B2 group can be regarded as ‘house keeping genes’. Gene groups A1 and A2 are up-regulated during differentiation in topo IIβ-dependent or independent manner, respectively. In contrast, groups C3 and C2 are down-regulated in topo IIβ-dependent or independent manner, respectively. Hereafter, we analyze mainly these “diagonal groups” by ignoring the others that are relatively minor in abundance (Fig. 8c).

**Fig. 8.**
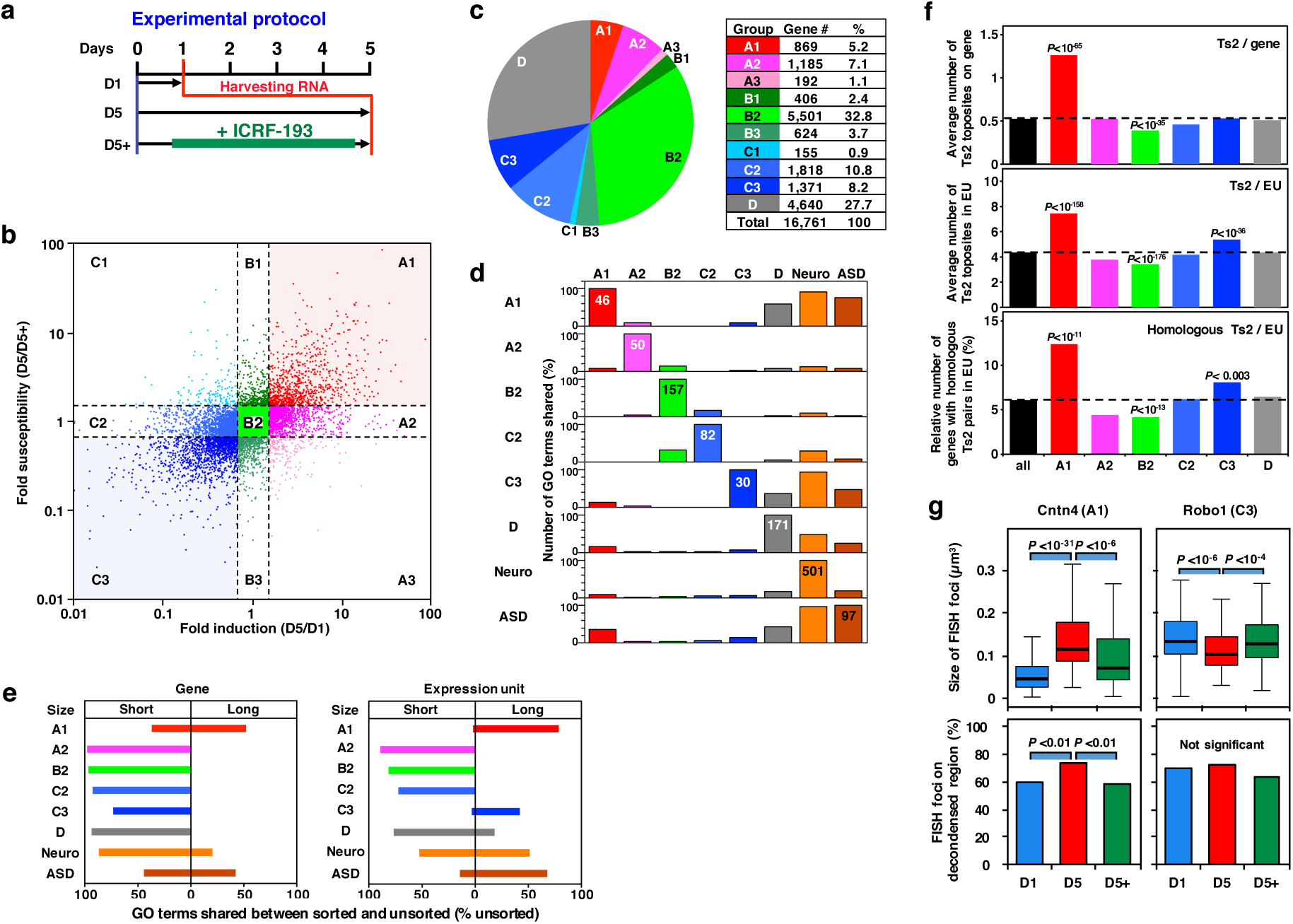
Analysis of categorized genes expressed in differentiating CGN. **a** Outline of the protocol used for the expression analysis in culture. RNA samples used for mRNA-seq analysis were prepared at the time points indicated. **b** Two-dimensional logarithmic plot of FPKM values obtained from mRNA-seq experiment. Fold-induction during differentiation (D5/D1) was plotted against fold-susceptibility to topo II inhibitor (D5/D5+). exRefSeq genes were divided into 9 groups by the dotted boarder lines that indicate 1.5- or 0.66-fold. **c** A pie graph and a breakdown list for the constituent expression groups. Group ‘D’ represents unexpressed genes. **d** Shared GO term analysis. Percentages of significant terms shared between other groups are plotted in the bar graph. Term numbers for hundred percent are shown in the bar. **e** Shared GO term analysis conducted with expression groups that are sorted into ‘long’ and ‘short’ subgroups. The threshold length was determined as in Fig. S11. Shared term numbers between sorted and unsorted were expressed as percentages of unsorted groups. **f** Correlation of Ts2 toposites to topo IIβ dependency. Ts2 toposites were counted either within the gene body (top) or within EU (middle). By pairwise homology search with EMBOSS Water algorithm, Ts2 pairs with high SW score (1000<) and identity (50%<) were identified. Relative number of genes with more than one homologous Ts2 pairs in EU was plotted for each expression group (bottom). P-values were calculated by Fisher’s exact test against the ungrouped (all). **g** Different behavior of A1 and C3 genes with respect to chromatin condensation as detected by FISH and DNA staining. Volumetric changes in FISH foci reflect expression levels (upper panels). Percentage of FISH foci residing in decondensed chromatin region was determined as in Fig. 7d and plotted (lower panels). P-values are from chi-squared test with Holm’s correction.

A1 is the most noteworthy group because its expression is increased only when topo IIβ is in action. Unexpectedly, there existed a gene group whose expression is repressed by topo IIβ during differentiation (C3). Gene ontology (GO) analysis revealed with statistical significance that A1 genes are involved in mature neural functions (ion channel, receptor, transporter, synapse, and adhesion proteins), whereas C3 genes are involved in earlier neural developments (developmental process, cell communication, cell motility, signal transduction, and cell differentiation) (Table S7). Functional difference between the gene groups can be easily captured by the list of associated GO terms ordered by statistical significance and sorted for each gene group (Fig. S10). To assess functional similarities between expression groups, we devised a “shared GO term analysis” (Fig. 8d). Although both A1 and C3 genes share a high percentage of GO terms with ‘neuronal’ genes, the functional contrast between these groups is very interesting. Since the CGN cells in culture are in the process of terminal differentiation, transition of neuronal phenotypes is expected to occur at this stage: from precursor phenotype (C3) to mature phenotype (A1). These results, therefore, are consistent with the notion that topo IIβ is a key molecule responsible for the coordinated reciprocal switching of the two gene groups. GO analysis provided additional significant findings on topo IIβ-independent gene groups, as well. A2 genes that are up-regulated during differentiation are enriched with GO terms related to mitochondrial function and energy metabolism, whereas down-regulated C2 genes are involved in cell cycle and mitosis (Fig. S10). This observation may reflect the shutoff of mitotic functions that are no longer required and the up-regulation of ATP synthesis to meet the mature neuronal activities.

Another remarkable finding from the shared GO term analysis was that A1 genes shared a high percentage of GO terms with ASD (autism spectrum disorder)-related genes as well as with neuronal genes. However, C3 group showed less similarity to ASD-related genes, probably reflecting that neuronal function of relatively late onset is impaired in ASD. Other gene groups showed little similarity to neuronal/ASD phenotypes. In our previous study, we have categorized gene expression patterns into similar groups and shown that about 20% of A1 genes are adjacent to long and AT-rich intergenic regions [11]. We specified these as “LA genes” and pointed out that a large proportion of ASD-related genes are LA genes. Our observation was confirmed later by a study showing that topo IIβ is involved in the transcription of long genes associated with ASD [36]. These studies suggested that the region of gene body combined with flanking intergenic regions, defined as ‘expression unit (EU)’ here, is likely to be an important contributing factor in terms of transcriptional control of A1 genes and possibly C3 genes. By contrasting the gene length and EU length, we performed a binomial clustering analysis, ‘long’ versus ‘short’, using thresholds determined by a ROC (Receiver Operating Characteristic) analysis (Fig. S11). The results showed that both A1 and C3 gene groups are enriched with long EU, suggesting that EU, not gene length, is relevant to the topo IIβ dependency of these genes.

This trend is manifested more clearly by the shared GO term analysis. Each gene group was divided into long and short subgroups and the relative numbers of significant GO terms that coincided with the term names of the undivided one were plotted in opposite direction (Fig. 8e). A dramatic shift toward ‘long’ side was observed in A1, C3, and ASD groups when EU size was considered instead of gene size. However, functional characteristics of other gene groups were represented by the short EU subgroup. These results strongly suggest that topo IIβ is exclusively involved in topological transactions on nerve-related genes with long EU. Physical evidence supporting this notion was obtained by counting the number of Ts2 toposites on gene or EU (Fig. 8f). Again, the number was significantly higher in A1 and C3 genes only when EU was considered (Fig. 8f, middle panel). Since the chimera efficiency is particularly low (only 0.9%) for Ts2/DSP (Fig. 3e), actual number of captured Ts2/DSP should be very little. To estimate the number of potential Ts2/DSP, we counted Ts2 toposite pairs with homologous sequences that reside within 2 Mb in EU domain. Genes with homologous Ts2 pairs were significantly enriched in A1 and C3 groups but depleted in B2 group (Fig. 8f, bottom panel). Taken together, topo IIβ-catalyzed distant segmental passages within EU would be a feasible mechanism for the observed topo IIβ-dependency of A1/C3 genes.

As demonstrated in Fig. 7d, topo IIβ is responsible for the alteration of global chromatin structure during differentiation, which can be described as decrease in condensed heterochromatic region or increase in dispersed euchromatic region. To investigate the behavior of genic chromatin in this nuclear context, we performed FISH analysis together with DNA staining (summarized in Fig. 8g). Two representative genes with long EU were selected from A1/C3 groups for FISH target. *Cntn4* (A1) encodes an axon-associated cell adhesion molecule involved in neuronal network formation and a candidate gene for ASD [37]. *Robo1* (C3) encodes an integral membrane protein that functions in axon guidance and neuronal precursor cell migration [38]. The volume of FISH signal, which enlarges when genic chromatin opens during transcription, can detect changes in the transcriptional state of long genes. Changing patterns of foci volume at D1, D5, D5+ were LHL (low, high, low) in *Cntn4* and HLH in *Robo1*, which is consistent with the mRNA levels detected by mRNA-seq (Table S6). Then, locations of FISH foci with respect to DNA signal intensity were determined to measure the local state of chromatin condensation where the target gene resides. As shown in lower panels of Fig. 7g, FISH signal for *Cntn4* migrated to more dispersed region at D5 in topo IIβ-dependent manner, indicating that the enzyme is required for the opening of chromatin structure to maintain the elevated transcription. As for *Robo1*, however, the structure of genic region chromatin appeared to be open already at D1 and stayed open at D5, suggesting that for the transcriptional suppression to occur the region must be open by continued action of topo IIβ.

## Discussion

### DSP is mediated by homologous pairing between remote genomic sites: a model

In the present study we demonstrated unequivocally for the first time that topo IIβ is engaged in strand passage events between distant genomic sites with sequence homology (Fig. 6). DNA crossovers targeted by the enzyme are likely to be triggered by association of homologous DNA segments. The homology-sensing can occur between chromatin segments, not necessarily between naked DNA, because oligo-nucleosomes with identical DNA sequence were shown to associate selectively in solution [39]. However, surprisingly little evidence has been published on the physical basis for the association between duplexes with homologous sequences despite the fact that it is an essential step for important biological processes such as meiotic pairing of homologous chromosomes. Only recently, some theoretical approaches suggested that interaction between the two duplexes of DNA that are aligned side-by-side is most stable if the pair has a certain sequence homology and the direction of alignment is parallel instead of anti-parallel [30, 40].

Here we propose a model that implies the presence of two distinct topo II-dependent processes: formation and disruption of higher order chromosomal structures (Fig. 9). Since major grooves of paired duplex face each other at the left-handed crossover [41], additional hydrogen bonds can be formed between identical base pairs in the paired DNA as suggested to occur in four-stranded DNA structure [30, 31]. Using computational methods, Mazur proposed a convincing model for the mutual recognition between two homologous duplex DNAs through direct complementary hydrogen bonding of major groove surfaces in parallel alignment [30]. At the contact area, a quadruplex structure is formed between the duplexes associated with left-handed crossover configuration. The quadruplex can extend three to four consecutive base pairs called ‘recognition unit’ that have to be spaced by at least several helical turns. Therefore, the pairing requires relatively long stretches of DNA, but only partial homology. Initial pairing is likely to occur in ‘paranemic’ mode (Fig.5 in [30]) to generate a right-handed crossover between the recognition units, which is a preferred target of topo II. After strand passage between the direct repeat (homologous pairs are on the same strand), number of recognition units are doubled and the pairing mode turns to intertwined mode introducing knots into the looped domain between the homologous pairs (Fig. 9, upper section). In contrast, when homologous pairs are on opposite strands (inverted repeat), negative supercoils, instead of knots, are produced within the looped domain (Fig. 9, lower section).

The paranemic association between homologous pairs should be quite unstable without the consecutive action of topo II. Although, in principle, the topo II enzyme involved here can be either topo IIα or topo IIβ, topo IIα is the feasible one since it is the major enzyme in cycling cells that is required for the mitotic chromosome condensation in G2/M phase [42]. We speculate that the knotting reaction between LINEs, an abundant repetitive element in mammals, could significantly contribute to the chromatin condensation along the chromosome axis. Roles of repetitive elements in organization of higher-order chromatin structures have been proposed [43].

**Fig. 9.**
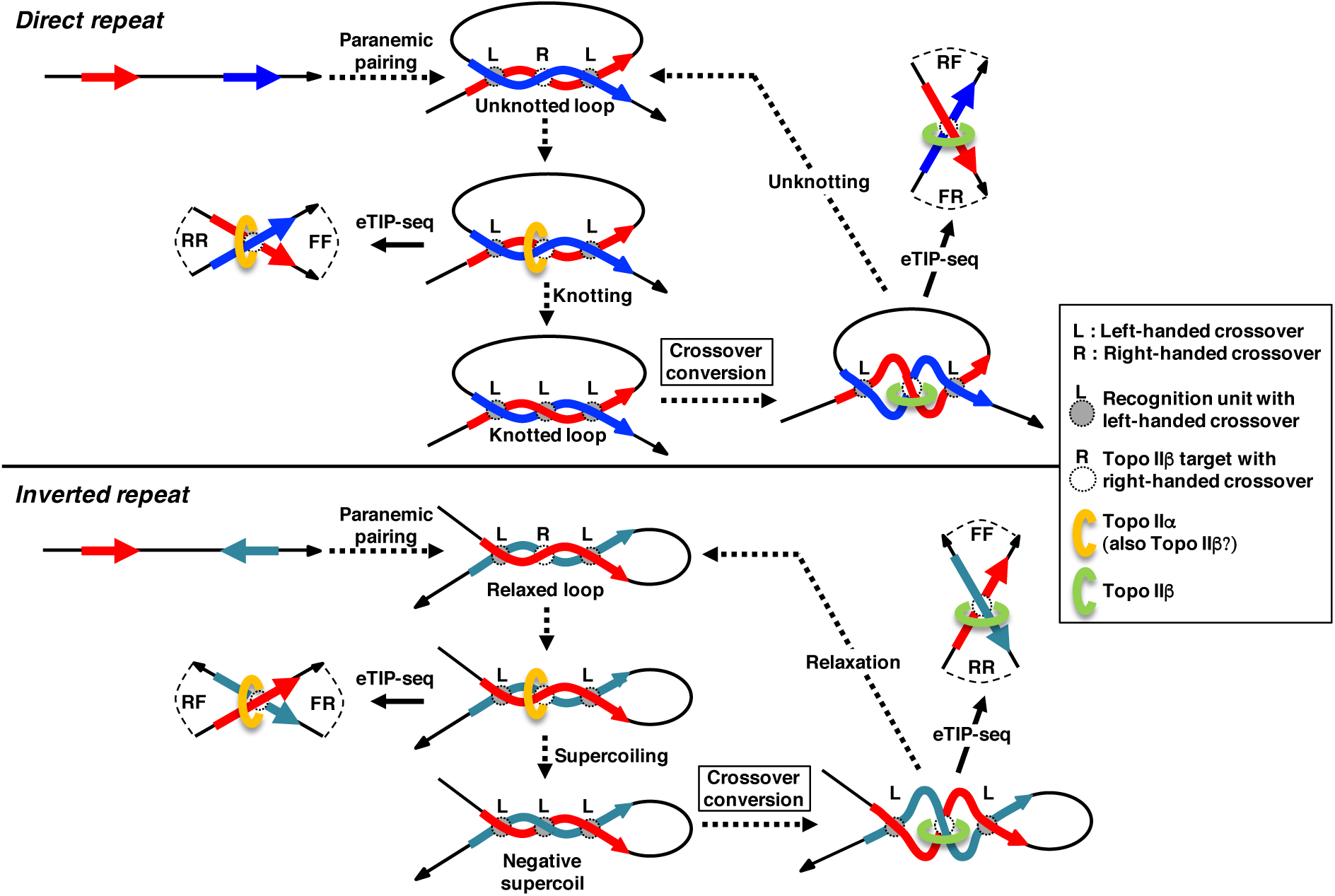
Homologous pairing between repetitive DNA segments at the DSP site: a model. Topo II action at these sites leads to different consequences depending on the repeat orientations, direct and inverted. Plus-strand DNA path from 5’ to 3’ direction is depicted by black arrows. Homologous segments are shown by red arrows (upstream) and blue arrows (downstream) on plus strand or by dark-green arrows on minus strand. Homologous pairing between duplex DNAs aligned in parallel starts by ‘paranemic’ mode and converted to intertwined mode after topo II action. Left-handed crossovers facilitate the interaction between major grooves to form ‘recognition unit’, which is a quadruplex structure required for stable pairing [30]. Since the pairing occurs only when the two DNA segments are aligned in parallel, direct and inverted repeats bring about very different results both in the loop configuration between paired repeats and in the topological structure generated after topo II action. Direct and inverted repeats result in knotted loops and negative supercoils, respectively. The ‘crossover conversion’, which is an energetically favored step, is a mandatory process for the reverse reaction (unknotting or relaxation) to occur. The eTIP-seq experiments performed in the present study produced dominant DSP chimeras with RF/FR read orientations that are originated from direct repeats. This suggests that in terminally differentiating CGN cells topo IIβ is almost exclusively involved in unknotting reactions (illustrated in the upper right).

The model predicts that at the time of eTIP-seq analysis (day 2 in culture) topo II is engaged in the removal of knots between direct repeats or supercoils between inverted repeats because the combination of read orientations detected in DSP chimeras was exclusively RF/FR for direct repeats and FF/RR for inverted repeats (Table S5). We can state definitively that topo IIβ is the enzyme involved in these reactions since β-specific antibody is used for eTIP-seq and topo IIα is not expressed in these cells. We emphasize that eTIP-seq can discriminate the directionality of topo II reaction, knotting or unknotting, by determining the read orientations on the chimera between direct repeats. The favored read orientations in DSP chimeras, RF/FR over FF/RR (Fig. 6b), indicate that direct repeats are the preferred target of topo IIβ probably because knots are detrimental to transcription [44, 45]. This is consistent with the potential DSP sites estimated by homology search. When abundance of homologous Ts2 toposite pairs in EU was calculated separately for direct repeats and inverted repeats, their enrichment in EU of A1 gene showed much higher significance for direct repeats than for inverted repeats (Fig. S12a). The reason why inverted repeats are only minor target of the enzyme would be that negative supercoils are constantly removed by the action of topoisomerase I. Also, negative supercoils are rather favorable for transcription and thus need not to be removed.

The left-handed crossover in the knot should be converted to more stable right-handed configuration by thermal wobbling that is recognizable by topo IIβ [41, 46]. It appears that the conversion of crossover handedness is a mandatory process because topo IIβ does not recognize left-handed crossovers. The crossover handedness at the DNA gate has been confirmed recently by crystallographic evidence [47]. There might be some protein factors involved in additional stabilization of DNA crossovers at DSP sites. Since about 68% of Ts3/DSP is overlapped with SP120 sites (Fig. S13a), the protein may serve as a crossover stabilizer at these sites. But it is not so at Ts2/DSP sites where SP120 is almost absent. Although the possibility that other protein is responsible for the crossover stability cannot be eliminated, we suggest that homologous pairing at Ts2/DSP sites can be conducted purely by ‘DNA-centric’ manner without any assistance from proteins.

This model applies to only DSPs because Ts1/PSP chimera showed very little homology between the chimera ends (less than 2% showed high homology scores). PSP crossovers are enriched in local PSP loops and thus homologous pairing would not be required to facilitate crossovers.

### Genome-wide prediction of homologous pairs for probing potential DSP sites

It should be possible to predict the position of homologous pairs by a genome-wide search with a sliding-window algorithm similar to the Harr plot [48]. Needless to say, the eTIP-seq read positions are close to but not right on the actual crossover point where topo IIβ binds, although these sites may not be apart from each other by more than ∼2 kb. We applied the DotMatcher analysis to calculate the homology score for 2 kb-segments within 2 Mb-bins. The resulting map represents the distribution of pHP (predicted homologous pairs), which can be regarded as potential crossover points for topo IIβ targets. As expected, a representative region from Chr1 shows that most of the Ts2 toposite clusters are enriched with pHP (Fig. S14). The reason why not all the pHP-enriched region coincide with Ts2 toposite clusters is probably that only uniquely mapped reads are adopted in the eTIP-seq procedure so that the reads located close to pHP are frequently excluded from the analysis as reads mapped at multiple positions. This situation is represented by the track labeled with ‘NSR’ or no signal region (the second track from the bottom). Accordingly, only restricted number of Ts2 toposites and Ts2/DSP sites will be detectable even if topo IIβ is actually acting on them. It is also possible that a significant portion of pHPs is in unknotted state at the time of eTIP-seq analysis in CGN culture. We speculate that one of the functional significances of repetitive sequences, especially LINEs, is to provide one level of genome DNA compaction through topo II-catalyzed knotting. Although we have not applied eTIP-seq to other cell types or different phase of CGN differentiation, it would be safe to suppose that in terminally differentiating neurons topo IIβ is involved in mainly knot-resolving reaction, which is probably reflected by the decondensation of global chromatin structure (Fig. 7a-d). Also clearly demonstrated in Fig. S14 is that the pHP-rich region is located in AT-rich area flanked by GC-rich area enriched with protein-coding genes (gene city). This pattern is generally observed throughout the genome.

### Genomic features that coincide with the predicted homologous pairs

We found that the density profile of Ts2 toposites and the position of Ts2-hotspots showed remarkable similarities to that of the segmental duplication (Fig. S15). Patterns for only chromosome 1 are compared here but the similarity holds throughout other chromosomes. Segmental duplications, tentatively defined as >90% sequence identity and >10 kb length, have common features in mammals and had been mapped in several species including the rat [49]. The duplication occurs as tandem and tightly clustered intrachromosomal regions of megabases long termed “duplication blocks”. Forty-one blocks that comprise ∼3% of the genome were identified in rat. Several gene groups that are commonly duplicated in mammals have been identified. They include the genes involved in drug detoxification (Cyp), olfaction (Olr, Vomr), and sperm competition (Smok, Spetex). In agreement with this finding, all these genes are found to associate with Ts2-hotspots (Table S4).

SNP density maps for rat chromosomes are available from the rat BAC browser website of Riken. As shown in Fig. S15, the SNP rate profiles for two rat strains closely correlate with Ts2 toposite-dense regions. This observation is quite comprehensible since point mutations occurring in duplicated sequences without significant function would have a great chance to survive during long period of time to accumulate as SNPs.

When the pHP map of Chr1 was compared to other density maps (Fig. S15), again, high-density regions of pHP showed a good agreement with other high-density peaks. This is also reasonable because duplicated sequences should be counted as homologous pairs. Taken together, the present study demonstrates the mechanistic link between DSP targets of topo IIβ and other genomic features that have long been known.

### The Ts2/DSP site is a key determinant of topo IIβ-dependent gene expression

The TSS-associated Ts1/PSP sites are likely to be involved in transcriptional regulation. Although they do have positive correlation with the level of transcription, we were unable to find any evidence for their involvement in the regulation of A1/C3 genes. After several trial-and-errors, we have been convinced that Ts2/DSP sites must be considered to solve the problem. Instead of homologous Ts2 toposite pairs experimentally detected, we tried to use the pHP information to analyze the topo IIβ dependency. When relative number of genes that contain pHP in their corresponding EU was calculated for each expression group (Fig. S12b), A1/C3 genes were significantly enriched with pHP-positive genes as compared to other gene groups. In contrast to the homologous Ts2 toposite pairs (Fig. S12a), however, no significant difference was observed between direct and inverted repeats. As length of A1/C3 gene EUs are significantly longer than that of other gene groups (Fig. S11a), the number of topo IIβ target (HP) associated with EU should be proportionally higher to confer the topo IIβ-dependency.

We think that genes in A1 and C3 gene groups, both nerve-related genes but expressed in different phase of neuronal development, are predesigned to have long expression unit in an evolutional context. Although this notion is somewhat different in nuance, the result itself does not contradict the previous reports that transcription of long genes is more dependent on topo II activity due to the necessity of resolving topological problems [27, 36]. Interestingly, long genes (>250 kb) bound by topo IIα in embryonic stem cells (ESC) are transcriptionally silenced and becomes bound by topo IIβ to be activated in terminally differentiated neurons [50]. They suggest that topo IIα ‘primes’ developmental genes in ESC for subsequent activation upon differentiation. Our model is consistent with this idea if we suppose that the knots created by topo IIα in neural precursor cells persist in terminally differentiating cells and become targeted by topo IIβ for unknotting and activation (Fig. 9). The probability of having knotted structure in EU will certainly increase as EU length increases. A recent report shows in yeast cells that topo II controls the knotting-unknotting homeostasis of DNA [51]. The knotting of chromatin fibers by topo II brings about 5-fold volume compaction and unknotting leads to chromatin decondensation. This idea is also consistent with our model (Fig. 9).

There is an apparent paradox that topo IIβ is required for both up-regulation of A1 genes and down-regulation of C3 genes. Judging from the enrichment of homologous Ts2 toposite pairs (Fig. 8f), topo IIβ is likely to be acting on EU region of both gene groups at the culture day 2 (D2). However, they behave differently with respect to the alteration of genic chromatin structure in the process of differentiation (Fig. 8g). Although the Cntn4 gene (A1) shows topo IIβ-dependent migration toward more dispersed or open chromatin region at D5, Robo1 gene (C3) appears to stay in open chromatin region all the time. Further studies are required to clarify the control mechanism of C3 genes.

### Inter-chromosomal DSPs link between Sat I repeats

Inter-chromosomal chimeras are more abundant than intra-chromosomal chimeras (Fig. S4c). However, combinatorial analysis of linked toposites, similar to the one shown in Fig. 3c, revealed that only Ts3-Ts3 chimera (Ts3/Inter) has a sufficient statistical significance (Fig. S16a). Almost all the Ts3/Inter chimera ends overlap with Sat I repeats that are either clustered or not clustered. Non-clustered Sat I (solitary Sat I) detected in Ts3/Inter chimera ends are frequently overlapped with Ts3/PSP sites that are characteristically short and homogeneous in size (Fig. 4a, bottom). For detailed analysis, we tried to clone Ts3/PSP sites overlapped with solitary Sat I sites using PCR with unique sequence primers flanking the site and found that resulting clones did not contain Sat I sequence. It appears then that these Sat I sequences were inserted by error in the fragment assembly process and thus the map positions of these Sat I sequences are unreliable. A logical consequence of this is that chimera links between solitary Sat I sites are not always “inter-chromosomal”.

Clustered Sat I detected in Ts3/Inter chimera ends are overlapped with pericentromeric Sat I clusters that contain abundant Ts3/DSP chimeras. The Ts3/Inter chimera ends have high sequence homology as in intra-chromosomal DSP chimeras. Sat I subrepeat sequences are already homologous each other (Fig. S7a), but when sequence reads for Ts3/Inter chimera ends were compared with individual subrepeats, both ends were most likely to reside within the same subrepeat and least likely so between non-neighboring subrepeats (Fig. S16b and c). This result is consistent with the fact that the distance between crossover points and sonic breakage points fluctuate somewhat and corroborates again that the homologous pairing is indeed the determinant of the target selection by topo IIβ.

Although sequence information of centromeric region is absent from the reference genome, the pericentromeric Sat I repeats are surely one of the major targets of topo IIβ in terminally differentiating neuronal cells (Fig. 7g). Since these cells never enter another mitotic cycle, structural integrity of centromeres may not be maintained. Involvement of topo IIβ in the modulation of centromeric structure was suggested by the FISH analysis (Fig. 7k and l).

### Relevance to TAD and CTCF

A number of contact mapping techniques including Hi-C revealed that interphase chromosome is organized into large looped domains called TAD (topologically associating domain) that are demarcated by CTCF (CCCTC-binding factor) and cohesin ring bound at the base of the loop [52–55]. Within the same TAD, genomic segments placed at a long distance in the primary structure can interact frequently. Association of homologous DNA segments would be greatly facilitated as in enhancer-promoter interactions. The size of TAD varies in the range of 10^5^-10^7^ base pairs. Since very little topological link is detectable between TADs [55], size of Ts2/DSP (median lengh∼260 kb) may imply that the topo IIβ-targeted HPs are most likely to reside within the same TAD.

The CTCF dimer formed between the domain boundaries of TADs may bring the two strands of DNA into close proximity at the base of loops [55]. This situation facilitates the formation of duplex crossovers, which may serve as preferred action sites for topo II enzyme [41]. Recently, topo IIβ has been shown to interact with CTCF and cohesin at the topological domain borders [56]. The enzyme was also shown to be active at these borders, though only a fraction at a given time, by co-mapping with etoposide-induced double strand breaks that are detected by a genome-wide method called End-seq [57]. These studies suggest that DSP sites may reside close to CTCF binding sites. However, both Ts3/DSP and Ts2/DSP sites showed little overlapping with CTCF binding sites when ChIP-seq data of rat liver CTCF binding sites were used for analysis (Fig. S13b).

Another line of evidence suggests that topo IIβ introduces double strand breakage within the promoter region or TSS zone of a subset of genes and facilitates their transcription initiation [58, 59]. Involved genes undergo signal-dependent activation mediated by nuclear receptors [58] or stimulation-triggered expression of neuronal early-response genes, in that a fraction of topo IIβ sites overlaps with CTCF binding sites [59]. Related observation in the present study would be that Ts1/PSP is highly enriched in TSS zone (Fig. 4c).

To date, transcriptional regulation by topo IIβ has been attributed to its action within relatively narrow DNA regions like promoters or enhancers. However, the present study demonstrated that the enzyme modulates long distance interactions to alter the transcriptional state of genes.

## Conclusions

Using a novel technique that captures authentic enzyme-DNA intermediates, we mapped the genomic distribution of topo IIβ action sites. The enzyme operates in two distinct modes, PSP and DSP, which differ significantly not only in mechanistic sense but also in physiological consequences. Most remarkable finding is that DSP is mediated by pairing of remote homologous DNA segments. The result may provide a novel perspective on the cellular function of type II DNA topoisomerases.

## Methods

### Primary cell culture

Cells used in this study were prepared as described previously [11]. Briefly, cerebellar tissue was isolated from the Wistar rat at 8th day after birth. Dispersed cells were plated at 2.5×10^7^ cells per 100-mm plastic dish pre-coated with poly-L-lysine. Cells were incubated at 37°C under 5% CO_2_ in DMEM containing 10% FCS, 25 mM KCl, and 60 µg/ml kanamycin sulfate. After 12 h, medium was replaced with fresh medium supplemented with 10 µM cytosine arabinoside to suppress the glial growth. The medium change was repeated at 24 h and at every 48 h afterwards. Cells maintained in the culture are mostly postmitotic cerebellar granule neurons (CGN) that continue to differentiate *in vitro*. When it is required, topo IIβ activity was inhibited specifically throughout the culture period by daily addition of 10 µM ICRF-193 that also degrades the enzyme [60] (first addition was made at 12 h). Cells were harvested either at day1 (D1), day 2 (D2), day 5 (D5) or day 5 in the presence of ICRF-193 (D5+) depending on the experiment.

For microscopic studies, cells were grown on coverslips and fixed by directly adding pre-warmed paraformaldehyde solution into the cultivating medium at a final concentration of 4%, followed by incubation at 37°C for 30 min. Fixed cells were then rinsed with phosphate-buffered saline (PBS; 150 mM NaCl in 10 mM sodium phosphate: pH7.4) and stored in 70% ethanol at 4°C until use.

### eTIPa-seq

Samples were prepared by modifying the protocol described previously [11]. CGN cells cultured in 100-mm dishes for 2 days were treated with 0.5 mM etoposide (VP-16) in serum-free medium for 15 min. Cells cultured in etoposide-free medium were used as control. The cells on a dish were lysed with 750 µl of the buffer containing 1% Sarkosyl, 10 mM Tris-HCl: pH 7.5, 10 mM EDTA, and protease inhibitor mixture (Complete Mini, Roche) and passed 10 times through a 23 G needle. Then, concentrated CsCl (7 M) was added to a final concentration of 0.5 M. The lysate was sonicated with a microprobe (TAITEC VP-5S) for 15 sec x 4 (cooled on ice for 15 sec each time) at the power setting of 4 and mixed well with 3 volumes of TEST-100 buffer (10 mM Tris-HCl: pH 7.5, 10 mM EDTA, 100 mM NaCl, 0.1% Triton X-100, and protease inhibitors). The mixture was centrifuged for 15 min at 15,000 rpm at 4°C. Supernatants were filtrated through MILEX HV (0.45 µm, Millipore) and combined into one tube, and pre-cleared with Dynabeads Protein G (DYNAL, Invitrogen) that had been coated with mouse IgG for 2 h at 4°C. After sampling a small portion, topo IIβ cleaved complex in the lysate was immuno-captured overnight at 4°C on a rotating wheel with Dynabeads Protein G coated with a specific monoclonal antibody to topo IIβ (3B6). After immuno-capturing, beads were washed three times with TEST-200 (10 mM Tris-HCl: pH 7.5, 10 mM EDTA, 200 mM NaCl, 0.1% Triton X-100).

Washed beads were combined into one tube by magnetic separation, and treated with TEST-500 buffer (10 mM Tris-HCl: pH 7.5, 10 mM EDTA, 500 mM NaCl, 0.1% Triton X-100) at 4°C to elute the complex-associated DNA that is non-covalently bound. Beads were separated on a magnetic stand and the solution containing eluted DNA was transferred to a new tube (P2 fraction). The bead fraction (P1 fraction), P2 fraction and the input were treated with RNase A at 55°C for 30 min followed by proteinase K at 55°C overnight. DNA was purified by phenol/chloroform extraction followed by ethanol-precipitation with carrier glycogen. Concentration of purified DNA was measured by Quant-it PicoGreen dsDNA kit (Invitrogen).

Purified DNA fractions (P1 and P2) were sheared to ∼200 bp using Covaris S2 sonicator. The eTIPa-seq libraries for NGS sequencing were prepared using the TruSeq DNA LT Sample Preparation Kit (Illumina) according to manufacturer’s instructions. After 10 cycles of PCR amplification, quality of the library was monitored by Agilent 2100 BioAnalyzer. Paired end sequencing (100 bp each) was performed on Illumina HiSeq 2000.

### eTIPb-seq

Entire procedure is illustrated schematically in Fig. S4a. The immuno-captured topo IIβ-DNA complex (IP complex) bound to magnetic beads was prepared as in the eTIP-a procedure. After washing with TEST-200 buffer, beads were combined into a 1.5 ml-tube and washed further with NEB ligation buffer (50 mM Tris-HCl: pH 7.5, 10 mM MgCl_2_, 1 mM ATP, 10 mM DTT, 100 µg/ml BSA) using magnetic separation each time. Washed beads were suspended in 0.6 ml of DNA polymerase mix (54 units T4 DNA polymerase, 100 µM each of dNTPs, and proteinase inhibitors in NEB ligation buffer) and incubated at 37°C for 30 min to blunt the ends of DNA fragments associated with topo IIβ. For dA-tailing of 3’ ends, beads were washed twice with TEST-200 buffer and once with NEBNext dA-Tailing Reaction Buffer (NEB), and then re-suspended in 0.6 ml of NEBNext module mix (Klenow fragment, proteinase inhibitors in NEB reaction buffer) followed by incubation at 37°C for 30 min.

After washing twice with TEST-200 buffer and once with NEB ligation buffer, beads were incubated for 6 h at 22°C in 0.9 ml of adaptor ligation mix (12,000 units of T4 DNA ligase, 5% PEG-8000, 1.5 µM ligation adaptor DNA, proteinase inhibitors in NEB ligation buffer) to attach the adaptor to the ends of topo IIβ-associated DNA fragments. The ligation reaction was performed under continuous rotation on Hula mixer (Thermo Fisher Scientific). Beads were washed with TEST-200 buffer 3 times followed by NEB ligation buffer. Then, 5’ end of the ligation adaptor was phosphorylated by incubating the beads for 30 min at 37°C in phosphorylation mix (120 units of T4 polynucleotide kinase, proteinase inhibitors in NEB ligation buffer). After phosphorylation, the supernatant was removed and the beads were re-suspended in 3 ml of 2nd ligation mix (6,000 units of T4 DNA ligase, proteinase inhibitors in NEB ligation buffer). The 2nd ligation between the adaptors was performed over night at 16°C with continuous rotation on Hula mixer. The reaction was terminated on ice by adding EDTA to final concentration of 10 mM and the supernatant was removed by magnetic separation. The bead-bound DNA (eTIPb-seq library) was purified as in eTIPa-seq procedure.

The ligation adaptor was prepared by annealing complementary oligomer DNAs (Operon): 5’-GTTGGATCCGATA[Bio-dT]CGC-3’ (top strand) and 5’-GGCCGCGATATCGGATCCAACT-3’ (bottom strand). We designed the adaptor sequence by modifying the one published elsewhere [61]. Top strand contains a biotinylated deoxy thymidine (Bio-dT) and bottom strand has 3’-dT tail and 5’ overhang of GGCC for the 2nd ligation between adaptors. Top strand was first treated with T4 polynucleotide kinase (NEB) for 30 min at 37°C to phosphorylate the 5’ end. After repeating the process with newly added enzyme, it was inactivated by heating at 65°C. To anneal the oligomers, equimolar amount of bottom stand oligo was added and the tube was placed for 5 min in a boiling water, then allowed to cool slowly to room temperature. The completion of annealing was confirmed by polyacrylamide gel electrophoresis.

The eTIPb-seq library DNA was incubated with T4 DNA polymerase in the presence of dATP and dTTP in NEBuffer 2 (NEB) for 2 hours at 20°C to remove the biotinylated nucleotide from terminal adaptors. After phenol/chloroform extraction and ethanol precipitation with glycogen, purified DNA was sheared to ∼200 bp with Covaris and processed for end repairing. The dA-tailing and sequence adapter ligation were done with TruSeq DNA LT Sample Preparation Kit (Illumina) according to manufacture’s instructions. After terminating the reaction by incubation at 65°C to inactivate the enzyme, biotinylated DNA fragments were captured with Dynabeads MyOne Streptavidin C1 (LifeTechnologies) suspended in the binding buffer (5 mM Tris-HCl: pH 7.5, 0.5 mM EDTA, 1 M NaCl). Beads were washed 3 times with Tween wash buffer (5 mM Tris-HCl: pH 7.5, 0.5 mM EDTA, 1 M NaCl, 0.05% Tween-20) and suspended in Resuspension-buffer of TruSeq kit and directly used for PCR amplification (15 cycles with PCR primer cocktail and PCR master mix, Illumina). Amplification products were purified using AMPure XP beads (Beckman) according to manufacturer’s instructions. Quality of the resulting library was monitored using Agilent 2100 BioAnalyzer. Paired end sequencing (100 bp each) was performed on Illumina HiSeq 2000.

### ChIP-seq

The CGN cells at the 2nd day *in vitro* (D2) were fixed with 1% formaldehyde for 10 min and quenched with 125 mM glycine for 5 min at 25°C. Cells on dish were washed twice with PBS containing proteinase inhibitors, scraped off and harvested by centrifugation. The cell pellets were stored at −80°C until use. The frozen cell pellets were resuspended in 125 µl of SDS-lysis buffer (50 mM Tris-HCl: pH8.0, 140 mM NaCl, 1 mM EDTA, 1% Triton X-100, 0.1% sodium deoxycholate, 0.8% SDS, proteinase inhibitors) and incubated on ice for 10 min. The cell suspension was diluted with 375 µl of the lysis buffer without SDS (ChIP dilution buffer) and sonicated for 2 min with intermittent poses by ultrasonic disruptor (UD-201, TOMY) attached with a special microtip (power setting at 5). After repeating the sonication two more times, cell debris was removed by centrifugation at 20,000 x g for 10 min at 4°C. The supernatant was diluted 2-fold with ChIP dilution buffer and filtered through 0.45 µm membrane (Millex HV, Millipore) and pre-cleared with Dynabeads Protein G that had been pre-coated with mouse IgG for 3 h. The final lysate was used for immunoprecipitation with anti-SP120 monoclonal antibody (1-67D) bound to Dynabeads Protein G. The beads were collected by magnetic separation and washed consecutively with RIPA buffer (50 mM Tris-HCl: pH8.0, 150 mM NaCl, 1 mM EDTA, 0.1% SDS, 0.5% NP-40), 500 mM-NaCl RIPA buffer, LiCl buffer (10 mM Tris-HCl: pH8.0, 250 mM LiCl, 1 mM EDTA, 0.5% sodium deoxycholate, 0.5% NP-40), and 0.1% Triton in 10 mM Tris-HCl: pH8. Elution was performed twice with 100 µl of Elution buffer (1% SDS, 100 mM NaHCO_3_, 10 mM DTT) for 10 min at 25°C. After adding 5 M NaCl to a final concentration of 200 mM, the combined eluate was heated at 65°C for 4 h to reverse the cross-link. After treating the eluate with RNase A at 55°C for 30 min followed by Proteinase K at 55°C for 30 min, DNA was purified by use of QIAquick PCR purification kit (Qiagen). Sixty nanograms of purified DNA was subjected to size selection (100–600 bp range) by AMPure XP beads (Beckman) and fragmented by Covaris to generate ∼200 bp segments. The fragmented DNA was end repaired, dA-tailed and ligated with paired-end sequencing adapter. Purified DNA was amplified with 14 cycles of PCR using PE PCR 1.0 and PE PCR 2.0 primers (Illumina). The concentration and size distribution of library DNA after PCR amplification was determined by BioAnalyzer profiles (Agilent Technologies) and quantitative PCR, and the libraries were paired-end sequenced (100 bp each) on Illumina MiSeq platforms.

### mRNA-seq

Total cellular RNA from D1, D5, and D5+ cells were prepared as described previously [11]. RNA-Sequencing libraries were generated using the Truseq SBS kit v3-HS (Illumina) with a polyA selection step. The samples were sequenced (75-bp paired-end sequencing) on one lane of HiSeq 2000 (Illumina) per sample. Expression levels of genes were determined based on FPKM values. Briefly, the paired-end mRNA-seq reads were aligned against the rat genome assembly rn4 using TopHat version 2.0.8 with optional settings of 0 bp for the “mate inner distance” and 1 Mb for maximum intron length. The remaining options were set to their default values. We have supplied transcript annotations from RefSeq and xenoRefSeq at the UCSC genome browser site, which specifies known splice junctions and exon boundaries. Since xenoRefSeq contains vast variety of species, we extracted and used the data from *Homo sapiens* and *Mus musculus* to construct exRefSeq as described previously [11]. Expression levels (FPKM) of exRefSeq genes were calculated with Cufflinks program (v. 2.2.1) [62]. Genes were classified into 9 groups based on two factors: the rate of induction (FPKM at D5/ FPKM at D1) and the rate of susceptibility to topo II inhibitor (FPKM at D5/ FPKM at D5+). To avoid infinite values, 0.1 count had been added to all FPKM values before the calculation. Genes with very low FPKM values (<1.0) in any conditions were classified as group ‘D’ to represent unexpressed genes. A list of exRefSeq genes sorted by the name of expression group and gene length is given in Table S6. The description column was created by use of the g:profiler program [63].

For gene ontology (GO) analysis, significant GO terms (p<10^-4^) enriched in each expression group were extracted by GOstat2 program (http://gostat.wehi.edu.au/cgi-bin/goStat2.pl). Neuronal genes (4351 in total) were selected from the NCBI gene database as described [11]. ASD genes (885 in total) were downloaded from AutDB, an autism gene database [64]. The GO data obtained are summarized in Table S7.

To corroborate the precision of grouping, relative mRNA levels were compared by RT-qPCR at D1, D3, D3+, D5, and D5+ for representative genes from group A1 *(Camk2d, Itpr1, Stx1a, Pkia*), group A2 (*Gabra1, Grm4*), and group B2 (*Matr3, Ncl, Cat, Actb*). All results were consistent with the classification criteria and also with the microarray results published previously [11].

### FAIRE-seq

We used the FAIRE-seq technique to map the genomic regions that are devoid of associated proteins and thus accessible to protein factors including topo II (open chromatin). Except for some modifications, we followed the experimental procedures for FAIRE (formaldehyde-assisted isolation of regulatory elements) described previously [65]. CGN cells grown on culture dishes for 1 day (D1), 5 days (D5), and 5days in the presence of ICRF-193 (D5+) were fixed with 1% formaldehyde for 5 min at room temperature. Washed cells were suspended in a hypotonic buffer containing protease inhibitors and homogenized with Dounce homogenizer. After centrifugation, the pelleted crude nuclei were lysed in the lysis buffer containing 1% SDS, and sonicated on TOMY ultrasonic disruptor attached with a special microtip (power setting at 3) for 2.5 min x 3 cycles. After aliquoting for input control DNA, the sonicated lysate was subjected to phenol/chloroform extraction and the water phase was isolated. DNA in the solution was recovered by ethanol precipitation, dissolved in buffer and treated with RNase A followed by proteinase K. To reverse the cross-link, the mixture was incubated at 55°C for 60 min and then at 65°C overnight. Finally, DNA was purified using QIAquick PCR purification kit (Qiagen). For preparing input control DNA, the step of cross-link reversal was performed before the phenol/chloroform extraction.

DNA samples were subjected to 2% agarose gel electrophoresis to monitor the fragment size, which was found to be an acceptable range of ∼200 bp. NGS protocols were the same as used for eTIP-seq.

### Mapping and peak-calling of sequence data

For eTIPa-seq and ChIP-seq analyses, the FASTQ paired-end read data were mapped by BWA against rat reference genome (UCSC rn4) and then filtered to remove frequently encountered experimental artifacts. If the sequencing adaptor sequence was present at the end of the read, the adaptor sequence was trimmed from the read sequence. Only paired reads with correct orientation (forward/upstream and reverse/downstream) and with reasonable distance (less than 500 bp) that is expected from the distribution of original fragment length were selected. To simplify the analysis, fragment length was normalized to 200 bp, the median length. Uniquely mapped pairs were used for peak calling. R-package ZINBA version 2.02.03 was used to find peaks by setting ‘refinepeaks’ to 1, ‘threshold’ to 0.5, ‘extension’ to 200 and other factors to default [66]. Selected fragments that are flanked by uniquely and correctly mapped read pairs were used to construct wiggle track formats. Pileup data was calculated by ‘basealigncount’ function of ZINBA with extension of 200.

For eTIPb-seq analysis, the FASTQ paired-end read data were mapped by BWA to rn4 separately as ‘read 1’ and ‘read 2’. Sequence reads with the ligation adaptor sequence at 5’ end were discarded. If the ligation adaptor sequence appears within the read sequence, the adaptor sequence and the following sequence were trimmed from the read sequence. If the remaining sequence is less than 17 nucleotides, the entire read was discarded. Uniquely mapped pairs were used for further analyses.

For FAIRE-seq analysis, the wiggle data was formatted through the same procedure as used in eTIP-seq.

### Genome-wide prediction of homologous pairs

For each chromosome starting from the left end, Watson strand of rat genomic sequence (rn4) was sequentially divided into 2 Mb-bins that overlap with neighboring bins by 1 Mb. Using EMBOSS DotMatcher program (http://emboss.sourceforge.net), we searched the position of homologous regions pairwise within every bin by setting optional parameters (window size: 2000 bp, threshold homology score: 4000). The resulting graph data drawn to every bin were exported into a text file as digital data. The homologous pairs detected are referred to as ‘direct repeat’. Next, reverse complementary sequences (Crick strand) were generated by Revseq program in the EMBOSS package and the homology search of Watson vs. Crick strand was done similarly to detect homologous pairs that are called ‘inverted repeats’.

As DotMatcher results contain a great deal of redundancy, redundant homologous pairs were removed from the list as follows. The “self-homology”, which is aligned on diagonal line in the dot matrix graph, and tandem repeats overlapping off-phase were discarded. Only homologous pairs residing within 1 Mb were counted. Except for the left-most bin, homologous pairs in the left half of the 2 Mb-bin were discarded because they are already counted in the previous bin. Since the same homologous pairs are arranged symmetrically against the diagonal, only those positioned above the diagonal were counted.

### Analysis of topo IIβ-DNA cleaved complex formed by hydrogen peroxide

CGN cells at the 2nd day *in vitro* (D2) were treated with 30 mM hydrogen peroxide (H_2_O_2_, Sigma) in serum-free medium for 30 min to trap topo IIβ-DNA cleaved complex [67]. The H_2_O_2_-treated cells were lysed with sarkosyl and processed as in the eTIPa-seq procedure described above except for sonication time (15 sec x 2 instead of 15 sec x 4). The topo IIβ-DNA cleaved complex was immuno-captured (IP) with the anti-topo IIβ antibody bound to Dynabeads Protein G. After washing with TEST-200 buffer followed by 1x CutSmart buffer (NEB), the beads were incubated with 1.25 unit/μl *Mbo* I (NEB) at 37°C for 4 h in 1x CutSmart buffer containing protease inhibitors. The beads were washed once with PBS containing 0.1% NP-40 and then with 1x T4 DNA ligase reaction buffer (NEB) before incubation with or without 2 unit/μl T4 DNA ligase (NEB) at 16°C for 12 h in the NEB ligase reaction buffer containing protease inhibitors. The reaction was terminated on ice by adding EDTA to final concentration of 10 mM and the beads were collected by magnetic separation.

The *Mbo* I-treated beads, ligase-treated and untreated beads, as well as the IP input were incubated with 50 ng/μl RNase A (Qiagen) at 55°C for 30 min in a buffer containing 10 mM Tris-HCl: pH8.0, 10 mM EDTA and 0.5% SDS followed by proteinase K at 55°C overnight. DNA was purified by phenol/chloroform extraction followed by ethanol-precipitation with carrier glycogen. Concentration of purified DNA was measured by Quant-it PicoGreen dsDNA kit (Invitrogen).

The purified DNA was subjected to PCR analysis as follows. To see whether the DNA in DSP chimeras detected in etoposide-treated cells is also present in immuno-captured DNA obtained from the H_2_O_2_-treated cells, the *Mbo* I-treated template DNA was amplified with primer pairs targeting upstream and downstream regions of three Ts3/DSP chimeras (ID21, ID130, and ID255 listed in Table S5). PCR amplification was performed using 1 ng of template and AmpliTaq Gold 360 Master Mix (Thermo Fisher) for 30 cycles (20 s at 95°C, 20 s at 55°C and 30 s at 72°C) in the presence of the primer pairs listed in Table S8. PCR products were separated in 2% agarose gel and visualized by ethidium bromide staining. To see whether or not DSP chimeras detected by eTIPb-seq are also formed in H_2_O_2_-treated cells, purified DNA from the ligase-treated and untreated beads were subjected to PCR amplification using primer pairs that amplify only ligation products between distant regions of the Ts3/DSP chimeras (ID21, ID130, and ID255). Six ng of the template DNA was subjected to the PCR amplification and the products were separated by electrophoresis in 1% agarose gel and visualized by SYBR green I.

### *In vitro* reaction with purified topo IIβ that mimics the eTIP procedure

Flag-tagged topo IIβ was expressed in the human embryonal kidney cell line HEK293E cells transfected with pFlag-top2b plasmid encoding the full-length rat topo IIβ, and purified by immunoprecipitation with Dynabeads Protein G that had been pre-coated with anti-Flag antibody as described previously [35]. As a substrate of topo IIβ enzyme reaction, pUC18 plasmid DNA containing *E*. *coli* DNA insert was prepared as follows. *E*. *coli* DNA fragments prepared by sonicaton were blunt-ended with S1 nuclease and cloned into the *Sma* I site of pUC18 plasmid. One of the clones that contained about 1.4 kb *E*. *coli* DNA fragment was used in the following experiments.

Flag-tagged topo IIβ on 6 μl of Dynabeads was incubated with 200 ng of supercoiled substrate DNA in the presence or absence of 100 μM etoposide at 30°C for 30 min in 15 μl of reaction mixture containing 50 mM Tris-HCl: pH 8.0, 120 mM KCl, 10 mM MgCl_2_, 0.5 mM dithiothreitol, 0.5 mM EDTA, 0.5 mM ATP and 30 μg/ml BSA. Then, concentrated sarkosyl and CsCl were added directly to final concentrations of 1% and 0.5 M, respectively. After dilution with 3 volumes of TEST-100 buffer, the bead-bound enzyme-DNA complex was separated and treated consecutively as follows with magnetic separation at each step: washing with 100 μl of 1x CutSmart buffer (NEB) containing protease inhibitors, cutting with *Eco* RI-HF (NEB) and *Sal* I-HF (NEB) at 37°C for 30 min in 15 μl of 1x CutSmart buffer containing protease inhibitors, and eluting with 15 μl of TEST-500 buffer on ice for 5 min. Bead-bound DNA (bound to topo IIβ) and unbound DNA (free in solution) were treated with 1% SDS, 50 mM EDTA and 100 μg/ml proteinase K. Samples were separated by electrophoresis in 1.2% agarose gel containing 0.5 μg/ml ethidium bromide.

### Microscopy and image processing

All fluorescence images were acquired at a resolution of 1388 x 1040 pixels by an AxioCam MRm camera installed on an Axiovert 200M inverted fluorescence microscope with a C-Apochromat 63x/NA 1.2/water immersion objective and an ApoTome structured illumination optical sectioning system, using AxioVision 4.5 software (Carl Zeiss). For three-dimensional (3D) DNA-FISH analysis of the cell nucleus, serial 25 optical sections were collected from 100 nuclei at 0.275 µm z-axis intervals through the entire nucleus and saved as 24-bit red-green-blue (RGB) color TIFF. The following steps were performed with ImageJ software 1.49v (NIH, Bethesda, MD). First, the original z-stack was cropped to obtain regions of interest (ROIs) of 150 x 150 pixels square centered on the nucleus. The resulting cropped z-stack was then converted to the compositions for three 8-bit monochromatic R (PLA signal), G (FISH signal), and B (Hoechst signal) channels. The B channel was binarized with a manually chosen threshold value and the binary image was used as a mask on the corresponding R and G channels to outline the border of each nucleus. Three-D objects of interest in each z-stack of the R and G channels were segmented and recorded in the ‘ROI Manager 3D’ plugin with a constant threshold for each channel. After creating a list of ROIs, the ‘Measure 3D’ option in the plugin was applied to measurements of the size and number of PLA and DNA-FISH foci in individual nuclei.

For 3D colocalization analysis, the area of interest was cropped from the original z-stack and split into the RGB channels as described above. The R and G channel images were then re-combined as a dual-color z-stack by the aid of the ‘Merge channels’ command. The B channel z-stack was used to create a binary mask that discriminates the pixels of the foreground from those of the background objects in the dual-color z-stack. Co-localization masks were made by using the ‘RG2B colocalization’ plugin. The resultant binary z-stack was added to the ‘ROI Manager 3D’ plugin and the number of colocalized PLA and DNA-FISH foci per nucleus was determined in 3D space by using the ‘Measure 3D’ option in the plugin.

### Voxel intensity measurement of Hoechst 33342-stain

Measurements were performed by ImageJ software. The B (Hoechst) channel of each cropped z-stack was converted to 8-bit grayscale and subjected to gray-level histogram quantification as follows: (1) unwanted background objects in each stacks were masked using the ‘Paintbrush’ tool; (2) the resulting stacks were normalized by linearly stretching the voxel intensities over a full range of 256 gray-levels with the ‘Histogram’ tool (termed normalized-stacks); (3) the normalized-stacks were smoothed by applying a mean filter (filtered-stacks); (4) using the normalized-stacks, three-dimensional nucleus segmentation was automatically performed by the aid of the NucleusJ plugin ‘Nucleus segmentation’ function [68] (segmented-stacks); (5) the corresponding filtered- and segmented-stacks were combined by using the ‘Subtract in the Image calculator’ (combined-stacks). The segmented-stack was used as a mask that discriminates the voxels of the foreground signals from those of background noises in the corresponding filtered-stack; (6) by using the ‘Histogram’ tool, intensities of all voxels within the combined-stacks were computed, expressed as histogram plots, and exported as text files.

To estimate the degree of chromatin condensation at various culture days, the staining intensity for 100 nuclei was recorded (8 bit, 256 grayscale). After normalizing the intensity to total intensity per nucleus, intensity versus median frequency curve was plotted. The isosbestic point for D1, D5, and D5+ curves indicates the border between condensed and dispersed chromatin regions. The condensed region corresponds to the right side of isosbestic point.

### Proximity ligation assay (PLA) and DNA-fluorescence in situ hybridization (FISH)

To improve detection sensitivity of a combined technique of PLA and FISH, cells on coverslips were air-dried, heated in 10 mM sodium citrate (pH6.0) for 5 min by using a microwave oven set at 500W, and allowed to cool to room temperature. After washing with PBS, samples were blocked with PBS containing 0.3% Triton X-100, 10% goat serum, and 1% BSA for 30 min and incubated with a mixture of 40 µg/ml mouse anti-topo IIβ monoclonal antibody (3B6) and 5 µg/ml rabbit anti-SP-120 polyclonal antibody [20] at 37°C overnight. Following washing with PBS, PLA was conducted using the mouse PLUS and rabbit MINUS Duolink In Situ PLA kit (Olink Bioscience). Ligation and amplification of DNA was done with the Duolink In Situ Detection Reagents Red (Olink Bioscience) according to the manufacturer’s manual. The resulting PLA signals were post-fixed in 2% paraformaldehyde in 1x Duolink In Situ buffer B for 10 min. The sample was then washed in 2x SSPE (0.3 M NaCl, 23 mM NaH_2_PO_4_, 2 mM EDTA; pH7.4), equilibrated in 50% deionized formamide and 2x SSPE for 30 min, and processed for FISH preparation.

For detecting the satellite I (Sat I) loci, a fluorescein isothiocyanate (FITC)-labeled DNA probe was prepared by means of a PCR-based method. Among dNTP substrates, 1/3 of dTTP was substituted by FITC-12-dUTP (Roche). We used a rat Sat I sequence cloned into pBluescript as the PCR template. Upon sequencing, this clone showed 97.3% sequence identity to the canonical Sat I sequence (Fig. S7a). The primer pair used for amplification was the same as the one used for initial cloning (5’-ATCGAATTCACAGAGAAACAGTG-3’ and 5’-ATCAGTTAGTTCCCAGTAGCCTG-3’). After amplification under standard conditions, agarose gel electrophoresis of the product revealed a single band of expected size (∼370 bp) without detectable levels of background. The PCR products (∼1 µg) were precipitated in ethanol with 50 µg of sonicated fish sperm DNA (Roche Diagnostics) and resuspended in 20 µl of a hybridization buffer (50% deionized formamide, 10% dextran sulfate, and 2x SSPE). Hybridization was done by floating sample coverslips, cell-side down, on drops of hybridization solution (∼100 ng probe/10 µl) in a moist chamber. Following pre-heating at 37°C for 1 h, cell nuclear and probe DNAs were denatured simultaneously at 75°C for 4 min and subsequent hybridization was carried out at 37°C overnight. The samples were then washed three times in 2x SSPE at 37°C for 10 min, twice in 0.1x SSPE at 55°C for 5 min, and once in 2x SSPE at room temperature for 2 min. To detect FITC-labeled probes, an Alexa Fluor 488-conjugated goat anti-fluorescein/Oregon green IgG (Molecular Probes, diluted in 1:200 in 2x SSPE containing 0.1% Tween 20 and 1% BSA) was employed. Cell nuclei were then stained with 10 µM Hoechst 33342 in 2x SSPE for 10 min, after which the sample coverslips were mounted onto glass slides in a ProLong Antifade kit (Molecular probes).

### Analysis of Hoechst-staining intensity where PLA or FISH signals reside

Using ImageJ software, the voxel grayscale values of Hoechst-fluorescence being in the same position as the centroid of FISH-signals were measured as follows: (1) original z-stacks were cropped and split into separate RGB channels (8-bit grayscale each) as described above; (2) the B channel stack images (Hoechst) were processed as described above for Hoechst-voxel intensity and were used to define the nuclear ROIs; (3) a median filter with an appropriate radius was applied to the R and G channel stack images (FISH-signals) to reduce salt-and-pepper noise; (4) by using the 3D Object Counter plugin’s Map commands ‘objects’ and ‘centroids’, FISH signal positions within a z-stack were computed as the 3D centroid coordinates of the thresholded signals and created a new 3D binary image stack having only two pixel values of 0 for black (background voxels) and 255 for white (FISH-centroid voxels). The 3D binary image stack was employed as a mask to discriminate the Hoechst voxels being located at FISH-centroid coordinates from the rest of the Hoechst images; (5) the B channel stacks obtained at process (2) were then masked by their corresponding 3D binary masks with the ‘AND create’ operation of the Image Calculator; (6) the voxel grayscale values of Hoechst-fluorescence being situated at FISH-centroids were measured using the ‘Analyze Particles’ function under a constant threshold range from 1 to 255 across all images. In this work, a total of 100 nuclei were semi-quantified per condition and the resultant data were imported into Microsoft Excel software for statistical analysis.

### Two color FISH for detection of genic regions of *Cntn4* and *Robo1*

Hybridization was performed basically as above except that the hybridization solution consisted of a mixture of two different labeled-probes (150 ng each), 15 µg of fish sperm DNA (Roche), and 6 µg of Rat-Hybloc competitor DNA (Applied Genetics Labs), in 50% formamide, 10% dextran sulfate, and 2x SSPE. We used digoxigenin- or biotin-labeled probes prepared from rat bacterial artificial chromosome (BAC) clones using a Digoxigenin or a Biotin Nick Translation Kit (Roche). Contiguous sets of BAC clones covering almost the entire gene region of interest were used: RNB2-011E20, RNB2-037D06, RNB1-354M17, and RNB2-043J12 for *Cntn4*; RNB1-126D17, RNB1-434C21, RNB1-309N01, and RNB2-261M17 for *Robo1*. All BAC clones were obtained from the RIKEN BioResource Center (Tsukuba, Japan). Prior to hybridization, nuclear DNA and probe DNA were denatured simultaneously at 80°C for 5 min. Detection of digoxigenin- and biotin-labeled probes was carried out with DyLight 594-conjugated goat anti-Digoxigenin (5 µg/ml; Vector Labs) and DyLight 488-conjugated streptavidin (5 µg/ml; Vector Labs), respectively. Nuclei were counterstained with 10 µM Hoechst 333342 in 2x SSPE for 10 min.

### Statistical analysis

For the continuous variables of interest, the Mann-Whitney U test with two sided significance (wilcox.exact function of R-package, exactRankTests) was used if the overall difference was statistically significant. Box plots were drawn by R with default settings. The chi-square test or Fisher’s exact test (chisq.test or fisher.test function of R) was used to examine the differences in the categorical characteristics and changes in the groups.

## List of abbreviations

FPKM: Fragments per kilobase per million reads
TSS: Transcription start site
MHC: Major histocompatibility complex
SNP: Single nucleotide polymorphism
CGN: Cerebellar granule neuron
NGS: New generation sequencer
FISH: Fluorescence in situ hybridization
PLA: Proximity ligation assay
exRefSeq: Extended RefSeq
PSP: Proximal strand passage
DSP: Distal strand passage
pHP: Predicted homologous pair

## Declarations

### Acknowledgements

We thank S. Sugano, Y. Suzuki, and Y. Kohara for providing us with NGS sequencing opportunities and suggestions on sequence analysis.

### Availability of data

The raw data obtained in this study are available from DDBJ Read Archive (http://ddbj.nig.ac.jp//DRASearch/) under accession numbers of DRA007399 for eTIPa-seq, DRA007479 for eTIPb-seq, DRA007375 for SP120 ChIP-seq, DRA002525 for mRNA-seq and DRA007436 for FAIRE-seq.

### Authors’ contributions

MM designed and conducted laboratory experiments for SP120 ChIP-seq, performed bioinformatic analyses on sequence data generated by eTIPa-seq, eTIPb-seq, FAIRE-seq, and mRNA-seq experiments. RF conducted eTIPa-seq experiments and optimized the eTIPb-seq procedure. OH performed PLA and FISH experiments and analyzed the data. KS conducted FAIRE-seq and mRNA-seq experiments and contributed to the construction of exRefSeq for gene expression analysis. NH and RK acquired the sequence data for ChIP-seq. JK and MT contributed to the mapping of sequence reads and ZINBA analysis for eTIP-seq and ChIP-seq. KMT analyzed and interpreted all the results, especially data from eTIP-seq, and revised the manuscript critically to ensure textual consistency. KT conceived and designed the study, analyzed and interpreted the results and wrote the manuscript with help from MM, RF, OH and KMT. All authors read and approved the final manuscript.

### Ethics approval and consent to participate

Not applicable.

### Consent for publication

Not applicable.

### Additional files

Additional file 1: Supplementary. This file contains a detailed description of basic mapping techniques used in the study and comprehensive analyses with additional data. Supplemental figures S1-S16 along with figure legends and references cited are also included. (PDF 3600 kb)

Additional file 2: Table S1. (XLSX 7500 kb)

Additional file 3: Table S2. (XLSX 18 kb)

Additional file 4: Table S3. (XLSX 13 kb)

Additional file 5: Table S4. (XLSX 40 kb)

Additional file 6: Table S5. (XLSX 53 kb)

Additional file 7: Table S6. (XLSX 2600 kb)

Additional file 8: Table S7. (XLSX 209 kb)

Additional file 9: Table S8. (XLSX 35 kb)

