## Supplementary for "Topoisomerase IIβ targets DNA crossovers formed between distant homologous sites to modulate chromatin structure and gene expression"

### 1. Notes on methodology

This section contains information on fundamental principles and procedure outlines for the newly developed techniques used in the study. Supportive experimental evidence is also provided below. Detailed experimental procedures are described in Methods of the main text.

#### Classification of topo IIβ target sites (eTIPa-seq)

Treatment of the IP complex with 0.5 M NaCl separates the DNA into P1 and P2 fractions (Fig. 1). P1 is mostly composed of DNA fragments covalently bound to the enzyme whereas the association of fragments released in P2 is non-covalent in nature. A large proportion of recovered DNA fragments distributed between 0.5 kb and 3.0 kb under these conditions (Fig. S1a, b). Upper limit of the smear (~3 kb) reflects the maximal size of sonication-resistant DNA, which was constant between experiments. Densitometric scanning of the smear showed that the fragment size peaks around 1.5 kb (Fig. S1b). Relative DNA yields from both fractions showed a strong dependency on etoposide treatment, indicating that the fragments are originated from the topo IIβ reaction intermediate, not from nonspecific interactions (Fig. S1c). Purified DNA from these fractions were subjected to sequencing on NGS. Sequence reads from eTIPa-seq experiments were processed as in Fig. S1d. Paired-end reads for the DNA fragments purified from P1 and P2 fractions were mapped on the rat reference genome, UCSC rn4 (Baylor Build 3.4, November 2004). We have adopted this version because more recent versions (rn5 and rn6) contained multiple rearrangements probably due to erroneous fragment assemblies. The mapped fragments between the read pairs were analyzed by a peak-finding program called ZINBA [1]. We defined 3 distinctive categories of topo IIβ-targeted sequences called toposites (Ts1, Ts2, and Ts3) by using ZINBA peaks for P1, P2, and their overlaps (Fig. S1d, bottom). In summary, toposites are regions with P1 peak alone (Ts1), P2 peak alone (Ts2), and both P1/P2 peaks (Ts3). Genomic position and site length of resulting toposites are listed in Table S1.

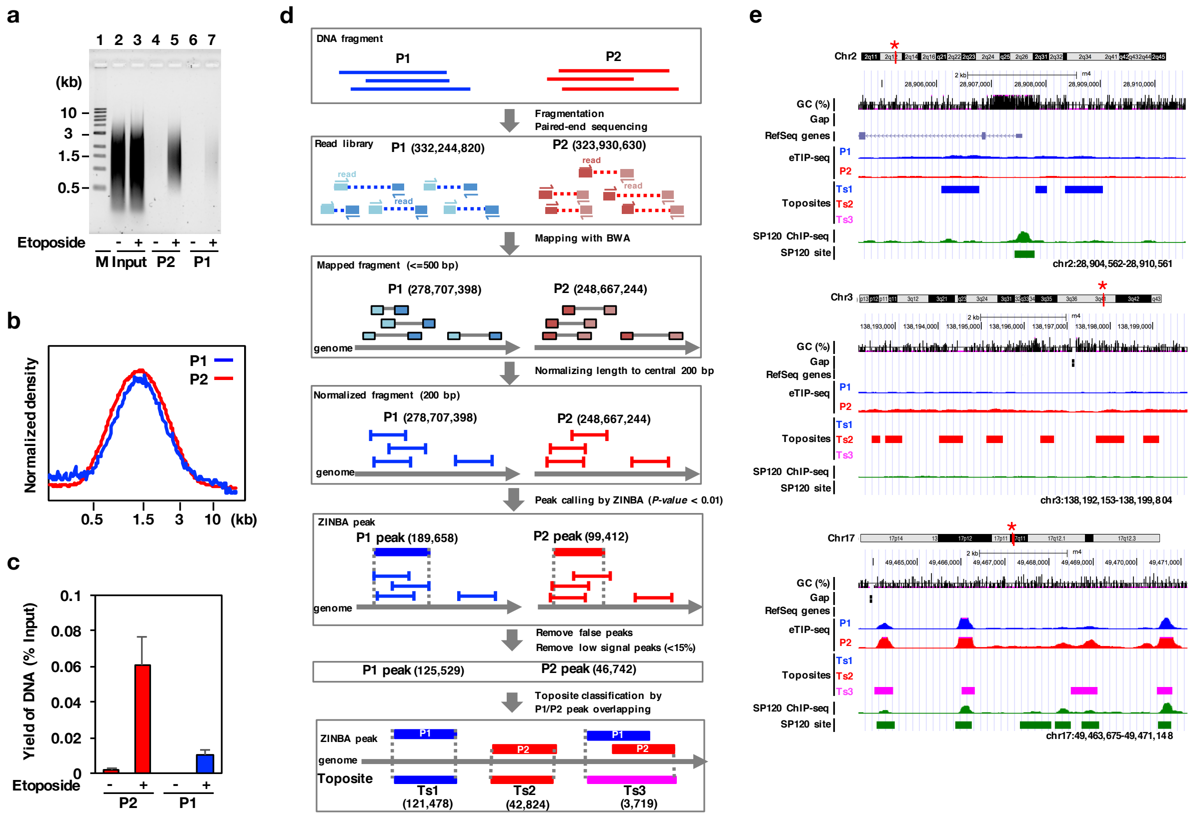

**Fig. S1** Partitioning of DNA fragments among fractions and toposite assignments in eTIPa-seq. **a** The fragment size of DNA from P1/P2 fractions as determined by agarose gel electrophoresis. The same procedure was conducted in the presence (+) or absence (-) of etoposide. M, size marker; Input, sonicated DNA before immunoprecipitation. **b** Fragment size distribution of P1/P2 fractions from etoposide-treated cells. There was no difference between the normalized densitometric profiles of DNA from P1/P2 fractions. **c** Relative amounts of DNA recovered from P1/P2 fractions. **d** Schematic representation of the entire signal processing. DNA amounts used for sequencing were 485 ng for both P1 and P2 fractions. Numbers in parentheses indicate actual signal numbers selected in each step. For P1/P2 overlaps, the sum area is regarded as a Ts3 peak if more than half portion of the shorter peak region is overlapped. P1 and P2 peaks showed little self-overlapping. **e** Regions enriched with toposites are shown on the UCSC browser (from top to bottom; Ts1, Ts2, and Ts3). Asterisks on the chromosome band indicate selected regions. Wiggle patterns for P1, P2, and SP120 are also shown with peak positions identified by a peak-caller ZINBA. Typical regions exhibiting the distinction between the three toposites were selected here.

Mammalian topo II binds and acts preferentially on DNA crossovers formed between two duplexes. Handedness of the crossover favored by the enzyme is predetermined by structural requirements. In the scheme shown in Fig. S2a (left), topo II approaches from the bottom of right-handed DNA crossover, whose wider angle is filled with the enzyme's identical subunits. Configuration of the two strands illustrated here designates that G-segment is already held by the enzyme and T-segment is approaching to the N-gate [2]. The same crossover configuration at the DNA gate of topo IIβ has been shown recently by a crystallographic study [3]. In the present study, we adopt this model as a presupposition to interpret the data obtained. It is worth noting that the crossover handedness is the same when the enzyme approaches from either side of the cross but the cleaved strand (G-segment) differs in each case (Fig. S2a, right).

The table shown in Fig. S2b represents all possible combinations of G- and T-segments that are formed by the enzyme approaching from either side. The majority of DNA fragments associated with IP complex is sheared to less than 3 kb by sonication (Fig. S1b). Therefore, G- and T- segments can be contiguous if they are located within 3 kb on the same chromosome, indicating that the strand passage event occurred between nearby DNA segments (termed PSP for proximal strand passage). As shown in Fig. S2b, the complex-bound DNA forms a loop (PSP loop). If G- and T- segments are discontinuous, however, they are recovered separately in P1 and P2 fractions, respectively. In this case they can be derived from distant locations on the same chromosome or even from different chromosomes (termed DSP for distal strand passage).

Reversal of DNA breaks in G-segment during the eTIP procedure complicates the situation. Under the eTIP conditions, some G-segments are re-ligated to lose covalent attachment to the enzyme and thus gets recovered in P2 fraction. In this case the site will be assigned to Ts2. The resealing probability of G-segments appears to depend on local GC content because the topo II-linked DNA breaks are generally more stable in GC-rich sequences than AT-rich sequences as reflected by average GC contents of toposites (Fig. 2c). The partitioning pattern of DNA fragments in P1 and P2 fractions defines the toposites. Wiggle patterns for these DNA fractions exhibit the way how toposites are actually assigned (Fig. S1e). Most importantly, Ts1 sites are exclusively composed of G/T contiguous fragments with stable DNA breaks and Ts2 sites lack the covalent linkage with the enzyme due to unstable breaks. Not all the breakage is reversed but remains intact so that the G-segment is recovered in P1 fraction as in Ts1 and Ts3 toposites. GC content of these sites are either GC-rich or neutral. The Ts3 site is unique in that genomic segments at a particular position are recovered both in P1 and P2 fractions, indicating that at these sites there is no strong bias for the selection of G/T-segments. The Ts3 site is also unique because it contains a distinct population of sonication-sensitive sites to generate short PSP loops (Figs. 4a and 4d).

**a**

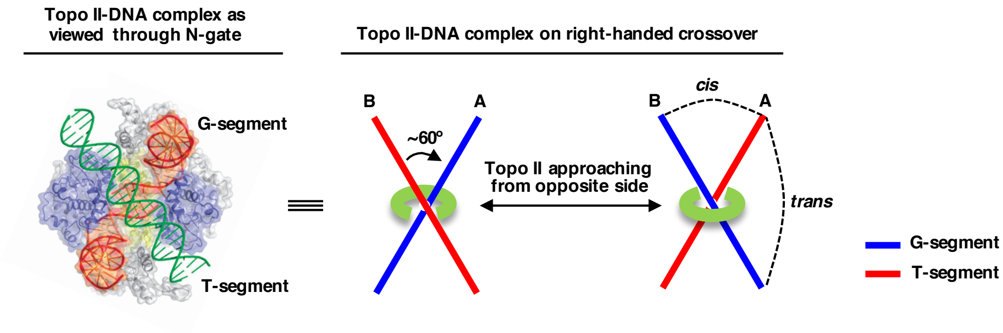


**b**

| Topo IIβ-DNA complex | | G/T contiguous (PSP) | | | | | | | | G/T segregated (DSP) | | | | | | | |
|---|---|---|---|---|---|---|---|---|---|---|---|---|---|---|---|---|---|
| | | 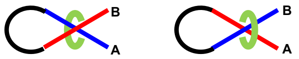 | | | | | | | | 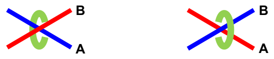 | | | | | | | |
| Type of complex | | 1 | | 2 | | 3 | | 4 | | 5 | | 6 | | 7 | | 8 | |
| G-segment | DSB or SSB | A | B | A | - | - | B | - | - | A | B | A | - | - | B | - | - |
| | Break resealed | - | - | - | B | A | - | A | B | - | - | - | B | A | - | A | B |
| T-segment | | B | A | B | A | B | A | B | A | B | A | B | A | B | A | B | A |
| 0.5 M NaCl treatment | P1 | AB | AB | AB | - | - | AB | - | - | A | B | A | - | - | B | - | - |
| | P2 | - | - | - | AB | AB | - | AB | AB | B | A | B | A, B | A, B | A | A, B | A, B |
| Toposite | Ts1 | AB | | | | | | | | | | | | | | | |
| | Ts2 | | | | | | | AB | | | | B | | A | | A, B | |
| | Ts3 | | | AB | | AB | | | | A, B | | A | | B | | | |

**Fig. S2** Configuration of DNA segments recognized by topo II and toposite assignments in eTIPa-seq. **a** Topological relationship between topo II and DNA segments in reaction intermediates. The complex shown on the left represents the time point when G-segment is bound at the DNA gate and T-segment is entering from the N gate [2]. A simplified scheme on the right depicts the handedness of crossover between G- and T-segments held in topo II-DNA complex, which is 'right-handed' by definition. Topo II molecule is drawn in a 'C' shape that can approach from either side of the crossover. Indicated on the right are two plausible modes of ligation between G- and T-segments. The *cis*-ligation occurs in the smaller angle of crossover and the *trans*-ligation occurs in the larger angle of crossover across topo II molecule (see Fig. S4d). **b** Combinations of G- and T-segments in the crossover. The two DNA segments at the crossover point is discriminated by 'A' and 'B' (represented 'AB' if contiguous). Using this distinction, the complex is classified into 8 types, which correspond to the mode of P1/P2 partitioning and toposite assignments. DSB, double strand breakage; SSB, single strand breakage.

### Critical factors in the eTIP procedure

In conventional immuno-capturing methods to separate the topo II-DNA cleaved complex, etoposide-treated cells are normally lysed by solutions containing SDS, a strong detergent, which denatures the enzyme instantly and dissociates all the DNA that are bound non-covalently. Thus, the fractionated topo II-DNA complex contains only DNA that are linked to the enzyme covalently (G-segment). In the present study, however, we lysed cells with a weaker detergent, sarkosyl (sodium lauroyl sarkosinate), to preserve the association of DNA interacting non-covalently with the enzyme, which may well include T-segments bound to the reaction intermediate. Although sarkosyl has been shown to dissociate histones effectively from chromatin DNA [4], it leaves RNA polymerase II bound to template DNA without inactivating the enzyme, as revealed by continued transcription even in the presence of 1% sarkosyl in nuclear run-on assays [5, 6]. Therefore, topo IIβ may also retain non-covalently bound DNA and a certain level of enzymatic activity in 1% sarkosyl. Since etoposide is diluted away after the cell lysis, the residual topo II activity may reseal the breaks in G-segment to some extent. When breaks are re-ligated completely, the G-segment might be dissociated from the topo IIβ immobilized on beads. However, this cannot be the case because G-segment should remain associated non-covalently with the enzyme until it is released by 0.5 M NaCl treatment together with T-segment in eTIPa-seq. This is a logical consequence derived from the existence of Ts2 toposites and also demonstrated by a model experiment described in the next section.

In the practical procedure, cell lysate in 1% sarkosyl was supplemented with 0.5 M CsCl to further remove materials interacting nonspecifically with topo IIβ. In the final step of eTIPa-seq, significant amounts of DNA are eluted in P2 fraction by 0.5 M NaCl treatment, implying that positively charged residues of topo IIβ interacting with phosphoryl groups of DNA are replaced by $Na^+$ ion but not by $Cs^+$ ion. This is consistent with the fact that the accumulation of alkali metal cations near the dsDNA phosphoryl groups varies inversely with its ionic size [7]. Thus, $Na^+$ is stronger than $Cs^+$ as an agent for eluting DNA fragments ionically bound to topo IIβ. We emphasize the fact that the DNA yield in P2 is highly dependent on etoposide (Fig. S1c). This is a strong implication that these DNA fragments are not just irrelevant ones but are genuine components of topo IIβ reaction intermediates.

**Evidence for the presence of T-segment in the topo IIβ-DNA complex**

We designed a model experiment in that tag-purified topo IIβ was immobilized on magnetic beads and supercoiled plasmid DNA was used as a substrate (Fig. S3a). We first confirmed the absence of contaminating topo I activity by a reaction without ATP, which showed no relaxation of supercoils (results not shown). The enzyme targets intramolecular crossovers in the circular substrate, cleaves one duplex (G-segment) to attach covalently to its ends, while holding the other (T-segment) non-covalently between protomers. These DNA segments are contiguous until the bound plasmid is cut by restriction enzymes to segregate insert and vector fragments. G/T-segments can be distinguished afterwards by high-salt treatment that releases the T-segment (Fig. S3a).

The standard topo II reaction converted all substrate to relaxed form (Ir), most part of which is found in unbound fraction (Fig. S3b, lane 3). In the presence of etoposide, the enzyme-bound fraction consisted of dsb products that are form III DNA (one cut) and smearing DNA (multiple cuts) in addition to form II DNA representing ssb product (lane 4). To mimic the cell lysis step of eTIP, concentrated sarkosyl and CsCl were added directly after the reaction with etoposide and then fractionated (lane 5). Changes of relative abundance are noticeable in topo IIβ adducts detected in the bound fraction: the smeared band showed a decrease in intensity and mobility but forms III and II bands were intensified. Moreover, the form Ir band newly appeared. These changes indicate that in the presence of sarkosyl/CsCl the enzyme-bound cleaved DNA undergoes significant religation processes resulting in decrease in multiple dsb (smear), transformation from dsb (form III) to ssb (form II) and from ssb to no break (form Ir). It is worth noting that all these DNA bands are still bound to the enzyme including Ir, which should be held by non-covalent interaction. These associations are quite stable against the wash with restriction buffer (lane 6). When bound DNA was cut with restriction enzymes to separate vector and insert fragments, only a portion of DNA was released into solution (lane 7). The ratio of the band intensity in U fraction should reflect the statistical abundance of crossovers in vector (2.7 kb) and insert (1.4 kb) portions. The enzyme-bound DNA was then treated with high-salt solution containing 0.5 M NaCl (lane 8). A significant amount of DNA was released into solution indicating that this DNA had been associated with the enzyme non-covalently. However, if the high-salt treatment was applied prior to restriction cutting, no DNA was detected in U fraction, indicating that the whole plasmid molecule is tethered to the enzyme covalently (forms II and III) or topologically (form Ir) (lane 9). Interpretations for the results are illustrated in Fig. S3c. Important finding from this experiment is that DNA fragments released by high-salt treatment are mostly accounted for by T-segments with crossovers between vector and insert portions (*xIV* and *xVI*) because all the other fragments (RVV, RxVV, RII, RxII) are derived from the bound DNA with no break (R), which should be only a minor component of released DNA.

Taken together, we show here unequivocally that in the eTIP procedure the enzyme-DNA complex after sarkosyl/CsCl treatment retains a DNA segment interacting non-covalently with the enzyme, which is released by high-salt treatment and recovered in P2 fraction. Considering the specificity of immuno-capturing and the fact that P2 DNA yield is highly dependent on etoposide, it is a logical consequence that the DNA in P2 fraction contains the T-segment released from topo IIβ reaction intermediates. These experiments also demonstrate the presence of G-segment held by the enzyme even after resealing reaction in the absence of etoposide. After all, the unique properties of eTIP procedure is attributable to the particularity of sarkosyl as a weak denaturant that preserves the basic structure and partial enzymatic activity of topo IIβ.

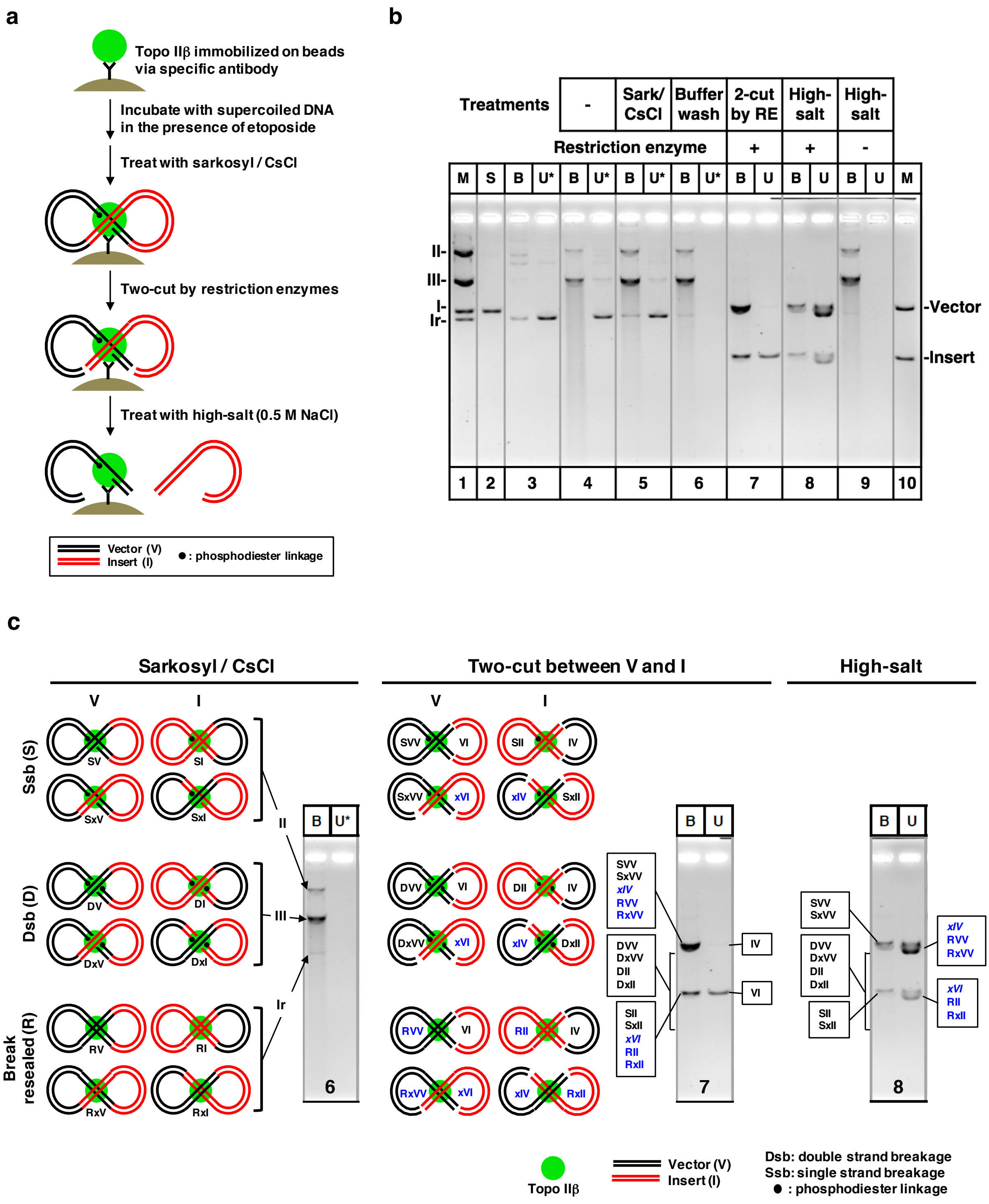


**Fig. S3** An *in vitro* experiment mimicking the *in vivo* eTIP procedure. **a** Outline of the model experiment. Flag-tagged rat topo IIβ expressed in cultured cells was immobilized on magnetic beads through anti-Flag antibody and used for the experiment. Supercoiled plasmid vector containing *E. coli* DNA insert was used as a substrate (not drawn in proportion). For simplicity, the enzyme-bound form of the plasmid depicted here has a crossover between vector and insert portions although other forms are possible as shown in panel c. **b** Fractionation of reaction products generated from the substrate DNA (labeled 'S'). In lanes 3-9, DNA bound to the enzyme (B) and unbound (U) were fractionated by magnetic separation and electrophoresed in an agarose gel containing ethidium bromide. DNA markers shown in lane 1 (M) are various forms of the same plasmid: supercoiled (I), relaxed closed circle (Ir), nicked circle (II), and linear form (III). The substrate DNA was allowed to react with topo IIβ on beads in the absence of etoposide (lane 3) or in the presence of etoposide (lane 4), to which sarkosyl/CsCl was added directly and fractionated (lane 5). Hereafter, the bead-bound enzyme-DNA complex was treated consecutively as follows: washing with restriction buffer (lane 6), cutting with *Eco* RI and *Sal* I (lane 7), eluted with high-salt solution (lane 8), and high-salt elution without restriction cutting (lane 9). The loaded sample amounts in B and U are comparable but only 1/7.5 amount is loaded in U*. Shown in lane 10 are the vector and insert fragments as a marker. **c** Interpretation of the results illustrated schematically. Gel patterns from steps 6, 7, and 8 are replicated here. All possible forms of substrate DNA bound to topo IIβ after sarkosyl/CsCl treatment are depicted on the left panel. The three cleavage states of G-segment are indicated on the left margin (S, D, and R). Cases where G-segment is formed within the vector portion (labeled 'V') are placed on the left column and those within the insert portion (labeled 'I') are placed on the right column. Also discriminated in the figure is the position of crossover. Cases where crossover is formed within vector (or insert) are placed in the upper row and those between vector and insert are placed in the lower row. The enzyme-bound plasmids are named systematically using the letters for abbreviation. The "x" stands for the presence of crossovers between vector and insert. Fragments after restriction cutting are depicted on the middle panel. The fragments grouped in boxes correspond to the gel bands indicated by connecting lines: upper band, vector; lower band, insert. The fragments written in blue are associated with the enzyme non-covalently and released into unbound fraction after high-salt treatment (right panel). Among these fragments, the ones set in italic type (*xIV* and *xVI*) are presumable T-segments. As the dsb fragments vary in size depending on the distance between the break point and restriction sites, they distribute as a smear band shown by brackets.

**Mapping G/T-segment pairs bound to topo IIβ (eTIPb-seq)**

To obtain direct evidence for PSP and DSP, we devised a new procedure illustrated step by step in Fig. S4a, to which a ligation step between G- and T-segments was included to measure their distance in genome. Briefly, the IP complex captured on magnetic beads are processed as follows. After an adaptor oligonucleotide is attached to the ends of enzyme-bound DNA fragments, the adaptor ends are ligated to generate chimeras between G- and T-segments (these steps were done *in situ* on magnetic beads). Only chimeric fragments containing an internal adaptor dimer were selected in the following steps and sequenced from both ends by a paired-end sequencing protocol. The resulting sequence reads still contain a considerable level of noise due to contaminating DNA fragments, which were selected out as follows. Reads were first mapped to the rat reference genome (rn4) by BWA and read pairs that are uniquely mapped on both ends were selected. Next, PCR duplicates and short fragments were removed. Chimeras were then sorted into intra-chromosomal and inter-chromosomal chimera. They were classified into 4 groups based on the combination of read orientation on paired ends (Fig. S4b). According to the mechanistic rationale, both reads on chimera should overlap or sit very close to toposites detected by eTIPa-seq (both reads should be closer than 1 kb to toposites). All chimeras that fail to meet the criteria were discarded at the final step.

Read sequences should match the genomic sequence of either Watson strand (forward=F) or Crick strand (reverse=R). In principle, therefore, intra-chromosomal chimeras can be designated RF, FR, FF, and RR in the order of upstream-downstream. When their abundance was compared, the ratio was extremely skewed, RF being by far the dominant orientation (Fig. S4c). This can be explained by the mechanisms described below (Fig. S4d). Two types of loop structure, ‘plectonemic’ and ‘toroidal’, can be formed when topo IIβ bound to right-handed crossover configuration. Since RF is the only combination left for ligation if the loop structure remains intact, its dominance may suggest that a significant proportion of intra-chromosomal chimera originates from the crossover at the base of loops that survived shearing (shorter than 3 kb). Analysis of the relationship between chimera length and read orientations clarified the whole picture (see Fig. 3 of main text). As topo II action on toroidal loop is ineffective compared to plectonemic loop when turnover rate of relaxation is considered [2], plectonemic configuration combined with *cis*-ligation is more feasible to occur than the other way around. As for the ligation mode, *cis*-ligation should be more efficient than *trans*-ligation, in which case the ligation may be interfered by the steric hindrance imposed by topo IIβ molecule (Fig. S2a). In the case of inter-chromosomal chimera, the two DNA duplexes interacting with topo II take two possible orientations (Fig. S4d). All the read orientations occur with equal frequency because difference in the efficiency between *cis*- and *trans*-ligations will be averaged out.

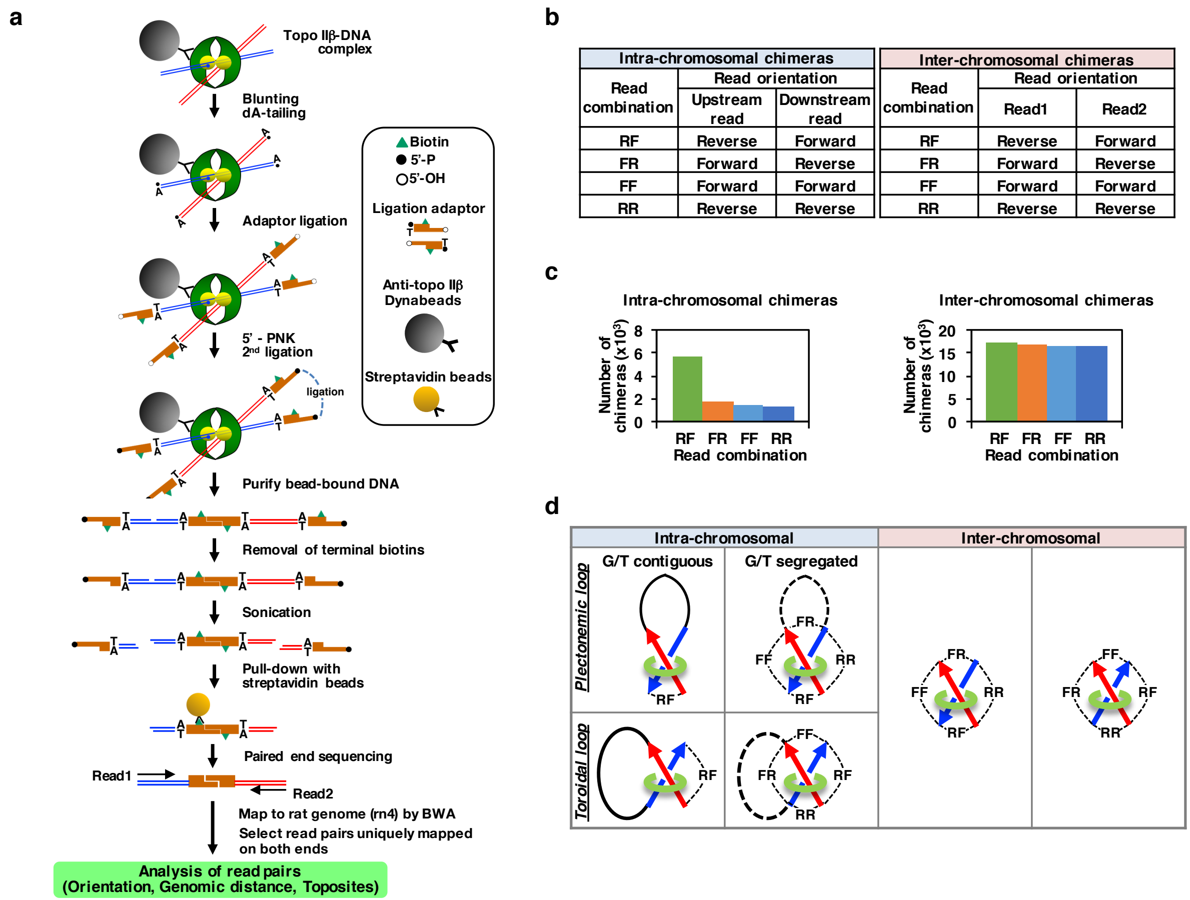


**Fig. S4** Outlines of eTIPb-seq procedure and analysis of chimeras. **a** A flowchart of eTIPb-seq procedure. The topo IIβ-DNA complex was first immuno-captured on magnetic beads. The ligation reactions were conducted on the beads. For detailed description on the following steps, see Methods. **b** Orientation of paired reads. Note that read-pairs mapped on the same chromosome can be located either upstream or downstream with respect to the Watson strand (+strand) of the reference sequence. **c** Number of resulting chimeras as a function of read orientation. **d** Possible configuration of the two DNA segments in the topo IIβ-DNA complex. Ligatable ends are tied by dotted lines and indicated by read orientations at the chimera ends. All crossovers are right-handed that is a preferred target of topo IIβ. Upstream and downstream DNA segments in intra-chromosomal crossovers are drawn by red and blue arrows, respectively.

### Logical analysis for the chimera identities

To facilitate the interpretation of the results on chimera analysis (Fig. 3), we made an exhaustive list of possible chimeras and compared with the experimental data (Fig. S5). Ligation products (chimeras) are categorized into 3 groups: auto-ligation, G-T chimera, and inter-complex chimera (Fig. S5a). Among these, G-T chimeras are most informative ligation products in the eTIPb-seq analysis. Ts1-Ts1, Ts2-Ts2, Ts3-Ts3 chimeras for G/T contiguous case correspond to Ts1/PSP, Ts2/PSP, and Ts3/PSP chimeras in Fig. 3e. Ts2-Ts2 and Ts3-Ts3 chimeras for G/T segregated case correspond to Ts2/DSP and Ts3/DSP chimeras in Fig. 3e. The G/T segregated DNA fragments may produce auto-ligation products (read distance<3 kb) with RF orientation, which might contaminate Ts2/PSP, and Ts3/PSP chimeras. However, auto-ligation products are likely to be formed by the *trans*-ligation mechanism, which is less efficient than *cis*-ligation and thus the product should be negligible in number compared to G-T chimeras.

Since the ligation between adaptors was performed on magnetic beads, the formation of "inter-complex chimera" between neighboring topo IIβ molecules is somewhat inevitable. As no preference in read orientation is expected for the inter-complex chimera, its basal levels are predictable as a background in the table of chimera abundance (Fig. S5b). For the long distance region (2 Mb<), it is obvious that all chimeras detected are inter-complex chimeras, reflecting a background noise. Similarly, Ts1-Ts1 chimera in the 3 kb-2 Mb region is mostly accounted for by inter-complex chimeras.

**a**

| | | G/T contiguous (Intact loop) | | | | | | | | G/T segregated (Disrupted loop) | | | | | | | |
|---|---|---|---|---|---|---|---|---|---|---|---|---|---|---|---|---|---|
| Topo IIb-DNA complex | | 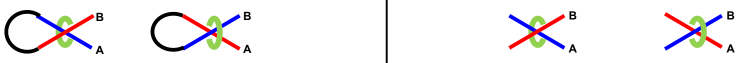 | | | | | | | | | | | | | | | |
| Type of complex | | 1 | | 2 | | 3 | | 4 | | 5 | | 6 | | 7 | | 8 | |
| G-segment | | A | B | A | B | A | B | A | B | A | B | A | B | A | B | A | B |
| Strand break | | + | + | + | - | - | + | - | - | + | + | + | - | - | + | - | - |
| Toposite | | Ts1 | Ts1 | Ts3 | Ts3 | Ts3 | Ts3 | Ts2 | Ts2 | Ts3 | Ts3 | Ts3 | Ts2 | Ts2 | Ts3 | Ts2 | Ts2 |
| Ligation | Auto-ligation | | | | | | | | | Ts3-Ts3 | Ts3-Ts3 | Ts3-Ts3 | Ts2-Ts2 | Ts2-Ts2 | Ts3-Ts3 | Ts2-Ts2 | Ts2-Ts2 |
| | | | | | | | | | | Read distance < 3 kb Read orientation : RF | | | | | | | |
| | G-T chimera | Ts1-Ts1 | | Ts3-Ts3 | | Ts3-Ts3 | | Ts2-Ts2 | | Ts3-Ts3 | | Ts2-Ts3 | | Ts2-Ts3 | | Ts2-Ts2 | |
| | | Read distance < 3 kb Read orientation : RF | | | | | | | | Read distance > 3 kb Read orientation : RF, FR, FF, RR | | | | | | | |
| | Inter-complex chimera | Ts1-Ts1, Ts1-Ts2, Ts1-Ts3, Ts2-Ts2, Ts2-Ts3, Ts3-Ts3 | | | | | | | | | | | | | | | |
| | | Read distance: any distance Read orientation : RF, FR, FF, RR | | | | | | | | | | | | | | | |

**b**

| Read distance | Read orientation | Ts1-Ts1 | Ts1-Ts2 | Ts1-ts3 | Ts2-Ts2 | Ts2-Ts3 | Ts3-Ts3 |
|---|---|---|---|---|---|---|---|
| 0 - 3 kb | RF | 3,432 | 90 | 25 | 446 | 18 | 291 |
| | FR | 300 | 10 | 17 | 47 | 5 | 35 |
| | FF | 257 | 0 | 0 | 24 | 0 | 5 |
| | RR | 236 | 0 | 0 | 16 | 0 | 4 |
| 3 kb - 2 Mb | RF | 66 | 31 | 50 | 86 | 83 | 155 |
| | FR | 57 | 32 | 42 | 76 | 81 | 144 |
| | FF | 43 | 9 | 14 | 15 | 21 | 19 |
| | RR | 46 | 8 | 14 | 15 | 17 | 18 |
| 2 Mb < | RF | 729 | 187 | 36 | 19 | 2 | 4 |
| | FR | 724 | 159 | 33 | 21 | 10 | 4 |
| | FF | 740 | 166 | 52 | 23 | 8 | 0 |
| | RR | 734 | 177 | 40 | 13 | 4 | 3 |
| Total chimera number | | 7,364 | 869 | 323 | 801 | 249 | 682 |

**Fig. S5** Interrelationship among the factors that define the chimeric species. **a** List of possible chimeras. Upper part of the table is a simplified version of Fig. S2b. G-T chimeras (Ts1-Ts1, Ts2-Ts2, Ts3-Ts3) are most meaningful ligation products in the present study. Auto-ligation products originate from end-to-end ligation of segment A or B that are shorter than 3 kb. Inter-complex chimera stands for the chimeras created from ligations between DNA fragments bound to different topo IIβ molecules. They are composed of any combination of toposites, any lengths and any read orientations. **b** List of actual numbers of chimera detected experimentally. Significant chimeras are highlighted. The Ts2-Ts3 chimeras are shaded gray because this toposite combination is statistically insignificant (Fig. 3c).

**Supporting evidence for the presence of DSP sites**

Most remarkable finding of the present study is that DSP occurs at the DNA crossovers facilitated by pairing of homologous DNA segments located at a long distance. Since this discovery is almost unprecedented, some criticism has been raised. Etoposide-induced dsb mediated by topo IIβ might be repaired by microhomology-mediated end-joining (MMEJ) or by other homologous end-joining mechanisms. During this process, two distally located homologous sites would be somehow brought into proximity and processed into chimeric ligation products, namely DSP chimeras. However, this is unlikely to occur for number of reasons. i) Topo IIβ molecule should be removed from dsb ends before the start of repair end-joining process, which requires multiple protein factors. There should be little possibility that all these factors are associated with topo IIβ-DNA complex after cell lysis and execute the expected reaction. ii) MMEJ usually operates between short homologous DNA segments in the vicinity of dsb site [8] that are not positioned kilobases apart from each other like in DSP chimeras. iii) The DSP chimera is derived from the ligation between two DNA fragments associated with topo IIβ. The ligation occurs between artificial adaptors attached to the ends of randomly sheared DNA. iv) MMEJ-mediated ligation should never bring about the biased read orientations observed in DSP chimeras (Fig. 6b).

For further validation of DSP, we designed a model experiment described below. To rule out the possibility that DSP is an artifact caused by some particularity of etoposide, the cleaved complex was induced by another agent, namely $H_2O_2$, an entirely different chemical substance compared to etoposide. Hydrogen peroxide had been shown to produce a topo II-DNA covalent complex indistinguishable from the one formed by etoposide [9]. $H_2O_2$-treated cells were lysed with sarkosyl/CsCl according to the eTIP procedure and immunoprecipitated with anti-topo IIβ antibody (see Methods for details). All the following reactions were conducted on the bead surface. As in eTIP, DNA fragments derived from the crossover between distant genomic sites should be bound to topo IIβ as G- and T-segments

(Fig. S6a). Ligation between these fragments was done differently from the eTIPb-seq procedure. Instead of adaptor-mediated ligation, DNA fragments were directly ligated after cutting the bead-bound DNA with *Mbo* I. We selected three Ts3/DSP chimeras from the list (Table S5) and set PCR primers within a 2 kb-region, upstream and downstream, harboring the sequence read. The primer pairs should uniquely amplify the target within 2 kb-region or within predicted ligation products between the regions. Since the chimera ends are homologous, it was rather difficult to find unique sequences for primers (Table S8).

We examined two criteria to see whether the selected Ts3/DSP chimeras truly reflect the association of two distant loci. First, if this is the case both upstream (U) and downstream (D) regions should be associated with the immunoselected topo IIβ. When appropriate PCR primers were used, both U and D regions were found in the immunoprecipitate (IP) (Fig. S6b). Under these conditions, however, no enrichment was observed although the negative control region (Mybpc2) was clearly depleted in IP. More definitive test would be to demonstrate the ligation-dependent PCR amplification between the two regions. *Mbo* I sites were first identified in U/D regions and from the probable combinations of ligation between the sites, a minimal product length for certain primer pairs can be predicted (Fig. S6a). For the same set of chimeras, PCR products were compared between template DNAs isolated before and after ligation (Fig. S6c). In all cases multiple bands were detected only when the ligated IP DNA was used as a template and their smallest size roughly coincided with the predicted ones. The longer products may arise from insufficient cutting or polymorphic nature of *Mbo* I sites in the genome. Presence of the same size band in all three lanes may indicate some off-target amplifications caused by miss-annealed primers. These results suggest that DSP sites detected by eTIPb-seq reflect genuine association of distant genomic sites that are targeted by topo IIβ.

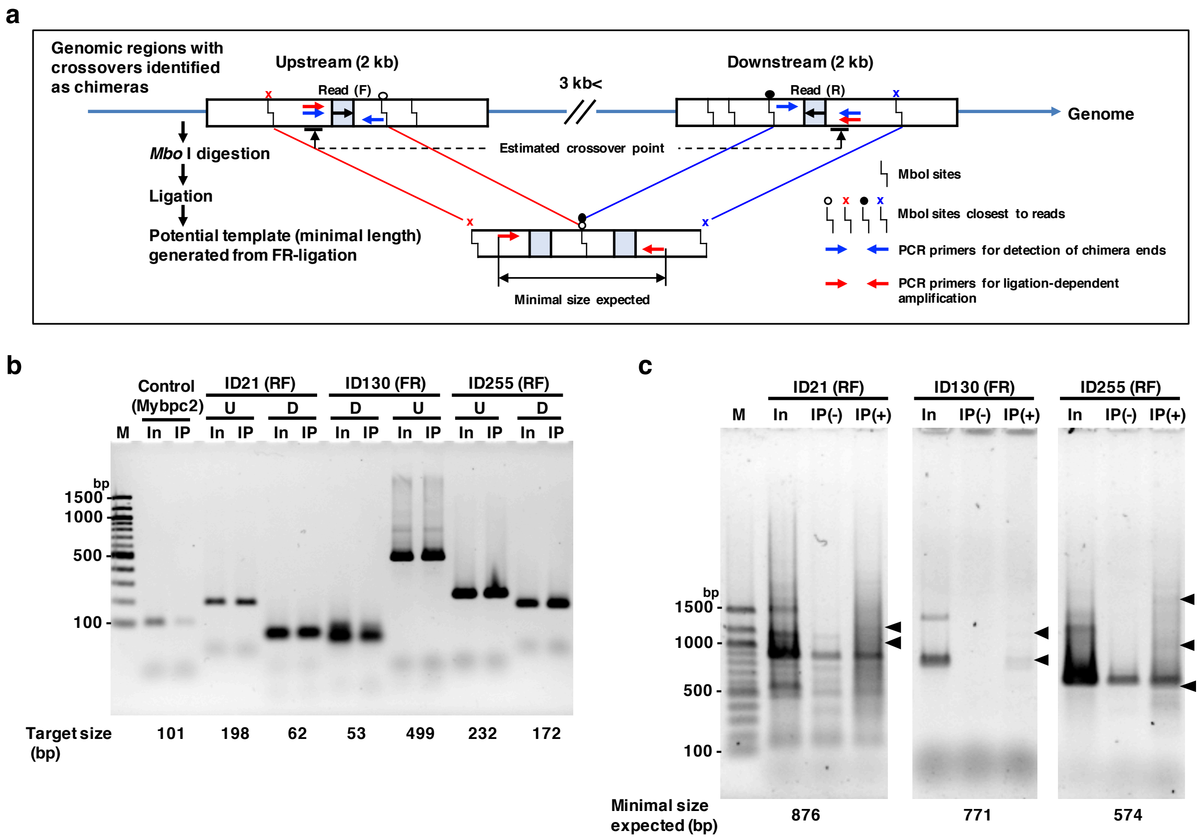


**Fig. S6** A model experiment that substantiates the DSP. **a** Schematic representation of the strategy. Shown here as an example is a Ts3/DSP chimera with FR orientation whose upstream (U) and downstream (D) regions form a crossover and associated with topo IIβ immobilized on magnetic beads (Table S5). Although exact position of crossover is unknown, estimated crossover point is indicated by bars tied with dotted line. *Mbo* I sites are detected on the rat reference sequence (marked by kinked line with symbols). DNA fragments between *Mbo* I sites adjacent to the read position remain on beads when DNA fragments bound to topo IIβ were cut with *Mbo* I and free fragments were washed out. Subsequent ligation occurs between the *Mbo* I sites marked by circles (open and filled) to generate FR orientation and should result in the potential template depicted below. Two kinds of primer pairs were designed by closely examining the sequences for the region of interest: one kind detects the U and D regions (blue arrows) and the other detects the predicted ligation product (red arrows). Candidate primer sequences were scrutinized by homology search and by an *in silico* PCR program (UCSC genome browser). See Table S8 for primer sequences used. **b** Presence of both U and D regions in the immunoprecipitate (IP). Chimeras selected for the analysis are indicated on top with ID numbers and read orientations (in parenthesis). M, ladder marker (numbers in bp); In, input DNA for IP; IP, bead-bound DNA (*Mbo* I-cut). Predicted target size is shown beneath the gel pattern. For control amplification, a primer pair was set in Mybpc2 gene. **c** Evidence for the close positioning of U/D regions in the IP DNA. In, input

DNA for IP; IP(-), bead-bound DNA before ligation; IP(+), bead-bound DNA after ligation. PCR products identified uniquely in IP(+) are marked by arrowheads. Note that the size of lowest bands roughly coincides the expected size of minimal length product (shown on bottom).

**Other genome-wide mapping for correlative analyses**

We performed additional genome-wide analyses together with topo IIβ-targeted sites. Most attention was given to the binding sites of hnRNP U/SAF-A/SP120 as determined by ChIP-seq.

While any DNA, with a certain sequence preference, can be a substrate for topo II *in vitro*, accessibility of the enzyme to DNA *in vivo* is restricted significantly by bound chromatin proteins. Therefore, FAIRE-seq assays were employed to probe the local chromatin accessibility to evaluate potential topo IIβ targets.

These mapping results were displayed on custom tracks of the UCSC genome browser together with RNA-seq signals representing levels of transcription. Using this display, a number of correlative analyses were performed as described in the text.

### 2. Other supplementary figures

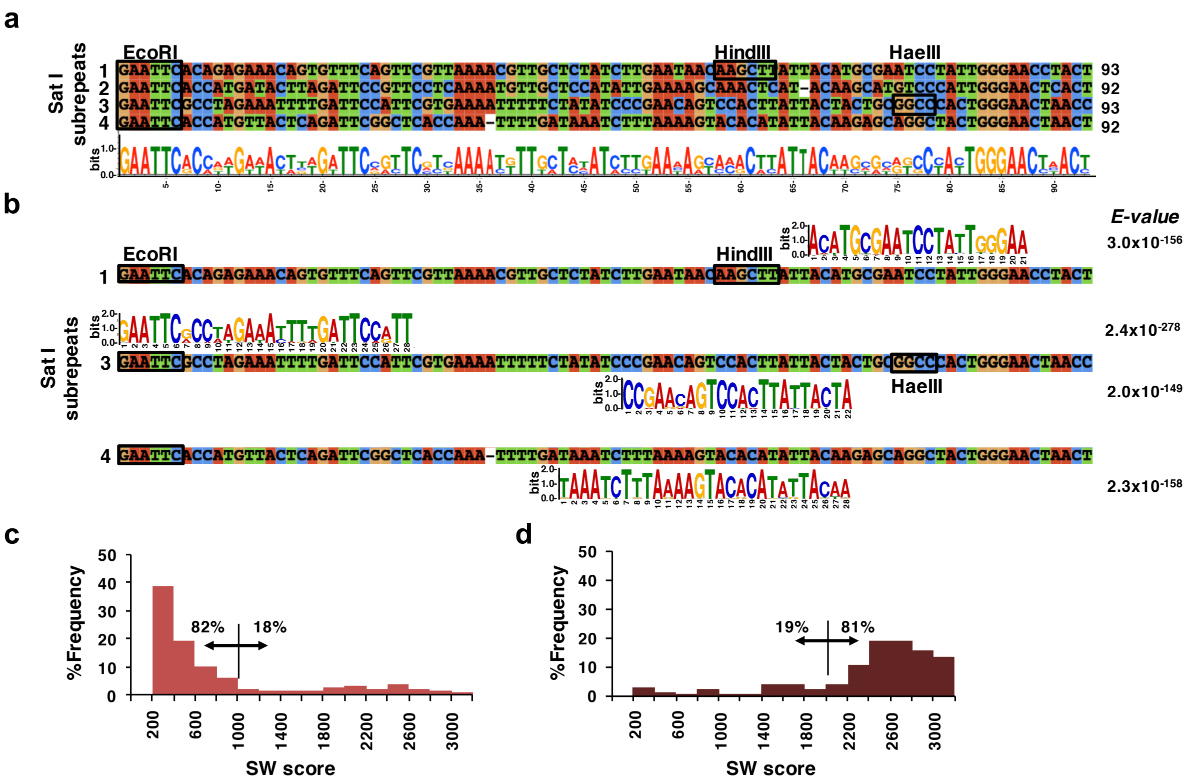


**Fig. S7** Strong correlation between satellite I (Sat I) repeats and the Ts3 toposite. **a** Sat I subrepeat structure. Rat Sat I is composed of 4 subrepeats of 92-93 bp. The canonical Sat I sequence of 370 bp [10] was cut into 4 fragments at the conserved EcoRI sites and subjected to multiple alignment analysis with Clustal X program and the result is shown with the sequence logo. Conserved restriction sites are indicated by boxed recognition sequences. HindIII and HaeIII sites are less conserved compared to EcoRI site. **b** Search for a consensus sequence in Ts3 toposites. Eight-hundred randomly selected Ts3 site sequences were subjected to analysis with MEME zoops program (version 4.11.3). A number of segments homologous to Sat I sequence with high statistical significance were detected. Matched positions are indicated by the Ts3-consensus sequence expressed in logo on the subrepeat sequence. The consensus sequence detected in plus strand is shown above and that in minus strand is shown beneath. **c** Histogram of SW scores for Sat I repeats obtained from UCSC repeat masker track data (n=1,884). **d** Histogram of SW scores for Sat I repeats that overlap with Ts3 toposites (n=143).

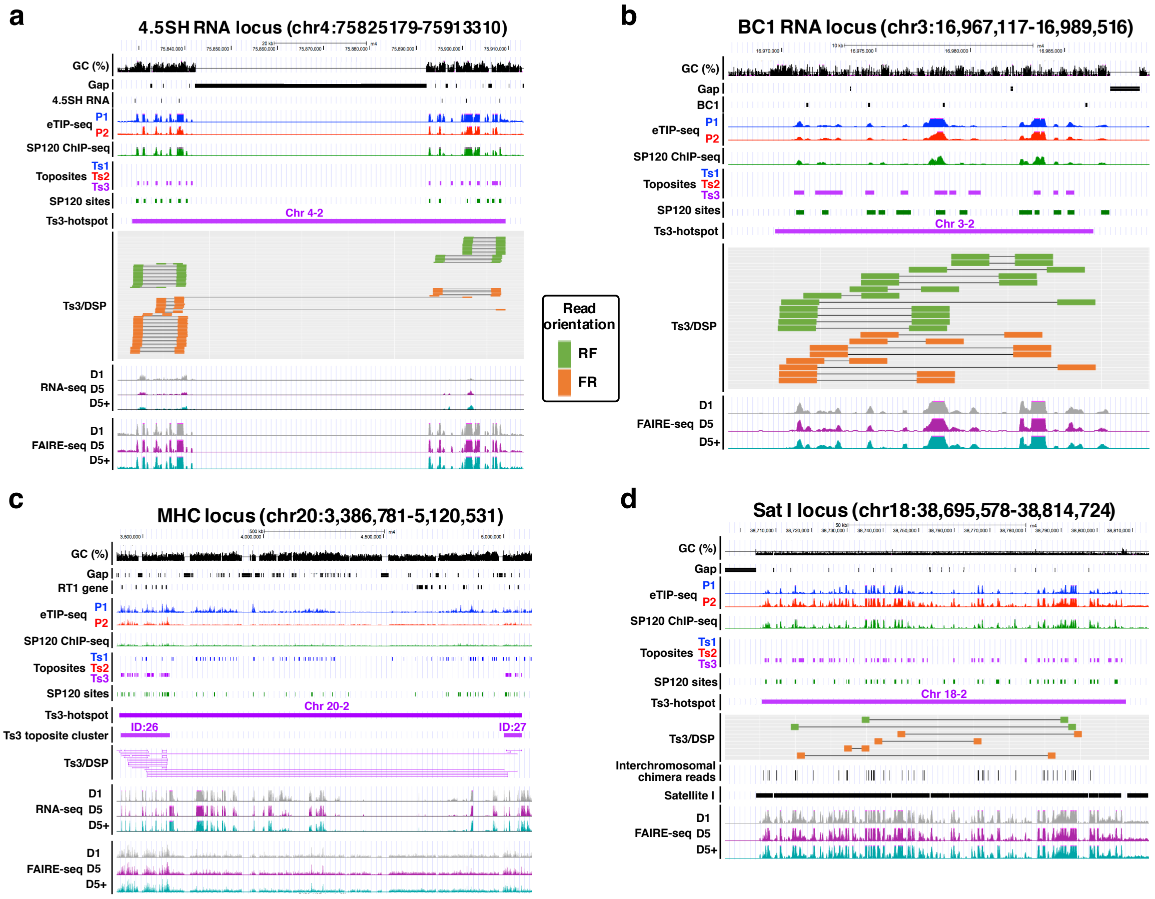


**Fig. S8** Ts3-hotspots that are worth featuring. **a** The hotspot Chr4-2 corresponds to a noncoding RNA gene cluster (4.5SH RNA) which is unique to myomorpha, a large group in rodents including rats, mice, and hamsters. The rat 4.5SH RNA (~94 bp) is a part of a 5.3 kb unit that is transcribed by RNA pol III, several hundred copies of which are arranged as tandem repeats on this locus of rat chromosome 4 [11]. In the rn4 browser view, the Ts3-hotspot is interrupted by a long sequence gap, whose position is very likely to be occupied by the same repeats flanking the gap and thus whole length may exceed 1 Mb. **b** The hotspot Chr3-2 also overlaps with a noncoding RNA cluster (BC1 RNA). This locus may have been formulated by post-reverse-transcriptional insertion of the RNA pol III transcript of BC1 RNA gene, which is located on rat Chr5 and expressed specifically in brain tissue [12]. **c** The hotspot Chr20-2 is composed of two Ts3 toposite clusters (ID:26 and ID:27) and resides within the rat MHC gene cluster. Some Ts3/DSPs connecting these clusters are among the longest that exceed 1 Mb. Detailed analysis of this region is given in Fig. S9. **d** The hotspot Chr18-2 is a Ts3-hotspot that encompasses about 100 kb of pericentromeric region adjacent to the p-q boundary of the cytoband. All the Ts3/DSP reads in this region (either RF or FR orientation) coincide with the positions of Ts3 toposites, SP120 sites, FAIRE-seq peaks and Sat I sequences with high SW score. Similar Sat I clusters are located in the pericentromeric region of Chr13, Chr17 and Chr18, as well. Therefore, Sat I repeats are preferential sites for DSP as well as PSP interactions (Fig. 4d). This region is also enriched with inter-chromosomal chimeras (see Discussion).

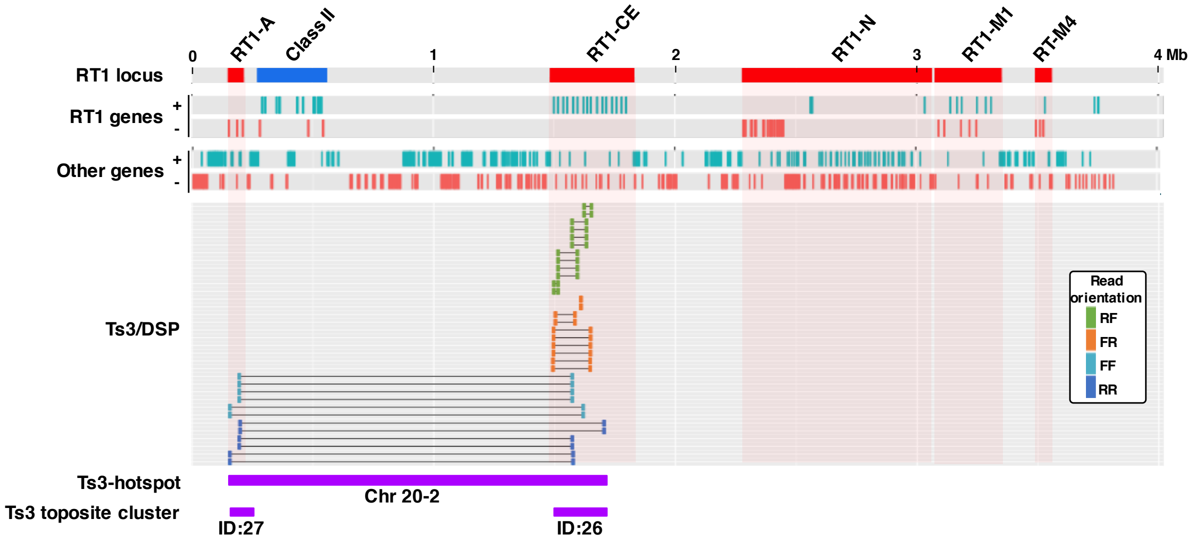


**Fig. S9** Reconstruction of the Ts3-hotspot Chr 20-2 on a complete sequence of rat MHC gene cluster. Using the published sequence [13], we improved the mapping accuracy of toposites and Ts3/DSPs on the reference sequence (rn4). As shown here (note that sequence direction is reversed), the rat MHC class Ia genes, termed RT1, are present in two gene clusters (designated RT1-A and RT1-CE). Reads belonging to Ts3/DSPs detected in this RT1 locus are all confined within the two Ts3 toposite clusters. In addition to intra-domain chimeras, cluster ID:26 contains long chimeras linking to the neighboring cluster ID:27. Read orientations of intra-domain chimeras are all RF or FR, whereas inter-domain chimeras are all RR or FF. This must be related to the fact that, in rodents, the RT1-CE domain is partially duplicated and inserted inversely to generate the RT1-A domain [13], which is consistent with the fact that RT1 genes are coded by opposite strands in these domains. The connection between the read orientation and the coding strand observed here is an additional strong evidence for the rule shown in Fig. 6a. In contrast to RT1-A/CE genes, other RT1 genes and MHC class II genes are devoid of Ts3/DSP.

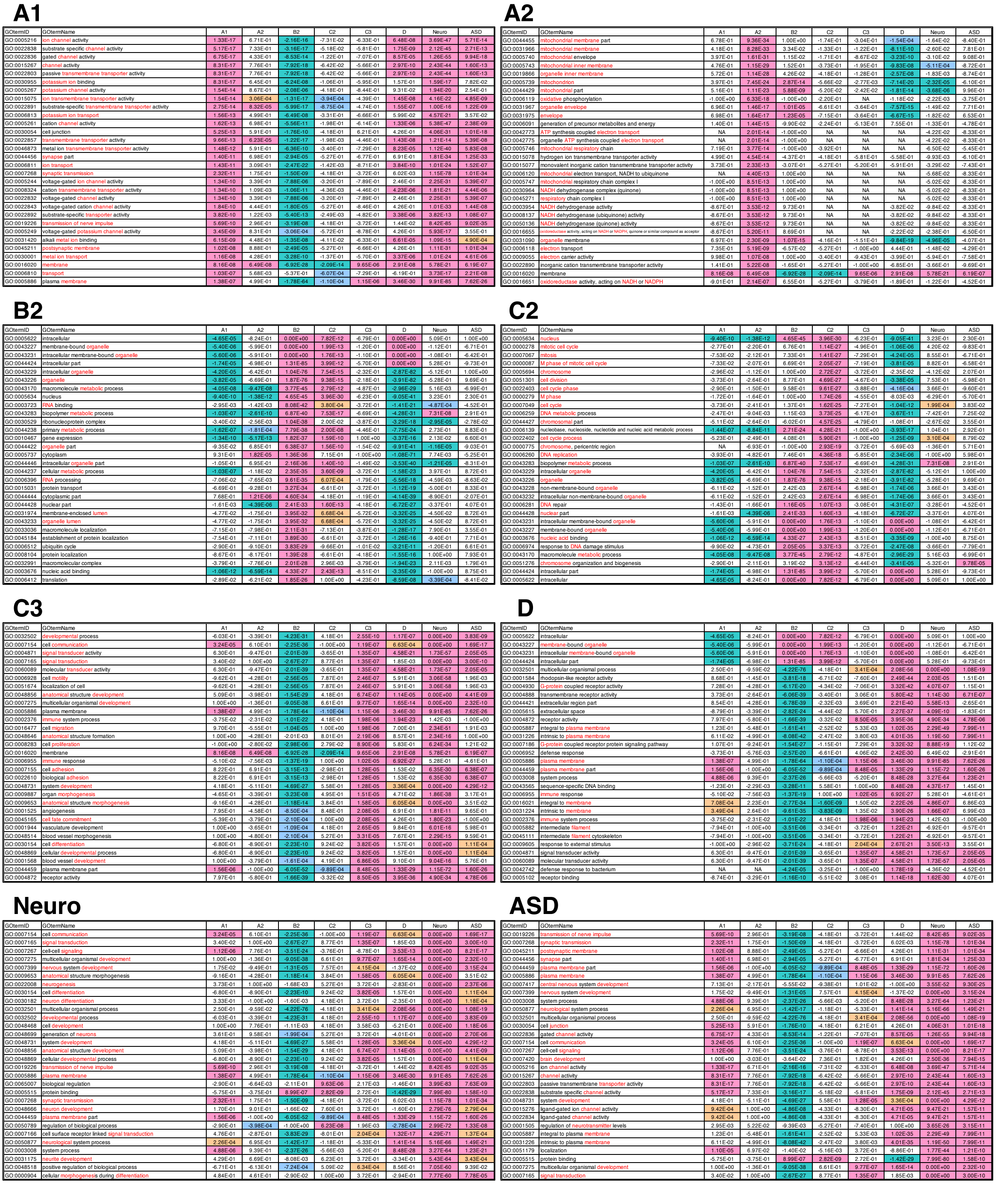

**Fig. S10** Gene ontology (GO) analysis of expression groups. This figure is a thumbnail representation of Table S7 (30 rows from the top are replicated here). GO term names are sorted for each group by arranging in descending order of significance. Words in the term name that characterize the functional aspect are highlighted in red. Highly enriched terms ($P<10^{-4}$) are painted in magenta and highly depleted ones, labeled with minus sign, are in blue-green. ‘NA’ designates ‘not applicable’ due to missing data.

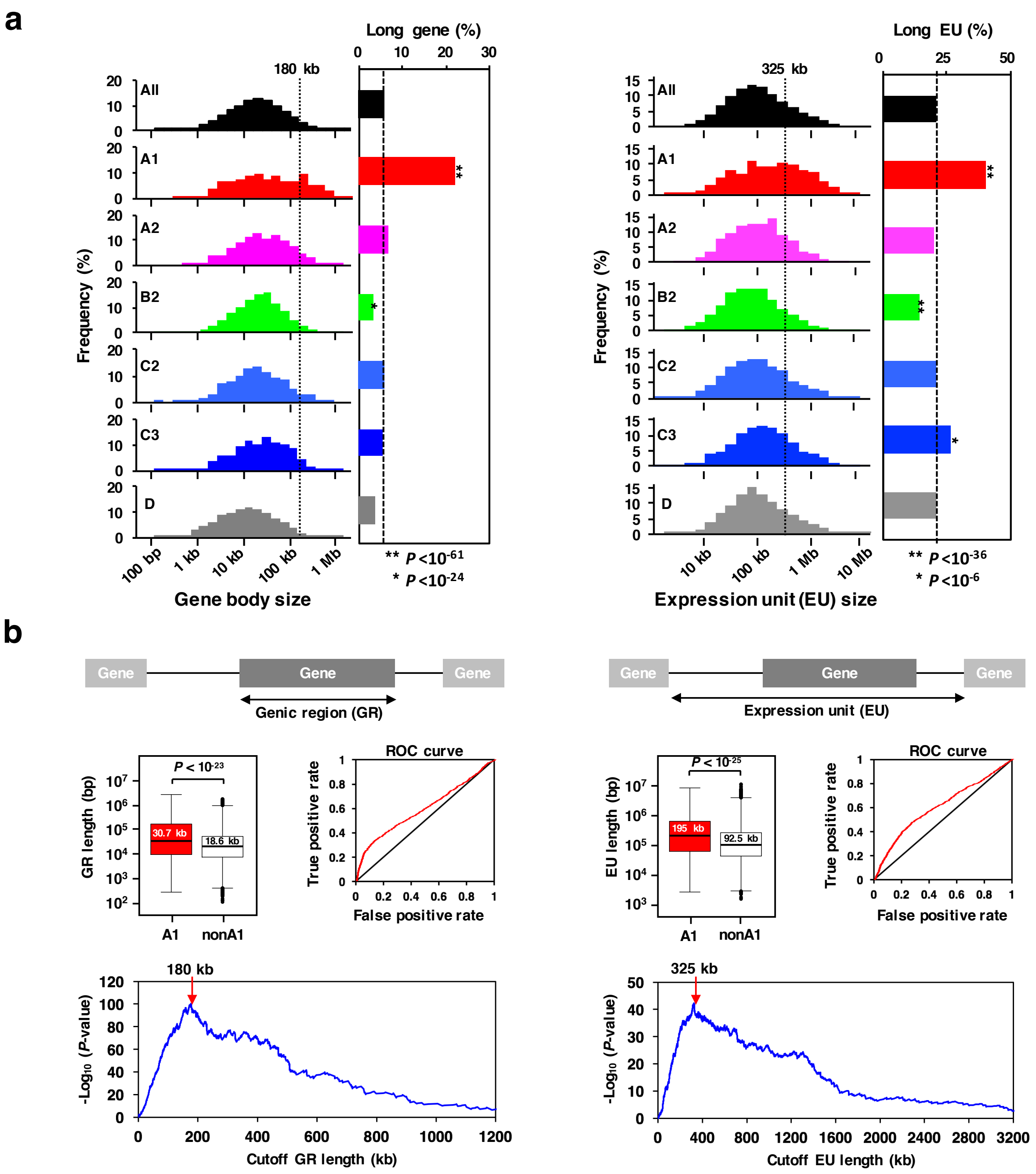


**Fig. S11** A long-vs-short binomial clustering analysis based on the threshold determined by ROC method. **a** Histograms for gene length and length of expression unit (EU). Relative abundance of the ‘long’ group for each expression group was calculated by applying the ROC-determined threshold (shown in b) and plotted in the bar graph. The statistical significance against ungrouped genes revealed that both A1 and C3 gene groups are enriched with long EU. Conversely, the house-keeping gene group (B2) is underrepresented in long EU. **b** ROC (Receiver Operating Characteristic) analysis to determine the threshold length between ‘long’ and ‘short’. Using exRefSeq data, the length distribution of genic region (GR) and expression unit (EU) with respect to A1 vs nonA1 genes was calculated and box-plotted. Median values are shown in the box and p-values are from Mann-Whitney test. ROC analysis was performed between the groups of A1 and nonA1 genes by the R library ROCR. The ROC curves presented are made between true positive rate (long A1/total A1) and false positive rate (long nonA1/total nonA1) at the varying threshold lengths. Significance levels of threshold values were estimated by chi-squared test and the resulting p-values are plotted against the cutoff length (bottom graph). The cutoff lengths with minimum p-value (red arrows) were chosen as the threshold length.

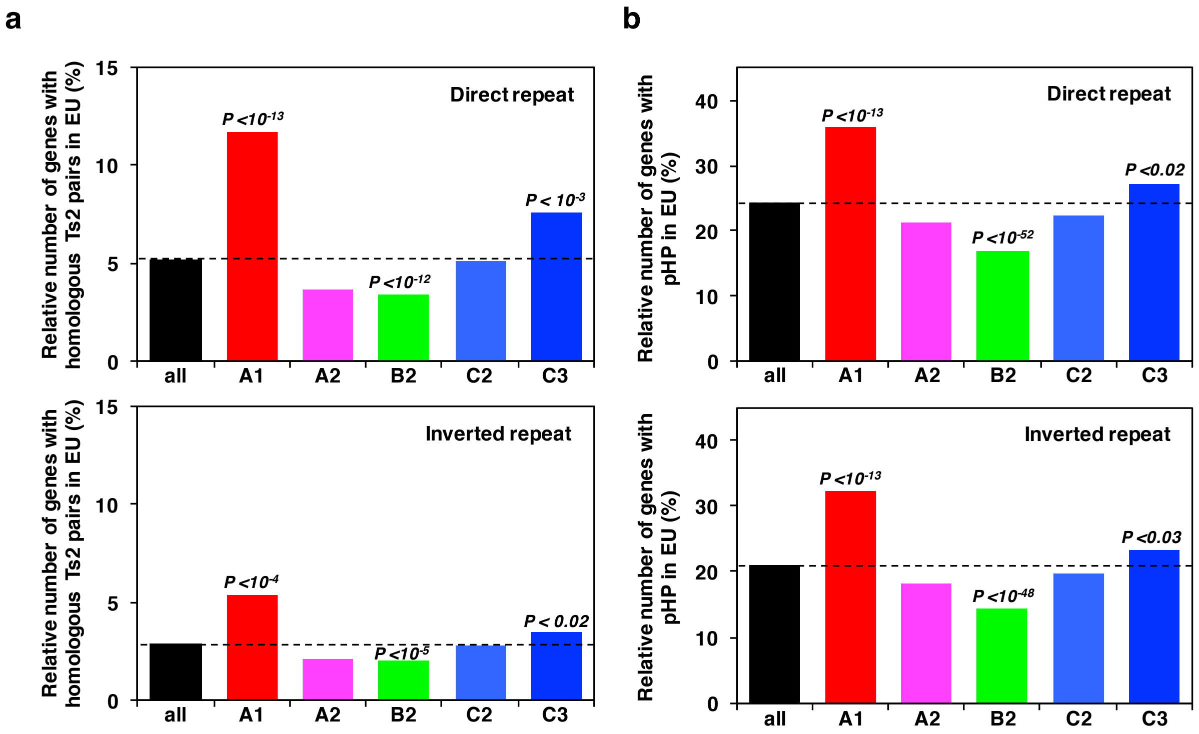


**Fig. S12** Relative abundance of genes containing homologous pairs: difference between experimental results and predicted results. **a** Homologous pairs searched among experimentally determined Ts2 sites. The analytical procedure used here is the same as described for Fig. 8f except that SW scores were calculated separately for direct and inverted repeats. The result shows that direct repeat is much abundant than inverted repeat. **b** Predicted homologous pairs (pHP). The result shows basically no difference between direct and inverted repeats.

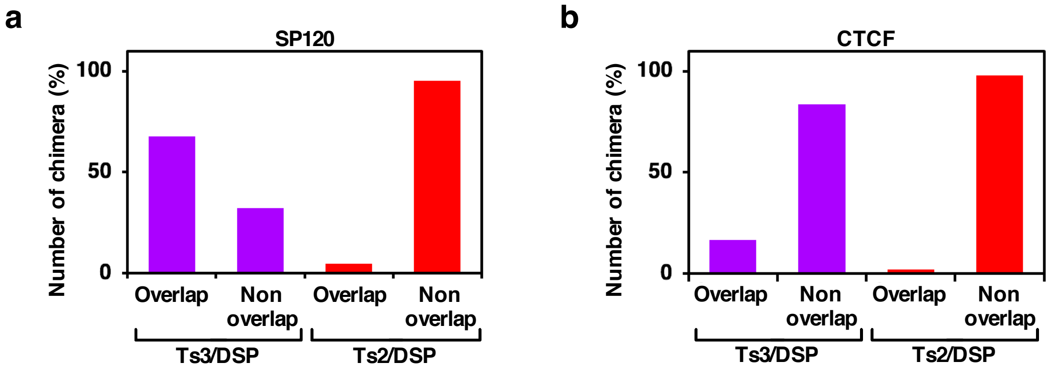


**Fig. S13** Coincidence of DSP sites with SP120 and CTCF binding sites that were determined by ChIP-seq. **a** Overlapping with SP120 sites. **b** Overlapping with CTCF sites that are taken from the rat liver ChIP-seq data [14].

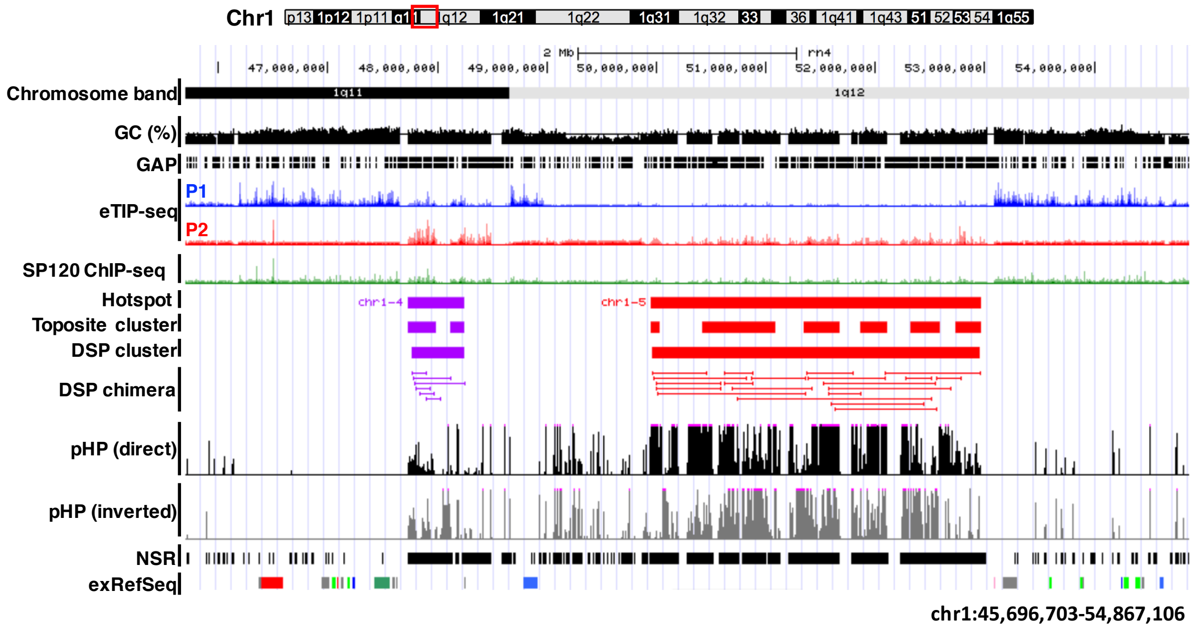


**Fig. S14** Correlation among the map positions of predicted homologous pairs (pHP) and other features. A representative region of rat chromosome 1 showing the UCSC browser view with custom tracks (hotspot, toposite cluster, DSP cluster, and DSP chimera). NSR (no signal region) designates genomic stretches that are unmappable. Hotspot, toposite cluster, DSP cluster and DSP chimera are depicted in magenta (Ts3-related) or in red (Ts2-related). The color code used for exRefSeq genes is the same as in Fig. 8c.

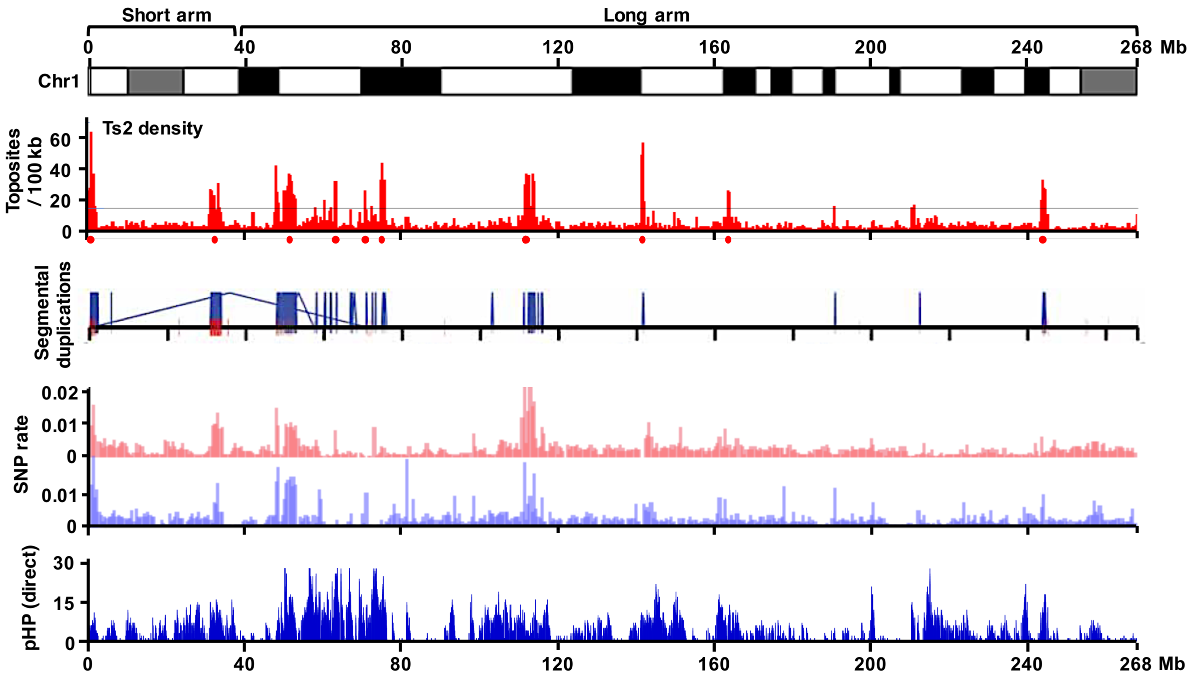


**Fig. S15** Remarkable correlation among Ts2 toposite, segmental duplication, and SNP rate. Entire chromosome 1 is shown. Ts2 toposite density was plotted as in Fig. 2a. Red dots indicate the position of Ts2-hotspots listed in Table S4. Intra-chromosomal duplications are designated by dark-blue tie lines and positions of inter-chromosomal duplications are indicated below by red bars. The SNP rate map for chromosome 1 was taken from the Rat Bac bowser of Riken. SNP rate maps for two rat strains are shown: F344/Stm (upper track in red) and LE/Stm (lower track in blue). Predicted homologous pair (pHP) was calculated as described in Methods. Only pHP for direct repeats is shown here.

**a**

- $\log_{10}$ (q-value)

| | Ts1 | Ts2 | Ts3 |
|---|---|---|---|
| Ts1 | 7 | 1* | 62* |
| Ts2 | | 6 | 0* |
| Ts3 | | | 120 |

* $N_{obs} < N_{exp}$

Ts3/Inter

**b**

Ts3/Inter on Sat I (N=617)

| | | Sat I subrepeats | | | |
|---|---|---|---|---|---|
| | | 1 | 2 | 3 | 4 |
| Sat I subrepeats | 1 | 76 | 53 | 11 | 69 |
| | 2 | | 74 | 92 | 21 |
| | 3 | | | 115 | 46 |
| | 4 | | | | 60 |

**c**

-$\log_{10}$ (q-value)

| | | Sat I subrepeats | | | |
|---|---|---|---|---|---|
| | | 1 | 2 | 3 | 4 |
| Sat I subrepeats | 1 | 4.2 | 1.0* | 14* | 0.3 |
| | 2 | | 2.7 | 0.0* | 5.2* |
| | 3 | | | 4.9 | 2.4* |
| | 4 | | | | 3.1 |

* $N_{obs} < N_{exp}$

**Fig. S16** Analysis of Sat I subrepeats linked by Ts3/inter chimera. **a** Statistical significance of toposite combinations for Ts3/Inter. Chi-squared test was conducted as in Fig. 3c between 'expected' and 'observed' frequencies of each combination. Resulting p-values were corrected by Benjamini-Hochberg method to obtain q-values. Shown in the matrix are minus-log transformed q-values. Asterisks indicate the combination in that observed number is smaller than expected one. The result clearly indicates that only Ts3-Ts3 combination is highly favored. **b** To reduce noisy background further, only Ts3/Inter chimeras that are clustered more than two within 100 kb span were selected (n=645). With respect to those overlapping with Sat I repeat (n=617), identity of subrepeats on chimera ends was determined by homology search and the combinatorial number is shown in the matrix. **c** Expected frequency was calculated from randomized combination of subrepeats and chi-squared test was done between the observed frequency. Statistical figures were obtained as in 'a'. Note that the homologous combinations (1-1, 2-2, 3-3, 4-4) are highly favored, the neighboring combinations (1-2, 2-3, 3-4, 4-1) are more or less neutral and the non-neighboring combinations (1-3, 2-4) are highly disfavored.
